# Discovery of lipid-mediated protein-protein interactions in living cells using metabolic labeling with photoactivatable clickable probes

**DOI:** 10.1101/2021.12.15.472799

**Authors:** Roman O. Fedoryshchak, Andrii Gorelik, Mengjie Shen, Maria M. Shchepinova, Inmaculada Pérez-Dorado, Edward W. Tate

## Abstract

Protein-protein interactions (PPIs) are essential and pervasive regulatory elements in cell biology. Despite development of a range of techniques to probe PPIs in living systems, there is a dearth of approaches to capture interactions driven by specific post-translational modifications (PTMs). Myristoylation is a lipid PTM added to more than 200 human proteins, where it may regulate membrane localization, stability or activity. Here we report design and synthesis of a panel of novel photocrosslinkable and clickable myristic acid analog probes, and their characterization as efficient substrates for human *N*-myristoyltransferases NMT1 and NMT2, both biochemically and through X-ray co-crystallography. We demonstrate metabolic incorporation of probes to label NMT substrates in cell culture and *in situ* intracellular photoactivation to form a covalent crosslink between modified proteins and their interactors, capturing a snapshot of interactions driven by the presence of the lipid PTM. Proteomic analyses revealed both known and multiple novel interactors of a series of myristoylated proteins, including ferroptosis suppressor protein FSP1 and spliceosome-associated RNA helicase DDX46. The concept exemplified by these probes offers an efficient approach for exploring the PTM-specific interactome, which may prove broadly applicable to other PTMs.

## Introduction

Protein-protein interactions (PPIs) are involved in virtually all aspects of cellular physiology, from signaling and protein trafficking, to protein turnover and enzymatic activity. Post-translational modifications (PTMs) are chemical changes to protein structure that often regulate PPIs. Identifying PPIs mediated specifically by PTMs in intact cells is challenging due to both the transient nature of PPIs and the fact that PTMs are not directly encoded in the amino acid sequence but rather are regulated dynamically by transferases or hydrolases.^1^ Whilst proximity labeling proteomics has emerged as a powerful tool for analysis of protein adjacency in cells,^2^ direct identification of PTM-dependent PPIs remains challenging. In-cell photocrosslinking, most often exploiting a photolabile diazirine coupled to affinity enrichment and shotgun proteomics^3^ has been highly successful in identification of ligand-protein interactions^4–7^ and metabolite-protein interactions, including lipid-protein interactions.^8,9^ More recently, diazirine-containing amino acid mimics (so-called photo-leucine, photo-isoleucine, photo-methionine^10,11^ and, most notably, photo-lysine^12^) have been used to identify PPIs in cells via photocrosslinking proteome-wide^13,14^ or encoded through codon reassignment in a protein of interest^15^. In one pioneering study, cells were metabolically labeled with a diazirine/alkyne dual-tagged palmitate probe, and photocrosslinking employed to detect multimerization of *S*-palmitoylated IFITM3.^16^

*N*-myristoylation is a lipid PTM whereby the 14-carbon saturated fatty acid myristate is transferred from myristoyl-Coenzyme A (**Myr-CoA**) to the N-terminal glycine of substrate proteins in cells, catalyzed by *N*-myristoyl transferases (NMT1 and NMT2 in humans). **Myr-CoA** is bound in a well-defined hydrophobic pocket, triggering binding of up to 10 residues of the N-terminus of a substrate protein in the substrate-binding groove.^17,18^ Myristate (**Myr**) is transferred to the substrate protein N-terminus followed by release of product myristoylated protein and CoA (Figure 1A), in a catalytic cycle recently described at high resolution.^19^ Myristoylation occurs both co-translationally and post-translationally in cells; it promotes dynamic association with cellular membranes which can be stabilized by additional signals such as further lipidation (often *S*-acylation), basic amino acids adjacent to the N-terminus or protein-protein interactions (PPIs), and reversed through further PTMs, soluble chaperone binding or an intramolecular change in conformation in a so-called myristoyl switch.^18,20^ There are only a few known examples of PPIs mediated directly by myristoylated N-termini; for example, myristate-mediated interaction of CHP1 to activate acyltransferase GPAT4,^21^ the well-defined myristate-binding pocket in chaperones UNC119A and UNC119B which recognize specific myristoylated clients for intracellular trafficking,^22,23^ and heme oxygenase 2, which can bind both free myristic acid and the myristoylated HIV-1 protein Gag.^24^

**Figure 1.**
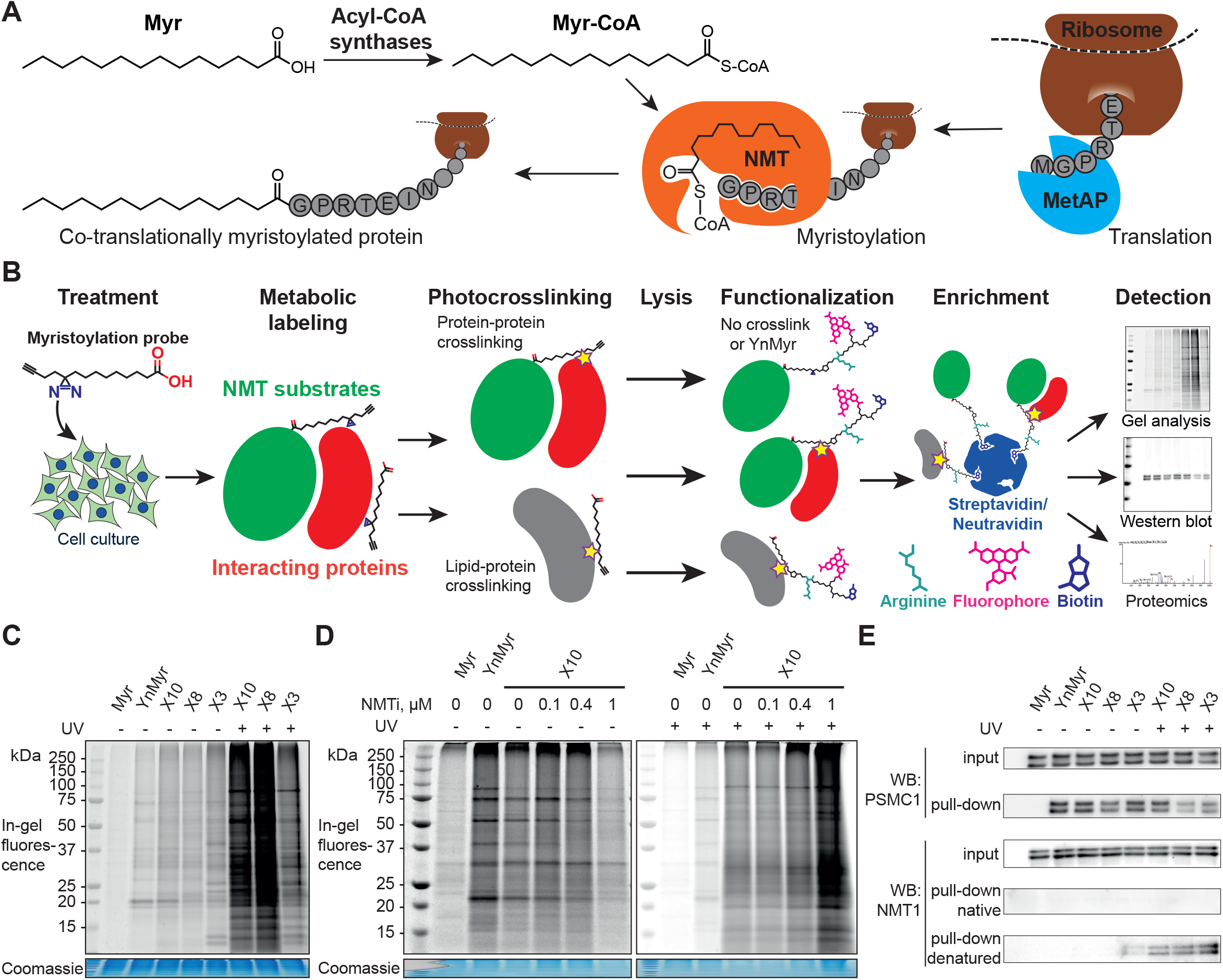
X3, X8 and X10 probes label myristoylated proteins in cells. **A**. Diagram showing cellular transformations leading to protein myristoylation. NMT acts following prior proteolysis to reveal an N-terminal glycine, either by initiator methionine aminopeptidase (MetAP) (co-translational myristoylation, shown) or by an endoprotease (post-translational myristoylation). **B**. Myristoylated proteome labeling strategy: diazirine probe is incubated with cells in culture and incorporated into the proteome, photocrosslinked by irradiation at 365 nm, and following lysis alkyne groups are functionalized with dye and/or biotin labels, optionally including a trypsin-digestible arginine linker. Proteins can be visualized by in-gel fluorescence or enriched on streptavidin/neutravidin beads for Western blotting and proteomics. **C**. Cells were treated with a panel of myristate analogs and visualized by in-gel fluorescence. A myristoylated protein labeling pattern is observed for all probes without UV-irradiation (UV) **YnMyr** (5 μM), **X3**/**X8**/**X10** (100 μM each), whilst **X3**/**X8**/**X10** probes are crosslinked to cellular proteins upon UV irradiation. **D. IMP-366** (NMTi) treatment inhibits myristoylated proteome labeling by **X10**; upon UV-irradiation, **X10** labeling is NMT-independent and is likely dominated by lipid-protein crosslinks. **E**. Cells were treated with a panel of myristate analogs, and labeled proteins enriched on streptavidin-conjugated beads. Western blots of NMT substrate PSMC1 before (input) and after enrichment (pull-down) indicate labeling with **X3**/**X8**/**X10** (100 μM each) probes comparable to **YnMyr** (5 μM). Equivalent blotting for NMT1 labeling (a buried site) shows photocrosslinking-dependent enrichment only with denaturation at 95 °C pre-CuAAC ligation.

Through structure guided design and optimization of novel myristate-based probes with dual diazirine and alkyne functionality, we report a technology platform to profile *N*-myristoylation-mediated PPIs in cells at the whole proteome level. Coupling these probes to enrichment and proteomic mass spectrometry, we identify multiple novel interactors for a series of myristoylated proteins, including ferroptosis suppressor protein FSP1 and spliceosome-associated RNA helicase DDX46. These data validate a novel tool for profiling myristoylation-mediated PPIs and exemplify a potentially generalizable approach for discovery of interactomes dependent on a specific PTMs.

## Results

### Design and synthesis of photocrosslinkable clickable myristate probes

Previously described photoactivatable myristate analogs are not suitable for metabolic incorporation into myristoylated proteome in human cells due to the incompatibility of these probes as human NMT substrates.^25,26^ We therefore elected to adopt a diazirine as a photoactivatable moiety, since this group is only slightly more sterically demanding than a methylene group and more likely to be accepted by the *N*-myristoylation machinery than bulkier alternatives. Upon UV irradiation, the diazirine group may eliminate nitrogen to generate a reactive carbene which readily crosslinks to neighboring molecules,^27,28^ and can be used for *de novo* mass-spectrometry identification of crosslinked proteins in combination with a bioorthogonal tag to enable click chemistry ligation and affinity purification.

We designed three photocrosslinkable clickable analogs of myristic acid, called **X3, X8** and **X10** based on the structure of alkyne-functionalized probe **YnMyr** that mimics myristic acid,^29^ and has previously been applied to identification of the *N*-myristoylated proteome and functional studies of NMT inhibition^30,31^ (**Myr**, Scheme 1A). The probes contain the diazirine moiety at specific positions along the 14-carbon fatty acid chain which were predicted to accommodate this group, judged by inspection of our previously reported structure of an HsNMT1:**Myr-CoA** complex (PDB: 4C2Y). Each of the analogs also carries an *ω*-alkyne group between carbons 13 and 14, similar to **YnMyr**, to enable functionalization via copper(I) catalyzed azide-alkyne cycloaddition (CuAAC) ligation (click chemistry) with various azide-bearing reagents.^30–32^ Syntheses of probes **X3, X8** and **X10** were designed such that the transformation of ketone to diazirine was left to the end of the synthesis (Scheme 1B-D). For **X3** synthesis, 10-bromodecanoic acid was reacted with trimethylsilylacetylene to yield trimethylsilyl-protected alkyne of lauric acid **2**,^33^ followed by condensation and homologation using the Masamune procedure^34^ to obtain the ketone precursor for **X3** (Scheme 1B). In contrast, the synthesis of ketone precursors **10** and **13** for probes **X8** (Scheme 1C) and **X10** (Scheme 1D) respectively, required a generally challenging sp^2^-sp^3^ C-C coupling; Pd(0) catalyzed coupling was performed by adding the respective chloroanhydrides to alkyl zinc iodides generated *in situ* from aliphatic alkyl iodides by Cu-Zn couple,^35^ resulting in desired ketone precursors of **X8** and **X10** in moderate yields. Each ketone precursor was treated with ammonia followed by hydroxylamine-*O-*sulfonic acid to obtain the corresponding diaziridine, which was oxidized to the diazirine using iodine in 20-42 % yields, similar to previously described protocols.^36^

### X3, X8, X10 probes metabolically label proteins in an NMT-dependent manner

The previously reported alkyne-bearing **YnMyr** probe is readily incorporated into the myristoylated proteome by incubation with cultured cells, and following lysis biotin- and/or fluorophore-functionalized capture reagents can be ligated to **YnMyr**-tagged proteins to enable subsequent in-gel visualization and/or affinity enrichment and proteomic analysis.^30,37,38^ A similar workflow was used in this study with the additional step of in-cell photocrosslinking for **X3**/**X8**/**X10** probes (Figure 1B).

To test the ability of diazirine containing probes to label the myristoylated proteome, each photocrosslinkable probe **X3**/**X8**/**X10** at 100 μM, or **YnMyr** at 5 μM concentration (yielding equal labeling intensity), or negative control myristic acid **Myr**, were added to HeLa cells in culture and incubated for 24 h to allow incorporation, followed by *in situ* irradiation with UV light at 365 nm (+UV; 5 min) to initiate in-cell diazirine photoactivation, or without irradiation (-UV) in control samples. CuAAC ligation of alkyne-tagged proteins to azido-TAMRA (AzT)^32^ was used for in-gel fluorescence visualization (Figure 1C). Without UV-irradiation (-UV), protein labeling for **X3**/**X8**/**X10** resembled **YnMyr**, suggesting similar proteins are labeled independently of the presence of diazirine group. Interestingly, **X3** showed some additional bands even without UV irradiation, suggesting additional reactivity or instability when the diazirine is located β- to the carbonyl. Upon UV-irradiation (+UV), the number and intensity of fluorescent bands was greatly increased, which we attributed to lipid-protein crosslinking whereby protein-bound **X3**/**X8**/**X10** fatty acids form crosslinks that greatly increase the range of proteins visualized in-gel (Figures 1C, 1D). To confirm the dependence of labeling on NMT activity, we examined the impact of a potent and selective NMT inhibitor (NMTi) **IMP-366**^39,40^ on labeling by a representative probe (**X10**); with increasing NMTi concentration the majority of fluorescently labeled bands disappeared, indicating a decrease in NMT-dependent metabolic labeling of myristoylated proteins, similarly to the previously published data.^30,37^ The lack of an evident NMTi-mediated decrease in labeling under UV-irradiation is likely due to masking of the signal from myristoylated proteins by a much more intense lipid-protein crosslinking background (Figure 1D). To confirm labeling of a specific NMT substrate, we ligated proteins tagged with **X3, X8** or **X10** to an azido-TAMRA-biotin (AzTB) capture reagent followed by streptavidin-conjugated magnetic bead enrichment to evaluate metabolic labeling and capture of PSMC1, a previously identified myristoylated protein and a component of the proteasome complex.^30^ Enrichment of PSMC1 was observed with all alkynylated probes with or without UV irradiation, but no enrichment in the absence of alkyne in the myristic acid control experiment was observed (Figure 1E); however, higher molecular weight bands on the blot signifying protein-protein crosslinking in the UV-irradiated samples were not observed for PSMC1 (data not shown).

### X3/X8/X10 probes form stable crosslinks with NMT in cells upon UV irradiation

As a substrate in the myristoylation reaction, **Myr-CoA** interacts with NMT non-covalently, and therefore NMT1 and NMT2 are not labeled in experiments with **YnMyr** enrichment. However, irradiation of **X3**/**X8**/**X10**-treated samples induced crosslinks between NMT1 and metabolically modified **X3**/**X8**/**X10**-CoA probes, which could be detected by western blotting (Figure 1E). Consistent with our expectations, without denaturation of proteins in the cell lysate prior to CuAAC, the alkyne functionality appears to be buried inside NMT1 and is inaccessible for ligation to azido-biotin. Upon heating (95 °C) the lysates in the presence of 1% sodium dodecyl sulfate (SDS), it was possible to ligate and enrich NMT1 photocrosslinked to each probe; again, probe **X3** showed evidence of background crosslinking even without UV irradiation.

### X3/X8/X10 label NMT substrates proteome-wide

Using **YnMyr** as a reference point for myristoylated proteome labeling,^30^ we assessed proteome-wide labeling with each of the **X3**/**X8**/**X10** probes using neutravidin enrichment and quantitative SILAC proteomics. An equal mixture of lysates from four HeLa cell cultures treated with each probe (**YnMyr, X3, X8** and **X10**) grown in heavy arginine and lysine (R10K8) -containing media was used as spike-in standard to normalize quantification across experiments. Cultures grown in light media (R0K0) were treated with each myristate analog and with either 0.5 μM NMT inhibitor **IMP-366**^39^ or DMSO vehicle, and heavy lysate spike-in added to each lysate to enable quantitative analysis of proteome-wide NMT-dependent probe incorporation (Figure 2A, Supplementary Table S1). Ligation to an azido-arginine-biotin capture reagent and enrichment for probe-tagged proteins on neutravidin-agarose beads demonstrated that the labeling of myristoylated proteins with **X3**/**X8**/**X10** or **YnMyr** decreased on NMTi treatment. Quantitative comparisons with/without NMTi show a Pearson correlation >0.9 between the samples treated with **YnMyr** *vs* **X10** or **X8**, and 0.84 *vs* **X3** (Figure 2B), suggesting these diazirine derivatives label myristoylated proteins equivalently to **YnMyr** in cells.

**Figure 2.**
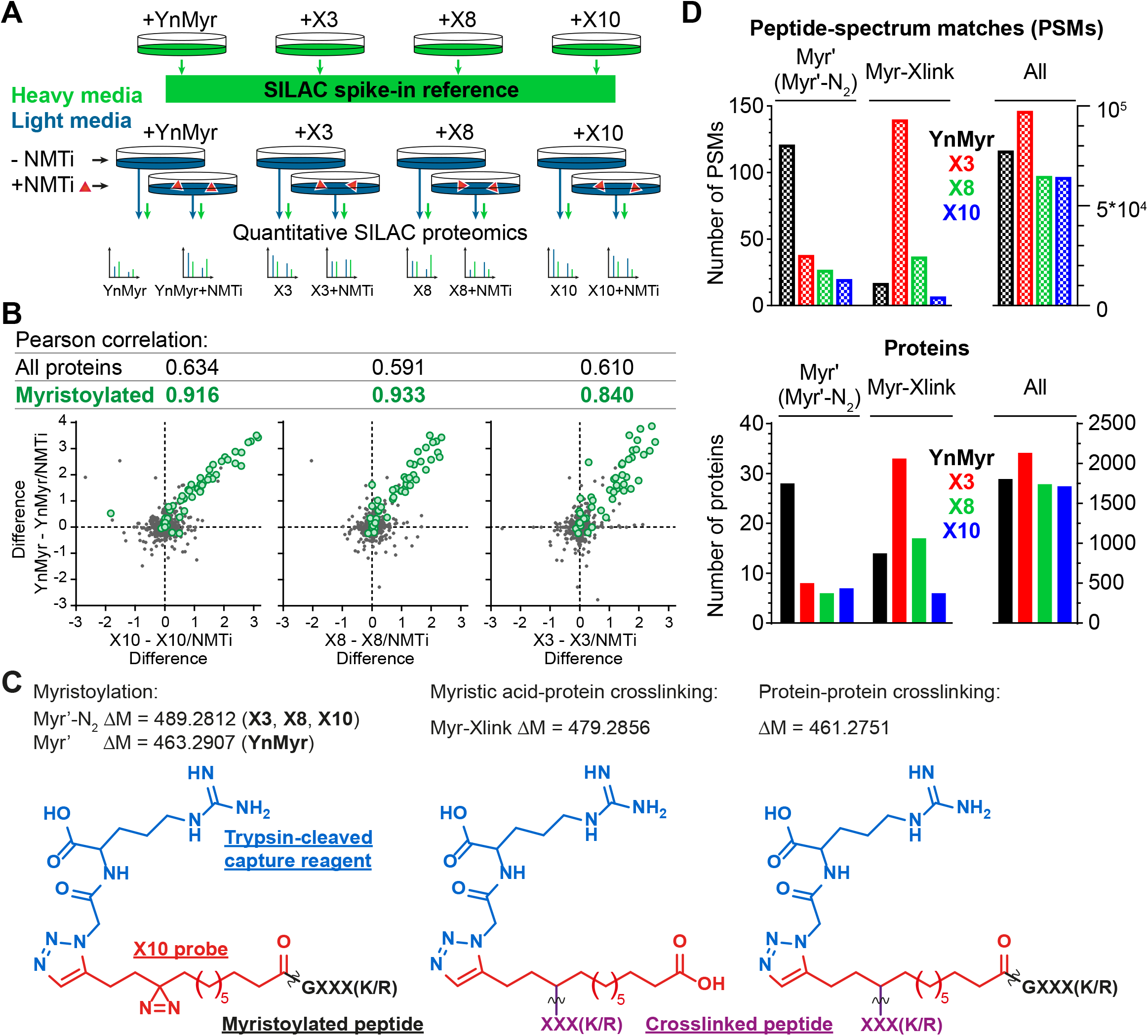
Incorporation of photocrosslinkable clickable probes into the myristoylated proteome. **A**. Experimental design for quantitative SILAC proteomics validating myristoylated proteome labeling by **X3**/**X8**/**X10** (100 μM each) compared to positive control **YnMyr** (5 μM) and negative controls obtained by 0.5 μM NMTi treatment. Treated cells were lysed, ligated to azido-arginine-biotin capture reagent and enriched on neutravidin-agarose resin. **B**. Pearson correlation between myristoylated protein enrichment (green) using either photocrosslinkable clickable probes **X3**/**X8**/**X10** or **YnMyr**. Grey - background (non-myristoylated) proteins. Axes are differences in protein enrichment between vehicle (DMSO) and NMTi-treated samples for **X3**/**X8**/**X10** (X-axes) and **YnMyr** (Y-axis). **C**. Structures and masses of PTMs searched to identify **X3**/**X8**/**X10**/**YnMyr**-related modifications. **D**. Left – total number of Myr’, Myr’-N2, Myr-Xlink (see Figure 2C) modified peptide-spectrum matches (PSMs, top) or proteins (bottom) identified using PEAKS search engine^41^ for **X3**/**X8**/**X10** or **YnMyr**. Right – all PSMs or proteins detected in the experiment for a given probe.

We used the PEAKS *de novo* sequencing analysis package^41^ to identify probe-modified peptides in MS/MS spectra, denoting Myr’ for **YnMyr**-labelled or Myr’-N2 for **X3**/**X8**/**X10**-labelled N-terminal peptides, respectively; and Myr-Xlink for **X3**/**X8**/**X10** probe-to-protein crosslinks (Figure 2C). We identified 121 peptide spectrum matches (PSMs) with Myr’ modification corresponding to 28 known myristoylated proteins. Probes **X3, X8** and **X10** were less efficient in direct detection of lipidation, although Myr’-N2 modifications for each diazirine-containing probe were detected on 6-8 proteins, with multiple PSMs per identification (Figure 2D).

### Identification of X3/X8/X10-direct-to-protein crosslinks

Intriguingly, several non-myristoylated proteins were enriched in the **X3**/**X8**/**X10** samples compared to **YnMyr** (Figures S1A, S1B, S1C). **X8** and **X10** probes consistently enriched for components of the fatty acid beta-oxidation pathway, such as long-chain-fatty-acid-CoA ligases (ACSL3, ACSL4), acyl-CoA dehydrogenases (ACAD9, ACADM), enoyl-CoA hydratase (ECHS1), 3-hydroxyacyl-CoA dehydrogenase (HSD17B4) and 3-ketoacyl-CoA thiolases (ACAA1, ACAA2). These proteins are involved in fatty acid and acyl-CoA metabolism, and apparent enrichment may result in part from changes in gene expression in response to treating with a relatively high concentration of fatty acid probe (100 μM **X8**/**X10** vs 5 μM **YnMyr**), or from metabolic activation in the absence of UV light.

Numerous additional proteins were enriched in **X3**-labeled samples, and we considered the possibility that some of these proteins may be crosslinked to **X3** non-specifically (without UV irradiation). We searched for PSMs corresponding to Myr-Xlink peptides (Figure 2C) on all residues and found 140 PSMs for 33 proteins with Myr-Xlink modification in addition to 35 PSMs for 8 proteins corresponding to Myr’-N2 modification (Figure 2D). Myr-Xlink identifications in **YnMyr**-treated samples served as false-positive controls, identifying 16 PSMs for 14 proteins. Similarly low numbers of Myr’-Xlink PSMs were observed in non-irradiated **X8**- and **X10**-treated samples. Taken together, these data indicate a degree of baseline reactivity for the diazirine group in β-position to the carbonyl and is consistent with the crosslinking ‘leakage’ of **X3** observed under non-irradiated conditions by Western blot (Figure 1E). Cytosolic acetyl-CoA acetyltransferase (ACAT2), a protein involved in lipid metabolism, is among the proteins modified by **X3** and quantitatively enriched compared to **YnMyr** (Figure S1A). We found 14 PSMs linking **X3** to ACAT2 with modifications on four residues (P236, Y237, G242, T245) located close to the CoA binding site (Figure 3A, PDB:1WL4^42^). Similarly, heme oxygenase 2 (HMOX2), also known as a myristate-binding protein,^24^ crosslinked with **X3** at two distinct sites (A48 and G49, 10 Myr-Xlink PSMs) which map to the myristate binding site in the published myristic acid-HMOX2 crystal structure (Figure 3B, PDB:5UC9^24^). Heme oxygenase 1 (HMOX1) is structurally homologous to HMOX2, and we identified 13 Myr-Xlink PSMs for **X3** on residues (H25, T26, E29) adjacent to the HMOX1 heme-binding site (Figure 3C, PDB:1N45^43^). This novel HMOX1 myristate binding site is found at the canonical heme site, closely analogous to HMOX2 despite HMOX1 and HMOX2 sequence divergence. Although not the main focus of the present study, these data highlight the potential of these probes for mapping known and novel protein/lipid or protein/acyl-CoA binding sites.

**Figure 3.**
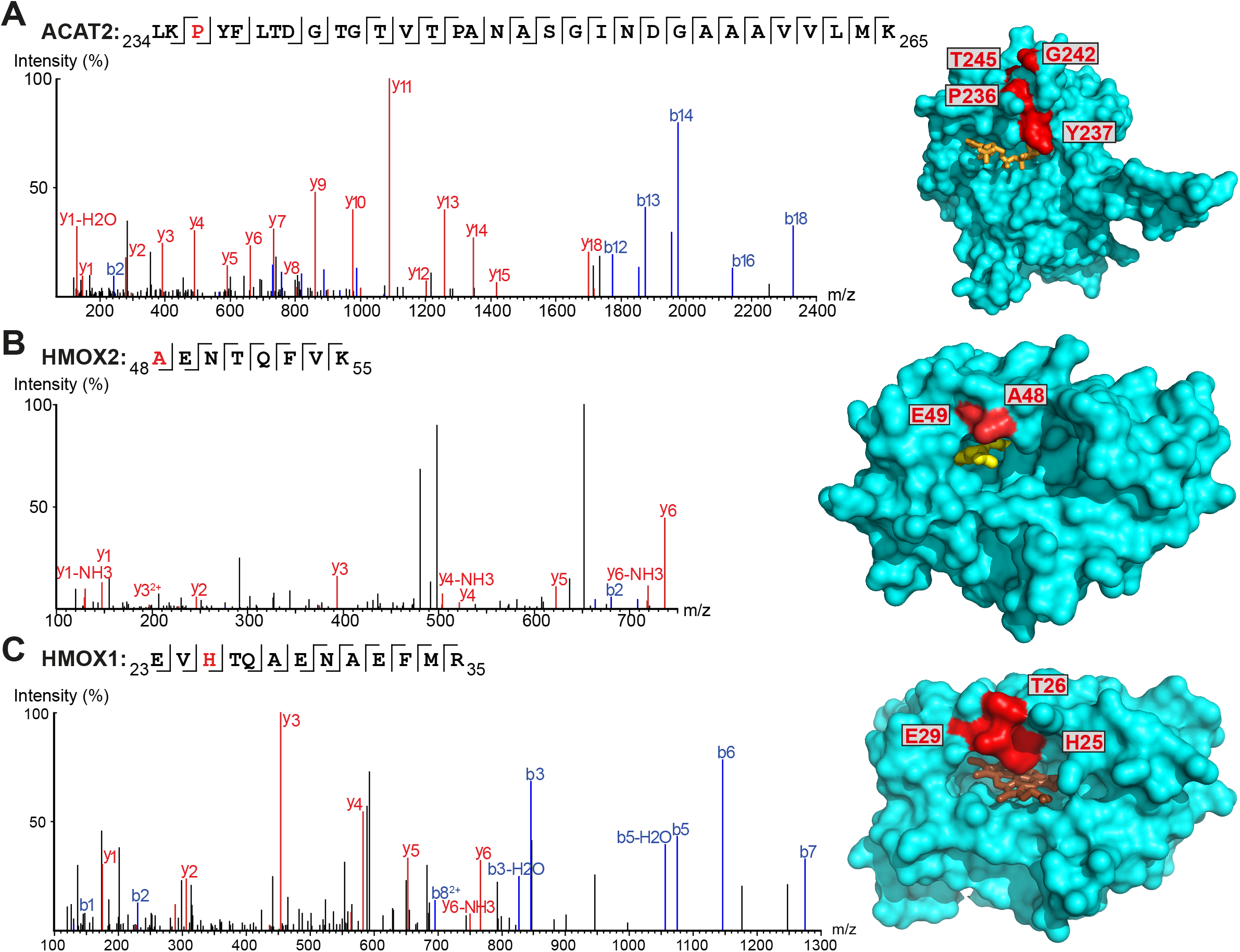
Myristate probes X3/X8/X10 form crosslinks with myristate-binding proteins. **A-C**. Representative MS/MS fragmentation spectra and structural positions of crosslinked amino acids (red) for ACAT2 (**A**, PDB:1WL4, orange – acetyl-CoA), HMOX2 (**B**, PDB:5UC9, yellow – myristic acid) and HMOX1 (**C**, PDB:1N45, brown - heme) structures.

### Myristoylated proteins labeled with X3 and X10 probes form photocrosslinks with interactors

To explore whether probe crosslinking could capture PPIs specific to *N*-myristoylated proteins in addition to binary protein-lipid interactions, we analyzed the impact of NMTi on crosslinking to identify non-myristoylated proteins captured by virtue of NMT-mediated probe incorporation (Supplementary Table S2), focusing on UV-irradiated **X3**- or **X10**-treated cells with or without NMTi. SILAC proteomic analysis identified PSMs that recapitulated crosslinking to ACAT2 (residues P236, G242 and G244) and HMOX2 (A48, G49) in **X3** samples, among other proteins, although PSMs for peptides crosslinked to **X10** were not identified, highlighting differences between these probes (see Discussion). As before, known myristoylated proteins were identified in both **X3**- (Figure S2A) and **X10**- (Figure S2B) treated samples, with their enrichment significantly reduced upon NMTi treatment. In addition, we observed a set of non-myristoylated proteins (Figure S2C) enriched specifically in non-NMTi samples, representing candidates for myristoylation-dependent PPIs. The number of significantly coenriched proteins was greatly increased in UV-treated samples in comparison to the previous analysis without UV-irradiation (Figures S2D and S2E). Identification of peptide-to-peptide crosslinks by MS/MS proved challenging, however, and we were unable to confidently resolve direct interactions between myristoylated proteins and enriched non-myristoylated proteins at the whole proteome level. Therefore, we turned to a targeted approach to identify PTM-dependent interactions of specific myristoylated proteins.

### X10-CoA is an efficient human NMT substrate *in vitro* and is accommodated in the Myr-CoA pocket

We next sought to confirm **X10** as an optimal photoactivatable NMT substrate biochemically and structurally, by exploring the binding mode of its activated CoA thioester form (**X10-CoA**) by X-ray crystallography. Thioester **X10-CoA** (Scheme 1E) was generated by conjugation of **X10** and Coenzyme A thiol (CoA-SH) in the presence of 1,1’-carbonyldiimidazole (CDI) and base.^29^ Enzyme kinetics were analyzed using an *in vitro* assay which detects CoA-SH release through a fluorogenic reaction with 7-diethylamino-3-(4-maleimidophenyl)-4-methylcoumarin (CPM),^44^ comparing the activity of HsNMT1 and HsNMT2 towards transferring the fatty acid moiety of **X10-CoA** and the native substrate **Myr-CoA** to the model peptide based on c-Src N-terminal peptide (16 μM H-GSNKSKPK-NH^2^). Catalytic efficiency of **X10-CoA** and **Myr-CoA** was very similar for both enzymes, NMT1 and NMT2, with a small change in K_M_ (Figures 4A, S3A), indicating excellent biochemical compatibility between **X10** and human NMT1/2 *in vitro*.

**Figure 4.**
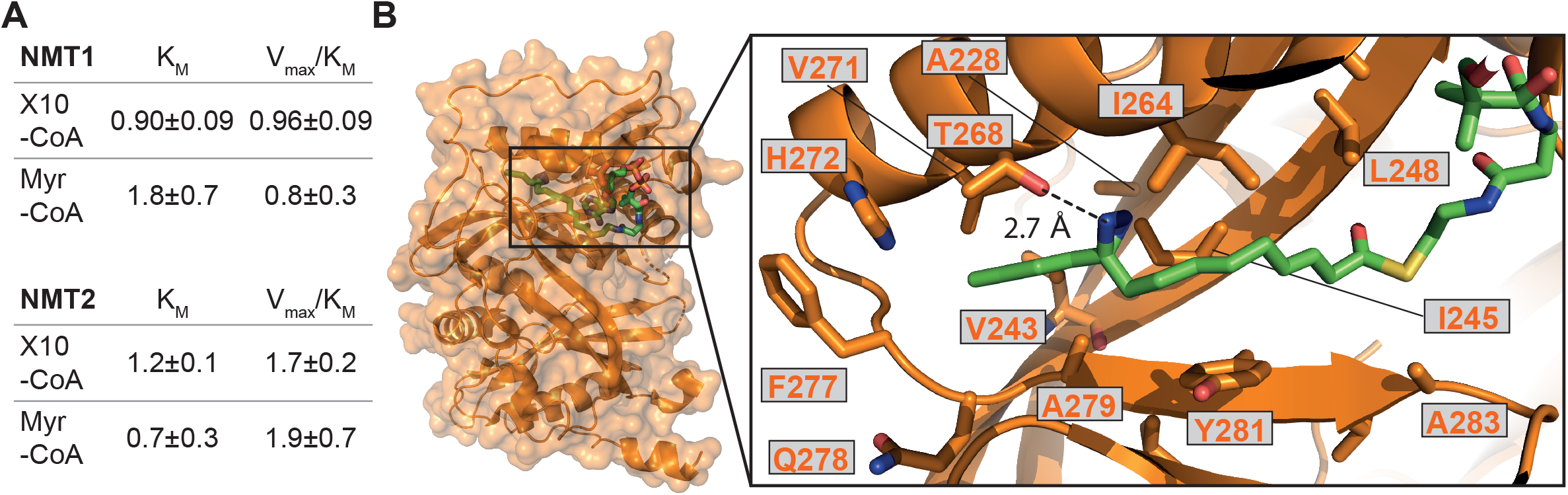
X10-CoA is an NMT1/2 substrate. **A**. Rates of enzymatic transfer of **Myr** or **X10** by HsNMT1 or HsNMT2 to a synthetic peptide measured *in vitro* by CPM assay;^44^ both natural and artificial substrate show similar catalytic efficiencies (V_max_/K_M_±SD, RFU*min^-1^*μM^-1^*10^6^) and Michaelis constants (K_M_±SD, μM); **B**. Crystal structure of HsNMT1 (orange) in complex with **X10-CoA** (green) showing overall structure and an enlarged view of the **X10** moiety (chain A) with a potential hydrogen bond between the diazirine and T268. Color code: blue (nitrogen), red (oxygen), yellow (sulfur), and orange (protein structure).

To gain insights into the binding mode of **X10-CoA** in the active site of NMT, we obtained crystals of HsNMT1 in complex with **X10-CoA** which enabled us to solve the structure at 2.37 Å resolution (Figures 4B, S3B-D, Supplementary Table S3). The two HsNMT1:**X10-CoA** complexes per asymmetric unit exhibited the same fold (root mean-square deviation (RMSD) of 0.497 Å for 354 Cα atoms), whilst structural superimposition of this complex with our previously reported HsNMT1:**Myr-CoA** structure (PDB: 4C2Y^30^) confirms an equivalent HsNMT1 fold in complexes with **X10-CoA** or the natural substrate (RMSD 0.587 Å over 355 Cα atoms, Figure S3B). Both **X10-CoA** molecules were well-defined in the electron density, which enabled us to model **X10-CoA** and its diazirine group unambiguously. Consistent with the capacity of **X10-CoA** to mimic the natural substrate, **X10-CoA** binds at the **Myr-CoA** binding site with the **X10** moiety lying along the myristate-binding groove in a very similar conformation to the **Myr-CoA** myristate (Figure S3C). The myristate binding site is highly hydrophobic except for the presence of T268 and H272 residues, and interestingly the diazirine is situated at hydrogen bonding distance from the T268 side-chain hydroxyl (Figure 4B).

### X10 probe enables *de novo* identification of PTM-dependent protein-protein interactions in live cells

We chose to focus on **X10** for targeted identification of myristoylation-dependent PPIs, as among the three probes it shows the highest efficiency and fidelity for myristoylation, with optimal NMT substrate properties and a well-defined binding mode (see above). HEK293 cells were transfected with various myristoylated proteins engineered to bear a C-terminal Twin-Strep-tag (TST) for affinity purification, or mCitrine-TST as a non-myristoylated negative control (for the full list of the 20 protein constructs used, and related proteomics data, see Supplementary Table S4). Upon transfection, cells were treated with 100 μM **X10** for 24 h before UV-irradiation. Cells were lysed with a buffer containing 1% SDS and heated at 95 °C for 10 min to eliminate non-covalent PPIs, after which myristoylated TST-tagged proteins were enriched (at ambient temperature) with Strep-Tactin resin, digested with trypsin and analyzed by proteomics (Figure 5A). Label-free quantification was used to determine the fold change of proteins enriched by a given myristoylated protein construct against the combined data of the 19 other protein constructs in the overall experiment. As expected, the corresponding TST-tagged constructs were the most strongly enriched proteins in each case, and no proteins were significantly enriched in the mCitrine control, demonstrating the specificity of TST enrichment and the dependence of photocrosslinking on the incorporation of **X10** (Figure 5B). However, several myristoylated protein constructs enriched putative photocrosslinked interactors (Supplementary Table S4), with DDX46 (Figure 5C) and FSP1 (Figure 5D) showing the highest hit rates.

**Figure 5.**
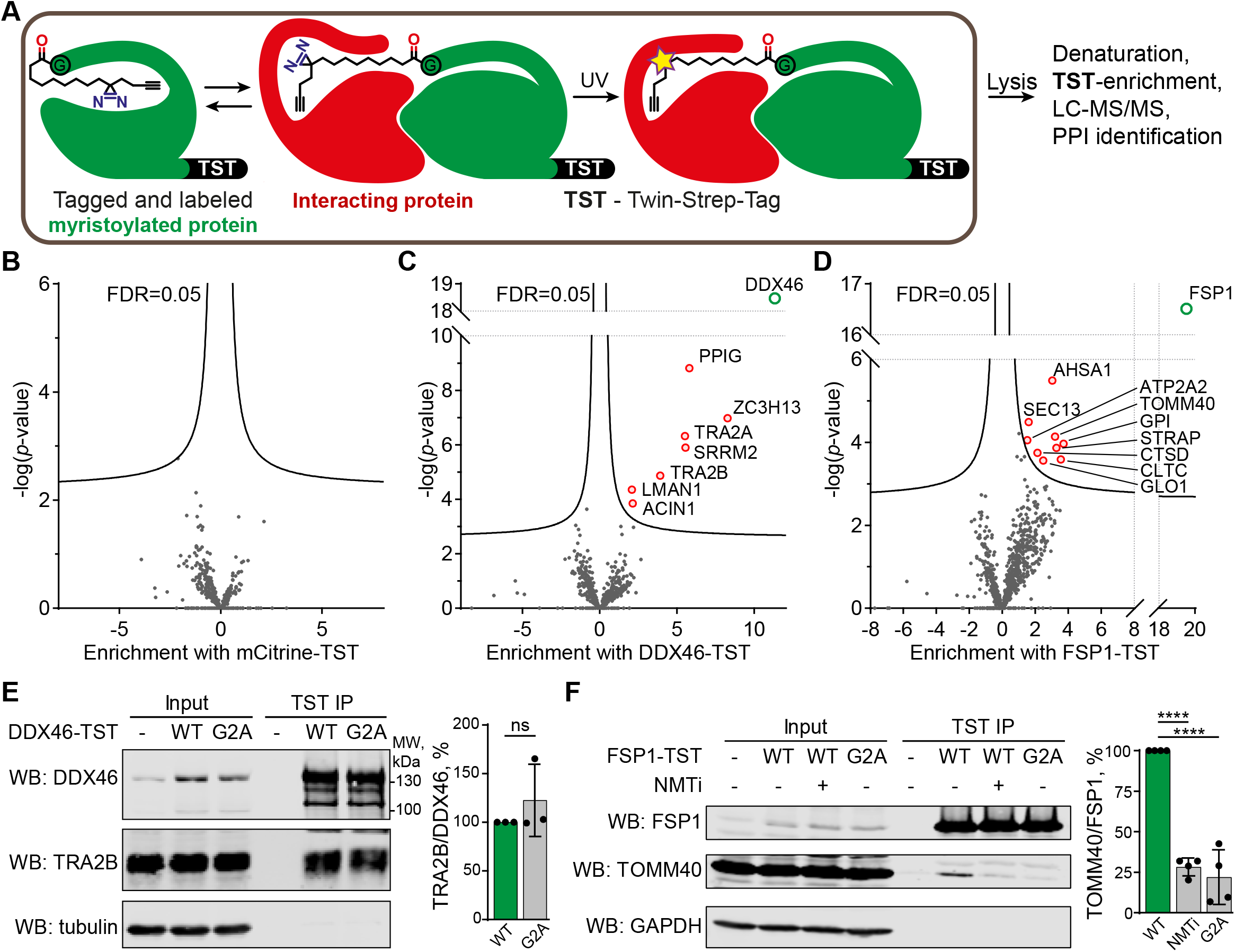
Identification of PPIs by photocrosslinking. **A**. Scheme of **X10** photocrosslinking experiment to identify PPIs of TST-tagged myristoylated proteins. **B-D**. Fold protein enrichment from cells transfected with mCitrine-TST (**B**), DDX46-TST (**C**) or FSP1-TST (**D**) in triplicate, relative to all other samples combined (19 proteins x 3 replicates) on X-axis, plotted against statistical significance (Y-axis). **E-F**. Validation of DDX46 (**E**) and FSP1 (**F**) interactors. Pulldowns for DDX46-TST and FSP1-TST were probed with TRA2B and TOMM40 antibodies, respectively. α-tubulin or GAPDH were used as loading controls. Normalization was performed by dividing TRA2B or TOMM40 antibody signal by DDX46 or FSP1 signal, respectively. Data are shown as mean ± s.e.m., n = 3-4. ****p = 0.0001, ns – no significant difference, calculated by Student’s t-test (two-tailed, unpaired).

Myristoylated ATP-dependent RNA helicase DDX46 (homologue of *Saccharomyces cerevisiae* PRP5) is localized in the nucleus and has essential role in splicing, where it participates in the formation of the 17S U2 snRNP complex, a subunit of the spliceosome A and E complexes.^45^ Consistent with DDX46 localization and function, proteins ACIN1, PPIG, SRRM2, TRA2A/B, ZC3H13 identified in our data as DDX46 interactors are all components of RNA splicing and have high confidence as interactors based on previously reported interactome data^46^ (Figure S4). We further analyzed the interaction of DDX46 with TRA2B, an RNA-binding protein involved in control of pre-mRNA splicing.^47,48^ DDX46-TST or its myristoylation-deficient G2A mutant were transfected and immunoprecipitated from HEK293 cells on Strep-Tactin resin, showing that both wild type (WT) and G2A mutant interacted with TRA2B to a similar extent (Figure 5E). These data indicate that the DDX46-TRA2B interaction is mediated by structural and sequence determinants other than myristoylation, but nevertheless demonstrate the utility of **X10** as an enzymatically incorporated photoaffinity probe for identification of novel interactors in intact cells.

FSP1 (ferroptosis suppressor protein 1) is a myristoylated oxidoreductase that converts Coenzyme Q_10_ into its reduced form at the membrane, preventing lipid oxidation by quenching reactive oxygen species, and suppressing ferroptosis, a type of iron-dependent cell death.^49,50^ We identified several FSP1-interacting proteins associated with mitochondria and oxidative stress, including peroxisomal oxidase GLO1, oxidative stress-associated ATPase ATP2A2 and protease cathepsin D among others. We used a similar co-immunoprecipitation approach to validate the interaction of FSP1-TST with TOMM40, which is a mitochondrial protein that facilitates protein import into mitochondria,^51,52^ and is itself also myristoylated. TOMM40 co-immunoprecipitated with FSP1, which was strongly suppressed in a myristoylation-deficient FSP1 G2A mutant (to 22% of WT FSP1), whilst NMT inhibition in the presence of wildtype FSP1 similarly reduced immunoprecipitation to 28% of WT (Figure 5F). These data support a novel interaction between two mitochondrial myristoylated proteins FSP1 and TOMM40, for which FSP1 myristoylation is essential.

## Discussion

Myristoylation is an essential cellular process, inhibition of which has been demonstrated as a potential treatment in a variety of infectious diseases^37,53^ and cancer.^54,55^ Besides providing target proteins with lipophilicity, myristoylation can also lead to changes in protein conformation and thus affect PPIs. However, only a few myristate-mediated PPIs have been described. We sought to identify weak and transient myristate-mediated PPIs with minimal perturbation in intact cells using *in situ* photocrosslinking through diazirine- and alkyne-modified myristate probes **X3, X8** and **X10**.

Probes **X3**/**X8**/**X10** are diazirine modified derivatives of the clickable myristate analog **YnMyr**, which was previously applied in cells to characterize myristoylated proteome and study the effects of NMT inhibition. Our data show that these probes mimic myristate in cells through metabolic incorporation into NMT substrates, a feature we attribute to the small size and minimal perturbation caused by diazirine and alkyne modification. Like **YnMyr**, these probes must first be activated as Coenzyme A derivatives, likely through long-chain-fatty-acid-CoA ligases 3 and 4 (ACSL3/4) that we found crosslinked with **X3**/**X8**/**X10** (Figure S2). Interestingly, 100 μM **X3**/**X8**/**X10** was required to reach the same level of myristoylated protein labeling achieved with 5 μM **YnMyr**, which we hypothesize is either due to more efficient catabolism of diazirine-containing probes or to a lower efficiency of uptake or activation by acyl-CoA ligases. **X3**/**X8**/**X10** are nevertheless recognized as substrates by native NMTs in cells, while in enzyme activity assays **X10-CoA** shows equal catalytic efficiency with HsNMT1 and HsNMT2 to the native substrate **Myr** and binds in the **Myr-CoA** binding site of HsNMT1 as determined by X-ray crystallography. Furthermore, we find that myristoylated proteins labelled with probes in cells can participate in PPIs and form crosslinks to respective interactors.

Myristoylation presents a convenient PTM system to apply photocrosslinking proteomics approaches to selected proteins and offers some complementary advantages over alternative approaches. Myristoylated proteins are typically stoichiometrically labeled at the point of synthesis, with no mechanism for subsequent removal, in contrast to reversible *S*-acylation, whilst photo-amino acid labeling requires either laborious and disruptive introduction of amber codon suppression or indiscriminate whole-proteome photo-amino acid labeling by photo-substitutes of natural amino acids. On the other hand, metabolic incorporation of **X3**/**X8**/**X10** into the myristoylated proteome may result in labeling of >100 natively myristoylated human proteins, and the resulting photocrosslinks may be due to interaction of one or several myristoylated proteins with an interactor. It may be possible to deconvolute such interactions in the future at the whole proteome level using identification of photocrosslinked peptide sequences from MS/MS spectra. Identification of such crosslinks proteome-wide is currently only possible using specialized chemical crosslinking reagents,^56^ and would likely require advances in mass spectrometry hardware (quality of spectra) and software (searching multi-dimensional MS/MS space) to be used with our current reagents. Another complication of using metabolic labeling to study *N*-myristoylation with myristic acid analogs is their propensity to metabolize and be incorporated into other lipid species. However, we took advantage of this property of **X3**/**X8**/**X10** probes to identify numerous lipid-protein interactions proteome-wide, in line with previous research,^57^ alongside precise sites of fatty acid-protein binding through crosslinking analysis by *de novo* sequencing;^58^ in contrast, standard database searches (e.g. in MaxQuant^59^) failed to identify crosslinks using the same search parameters.

When analyzing fatty acid-protein crosslinks, we noticed an increased baseline reactivity of diazirine group in β-position to carboxyl group (Figures 1E, 2D, S1A). To our knowledge, such a juxtaposition of functional groups has been reported only once previously, and was not applied for protein photocrosslinking,^60^ with the majority of recent reports placing a diazirine group γ- to a carboxyl/amide group, including so-called “minimalist photocrosslinkers”,^61^ or aromatic trifluoromethyl diazirines. Our observation that β-diazirine carboxylic acid **X3** undergoes spontaneous crosslinking may be explained by diazirine conversion to diazoalkane by protonation,^62^ providing a mechanism by which **X3** is more susceptible to non-irradiative decomposition: the weakly acidic proton of the neighboring carboxyl is ideally placed to hydrogen bond with a diazirine lone pair and thus facilitates transition to a reactive *N*-protonated carbocationic diazolakane species. Diazoalkanes may also be converted to reactive carbenes upon 365 nm UV-irradiation.

Comparing the **X3**/**X8**/**X10** probes, we show that **X10** offers improved and consistent myristoylated proteome labeling on par with **YnMyr**. Using **X10**, we were able to capture and identify myristate PTM-mediated interactions by transfection and affinity-tag enrichment of a protein of interest, whereby the diazirine functionality of **X10** was used for crosslinking but the alkyne was left unreacted. Interactions were identified and validated for myristoylated FSP1 and DDX46 in addition to known interactions, for example, for myristoylated Src kinase with KHDRBS1^63^ and HNRNPK^64^ (Supplementary table 3). Whilst the majority of DDX46 interactors identified in the present study are RNA splicing-associated proteins in line with its role as a component of the U2 snRNP complex, our data suggest a potential wider role for DDX46 in RNA splicing. For example, we have confirmed an interaction of DDX46 with pre-mRNA-binding protein TRA2B which regulates alternative splicing, and these findings may inform investigation of functional roles for DDX46 interaction with TRA2B as well as TRA2A, ACIN1, PPIG, SRRM2 and ZC3H13 in future studies of splicing regulation. FSP1 was recently identified as a novel ferroptosis suppressor protein and potential target for cancer therapy;^49,50^ however, understanding of the FSP1 interaction network is currently limited. Here, we have identified nine probable FSP1-interacting proteins, and confirmed a physical interaction with TOMM40. FSP1 myristoylation appears necessary for the FSP1-TOMM40 interaction, although it remains to be established whether the interaction is direct or through co-localization. Further investigation will be required to dissect the role of FSP1-TOMM40 interaction in normal cell state and ferroptosis.

## Supporting information

NMR spectra

## Associated content

Supporting Information. Materials and Methods, Supplementary Table S3 and Figures.pdf

Supplementary Table S1. Proteomics validation of metabolic labeling by X3, X8, X10.xlsx

Supplementary Table S2. X3 and X10 photocrosslinking proteomics.xlsx

Supplementary Table S4. Crosslinked interactors proteomics.xlsx

Supporting Information. 1H NMR spectra.pdf

RCSB PDB accession: 5NPQ.

The mass spectrometry proteomics data have been deposited to the ProteomeXchange Consortium via the PRIDE partner repository with the dataset identifiers PXD027239, PXD027394, PXD029944.

## Author information

### Notes

EWT is a founder, shareholder and Director of Myricx Pharma Ltd.

### Author Contributions

ROF and EWT conceived the project. ROF performed all experiments except the following. AG performed validation co-immunoprecipitation and Western blot experiments. MS and IPD crystallized HsNMT1:X10-CoA. MMS characterized spectroscopic data. EWT obtained funding and supervised the research. ROF and EWT wrote the manuscript with input from all authors.

## Acknowledgements

The authors thank Dr Antonio Konitsiotis for plasmids, invaluable support and critical reading of the manuscript, Dr Remigiusz Serwa and Dr Julia Morales-Sanfrutos for help with setting up proteomics experiments. This project was funded from the European Union’s Seventh Framework Programme for research, technological development and demonstration under grant agreement no. 607466. Work in the Tate laboratory was supported by Cancer Research UK (C29637/A20183), by the Biotechnology and Biological Sciences Research Council (BBSRC) grants BB/S001565/1 and BB/N016947/1, and by the Francis Crick Institute which receives its core funding from Cancer Research UK (FC001097, FC010636), the UK Medical Research Council (FC001097, FC010636), and the Wellcome Trust (FC001097, FC010636).

## Figure legends

**Scheme 1.**
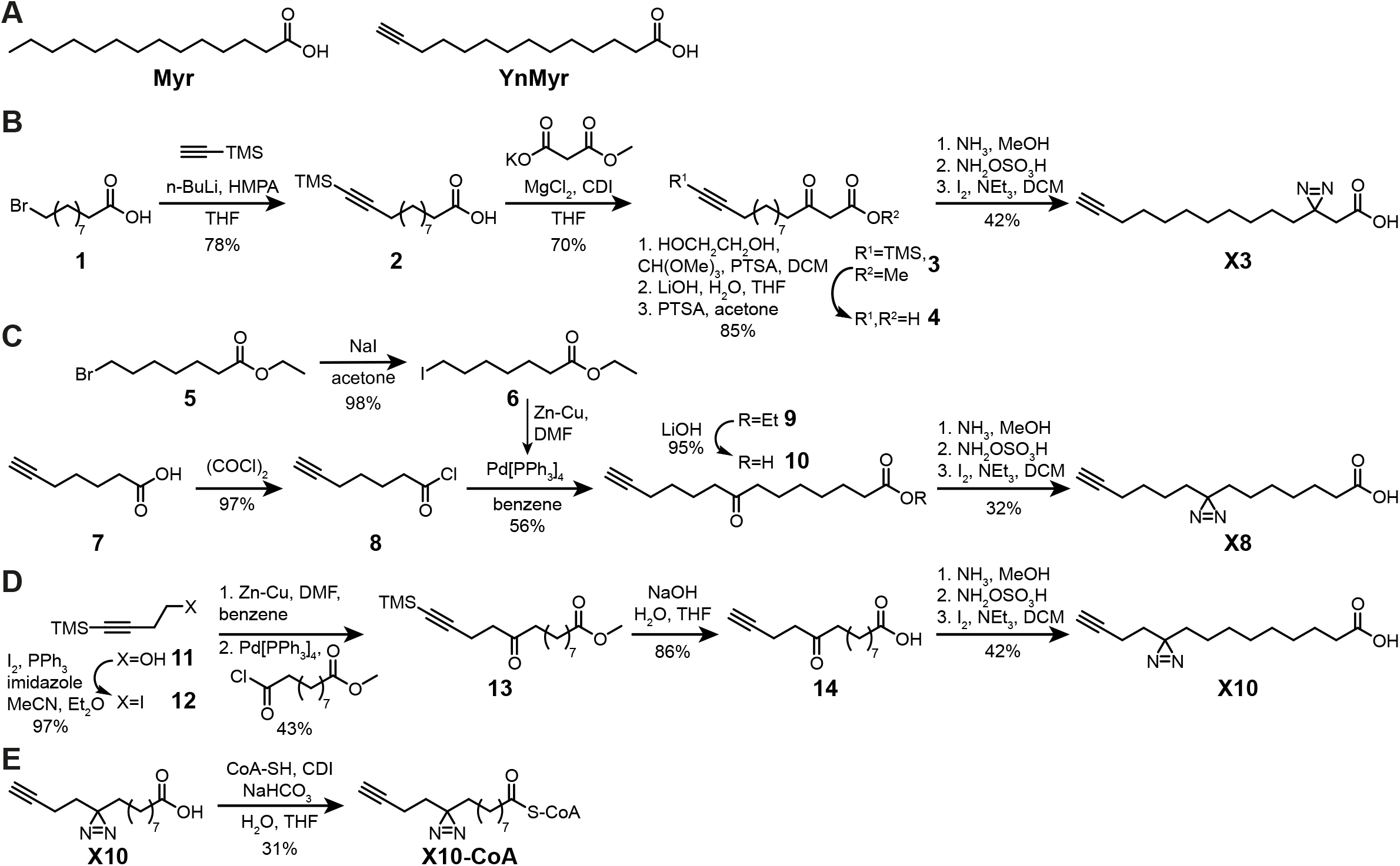
Synthesis of photocrosslinkable clickable myristate probes. **A**. Left, myristic acid (**Myr**). Right, **YnMyr** probe. **B-E**. Synthesis schemes for **X3 (B), X8 (C), X10 (D)** and **X10-CoA (E)**. Full details are provided in the Supplementary Information.

## Notes

https://www.rcsb.org/structure/5NPQ

### Competing Interest Statement

Edward W Tate is a founder, shareholder and Director of Myricx Pharma Ltd

