## Supporting Information. Materials and Methods, Supplementary Table S3 and Figures for "Discovery of lipid-mediated protein-protein interactions in living cells using metabolic labeling with photoactivatable clickable probes"

Supporting information contents

### Supplementary Figures and Tables

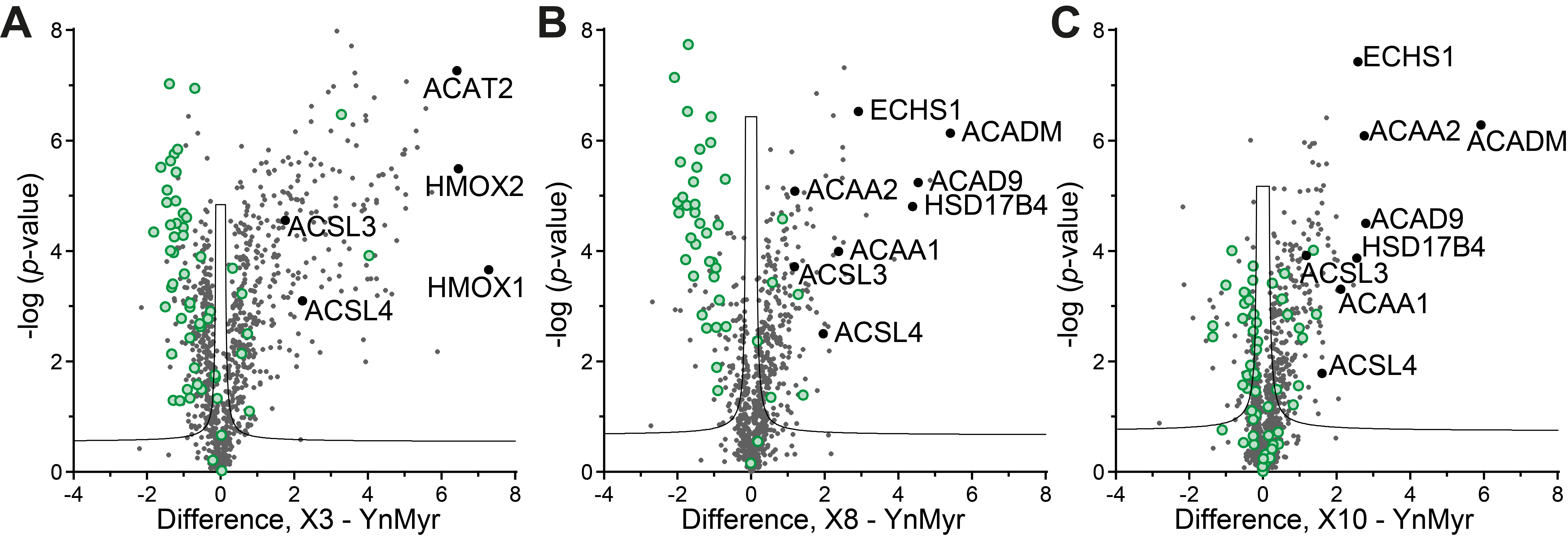

Figure S1. Lipid biosynthesis and metabolism pathway proteins are enriched with probes **X3**/**X8**/**X10**. **A-C**. Samples are in triplicates with relative enrichment plotted on X-axis against statistical significance on Y-axis. Photocrosslinkable clickable probes show differential enrichment of proteins in comparison to **YnMyr**. Fatty acid binding and metabolism proteins are highly enriched with **X3** (**A**), **X8** (**B**) and **X10** (**C**) probes, indicating background levels of non-irradiative crosslinking. **X10** probe shows smaller number and lower enrichment of fatty acid binding proteins. Myristoylated proteins (green circles) are enriched to significantly lower levels with **X3** and **X8** probes than with **YnMyr**. However, with **X10** probe, enrichment of myristoylated proteins is largely similar to **YnMyr**. Selected examples of fatty acid binding and metabolism proteins are indicated. False-discovery rate of 0.05 is shown as a black line.

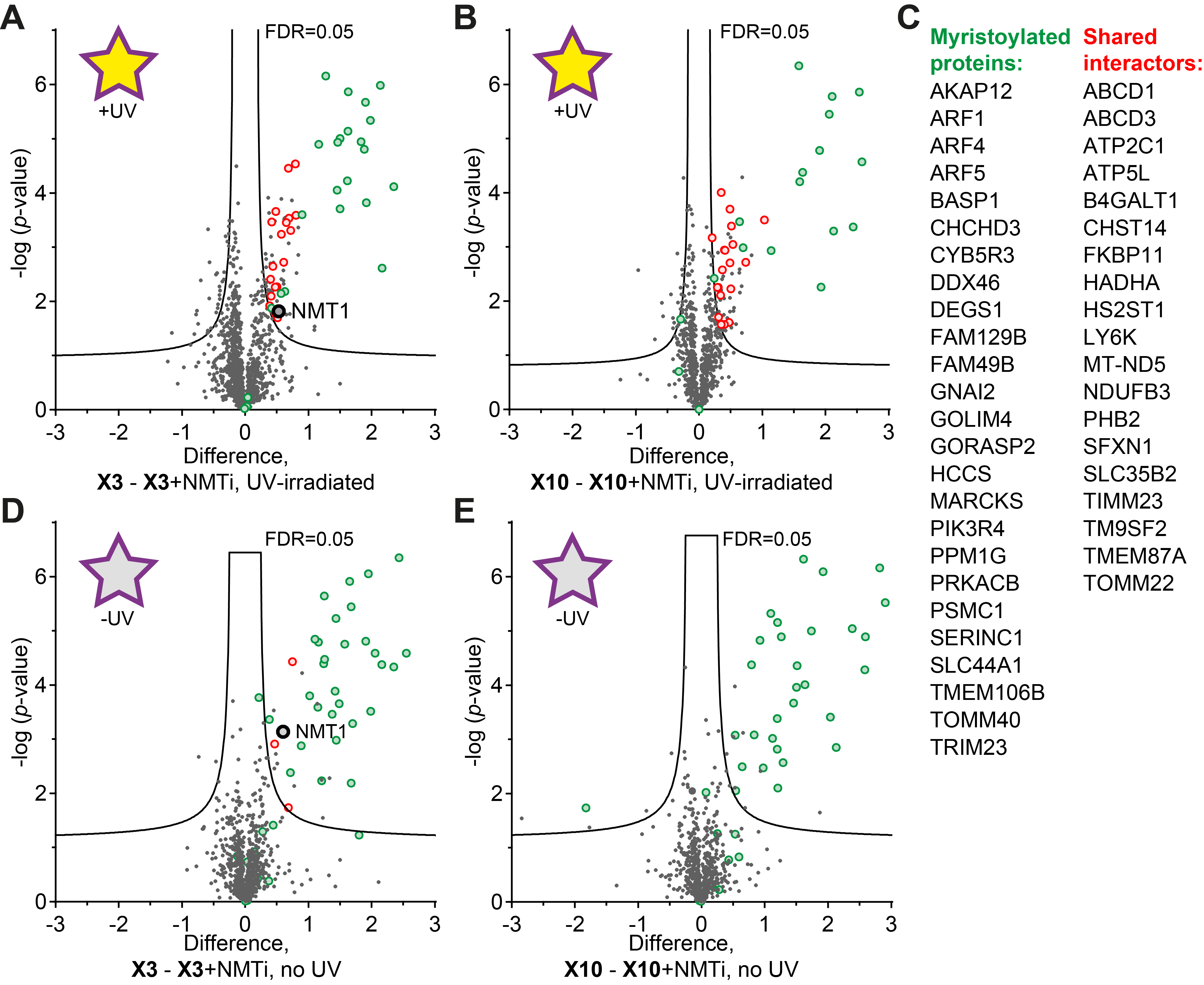

Figure S2. Identification of interactors of myristoylated proteins via photocrosslinking. **A-B**. HeLa cells were incubated with 100 µM probes **X3** (**A**) and **X10** (**B**) with or without an NMT inhibitor (NMTi). Cells were lysed in denaturing conditions and processed as shown on Figure 2A, followed by SILAC proteomics. Heavy-labelled spike-in sample was prepared from HeLa cells grown in the SILAC-heavy media and treated with 100 µM of corresponding probes, **X3** or **X10**. Samples are in triplicates with Light/Heavy ratio differences plotted on X‑axis against statistical significance on Y-axis. Several myristoylated proteins (green) were enriched compared to cells that were also incubated with 1 µM NMTi. Additionally, several non-myristoylated proteins (shared interactors) appeared as significantly enriched in both **X3** and **X10** experiments (indicated by red circles). **C**. Myristoylated proteins identified in the experiment (left) and proteins that appear significantly enriched in both **X3** and **X10** samples (shared interactors, right). **D-E**. For comparison are the same plots with probes **X3** (**D**) and **X10** (**E**) as on panels **A-B**, but without UV-irradiation. The data is from the SILAC proteomics experiment described in Figure 3A. The enrichment of concomitant proteins is much lower without UV crosslinking. However, NMT1 appears enriched with **X3** in both UV and no-UV conditions, correlating with results from Western blotting (Figure 2D). Three myristoylated proteins with higher enrichment and/or significance are not shown on panel **E** to preserve the scale.

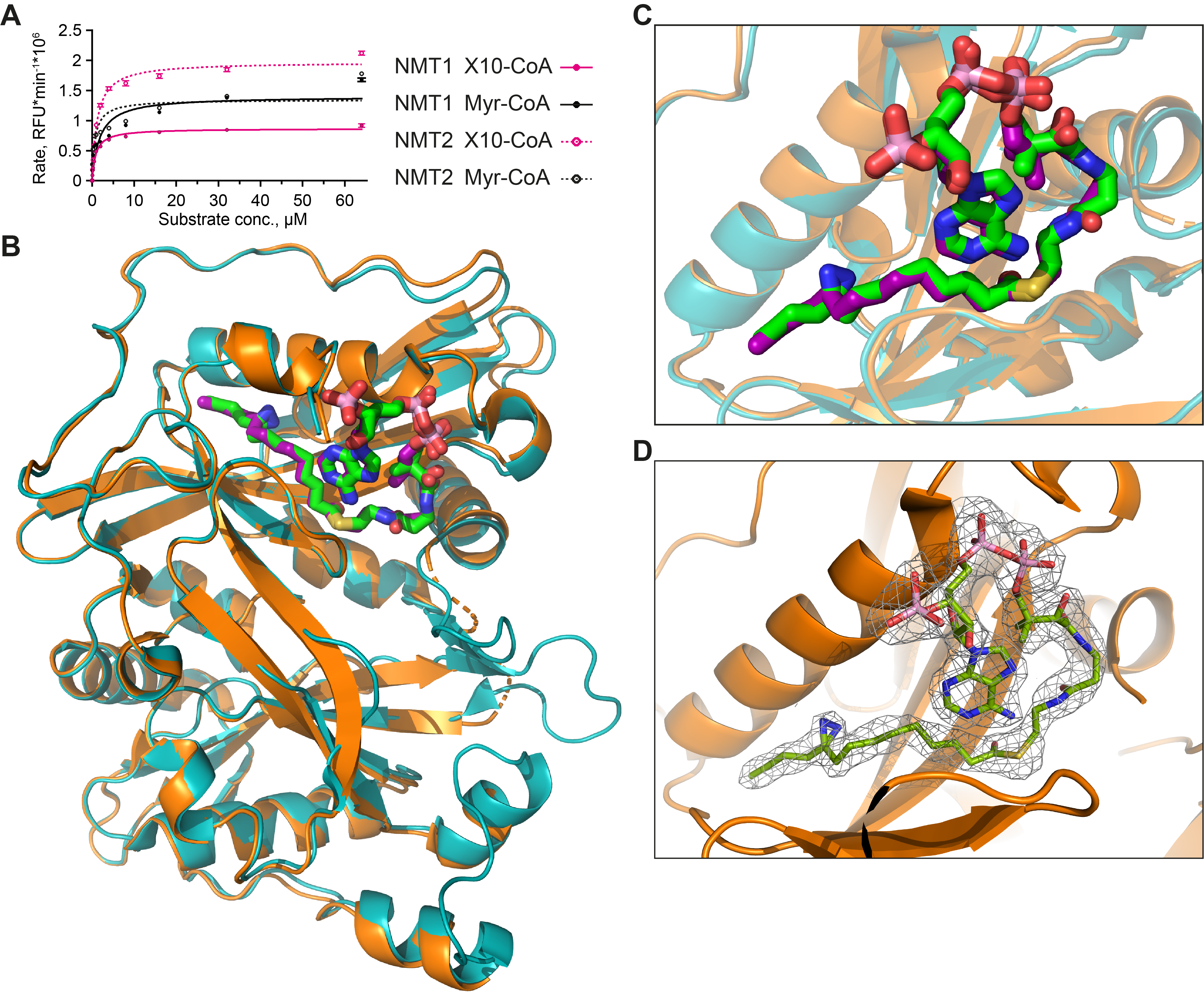

Figure S3. NMT enzymatic assay and crystal structure of HsNMT1:**X10-CoA**. **A**. Rates of enzymatic transfer of **Myr** or **X10** by HsNMT1 or HsNMT2 to a synthetic peptide based on c-Src N-terminal peptide (16 μM H‑GSNKSKPK‑NH2) were measured *in vitro* with CPM-based assay. **B-C**. Structural superimposition of NMT1 (orange) co-crystallized with **X10-CoA** (green) and NMT1 (cyan) co-crystallized with **Myr-CoA** (dark cyan) (PDB:4C2Y). RMSD value between both structures is 0.587 Å for 355 Cα atoms. Whole protein view (**B**) and close-up of the myristate pocket (**C**). **D**. 2Fo-Fc electron density map observed for **X10‑CoA** (chain A) contoured at sigma=1.0. Color code used is blue (nitrogen), red (oxygen), yellow (sulfur), and pink (phosphorus).

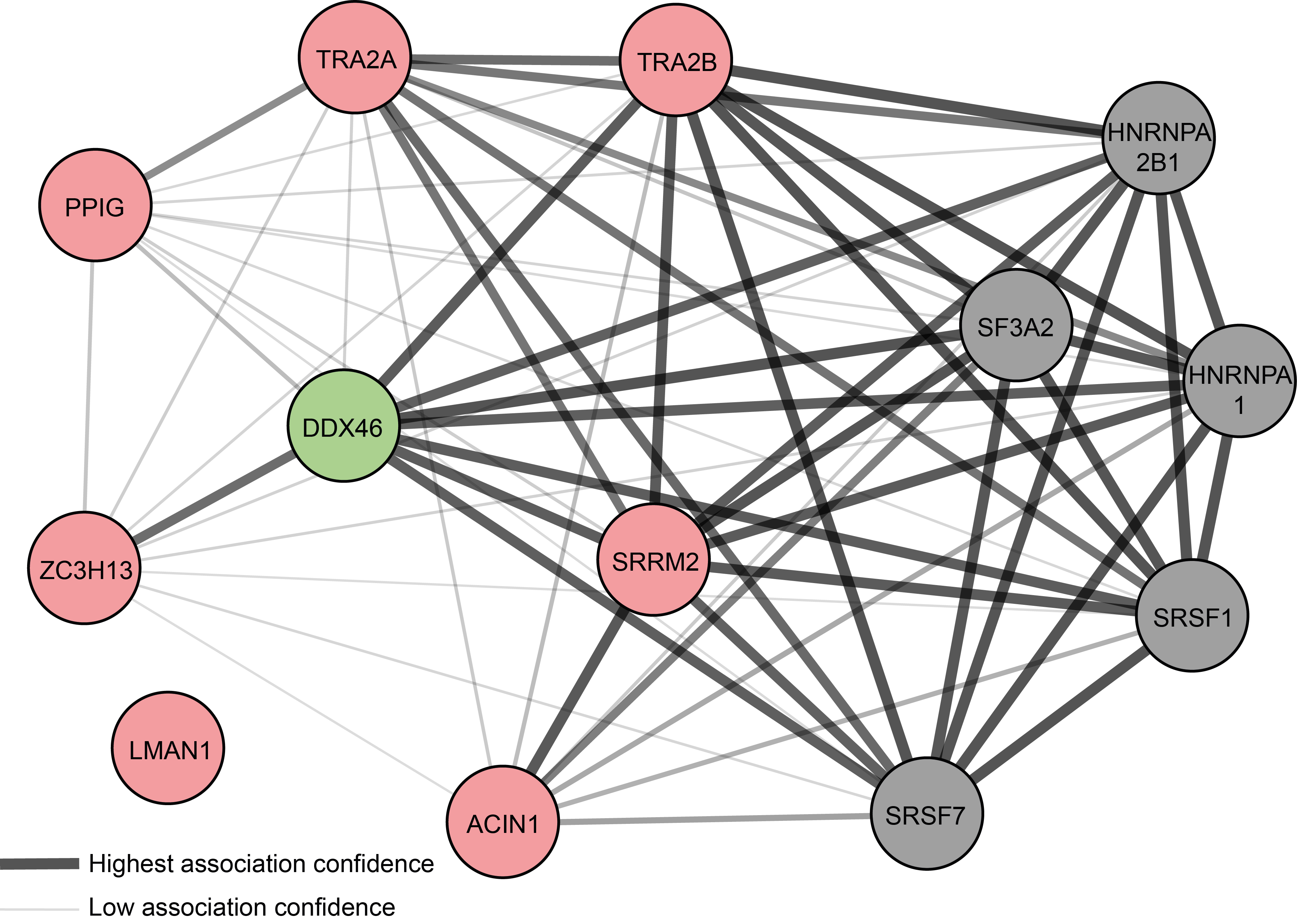

Figure S4**.** Association network of DDX46 (green) and its interactors identified in this work (red) based on analysis with StringDB [1]. Proteins that are high-confidence interactors that were not identified in our experiments are shown in grey. Lines indicate known protein associations.

Table S3. X-ray data collection and refinement statistics.

|  | HsNMT1-X10 |
| --- | --- |
| Data collection statistics |  |
| Space group | *P*2_1_2_1_2 |
| Number of complexes in ASU | 2 |
| Unit cell (Å) | *a*=79.97  *b*=177.45  *c*=58.24 |
| Wavelength (Å) | 0.97625 |
| Beamline | I03 (DIAMOND) |
| Resolution range (Å) | 177.46-2.37 (2.41-2.37) |
| *R*_merge_ | 0.191 (0.847) |
| *R*_pim_ | 0.076 (0.353) |
| Number of total reflections | 247356 (11404) |
| Number of unique reflections | 34474 (1709) |
| Mean ([I]/SD[I]) | 9.3 (2.2) |
| CC_1/2_ | 0.994 (0.661) |
| Completeness (%) | 100 (99.7) |
| Multiplicity | 7.2 (6.7) |
| Wilson plot B (Å2) | 32.66 |
| Refinement |  |
| Number of atoms | 6174 |
| Protein | 5889 |
| X10 | 130 |
| Ions / Glycerol | 2 Mg^2+^, 24 |
| Water | 129 |
| *R*_work_ (%) | 19.9 |
| *R*_free_ (%) | 26.1 |
| <B > (Å^2^) | 34.61 |
| rms bond length deviation (Å) | 0.008 |
| rmsd angle deviation (°) | 1.037 |

Values in parentheses correspond to the highest resolution shell.

*R*_merge_ = Σ(I_hl_ − < I_h_ >)/Σ < I_h_ >

*R*_meas_ = Σ√(n_h_/n_h_ – 1)(I_hl_ − < I_h_ >)/Σ < I_h_ >

*R*_pim_ = Σ√(1/n_h_ – 1)(I_hl_ − < I_h_ >)/Σ < I_h_ >

### Materials and Methods

#### General chemistry methods

All chemical reagents and solvents were purchased from commercial sources and used as supplied without any further purification. Dry solvents were dispensed using Pure SolvTM solvent drying towers (Innovative technology Inc.) or purchased anhydrous from commercial sources. All anhydrous conditions were performed in oven-dried glassware under argon atmosphere. For purifications, HPLC-grade solvents (≥ 99% purity) were used (Merck or Fisher Scientific UK). Room temperature (RT) reactions were carried out at 15-20°C. The term *in vacuo* refers to solvent removal using Büchi rotary evaporation in a water bath at 15-50°C, at ~ 10 mm Hg.

Reactions were monitored by thin-layer chromatography (TLC), using TLC plates pre-coated with silica gel 60 F_254_ on aluminium (Merck). TLC plates were visualized using UV light (254 nm) or by using the appropriate chemical stain. Flash column chromatography was carried out using Geduran® Si 60 (40-63 μm) silica gel (Merck). The purification and analysis of conjugate **X10-CoA** was performed on a Waters RP-HPLC system (Waters 2767 autosampler; Waters 515 HPLC pump; Waters 3100 mass spectrometer with ESI; Waters 2998 photodiode array (detection from 190 to 700 nm)), equipped with Xbridge C18 reverse-phase columns with dimensions 4.6 mm × 100 mm for analytical (1.2 mL/min flow rate) and 19 mm × 100 mm for preparative (20 mL/min flow rate) runs.

Nuclear magnetic resonance (NMR) spectra were recorded at ambient temperature on a Bruker AV400 instrument operating at 400 MHz for ^1^H and 101 MHz for ^13^C in the deuterated chloroform CDCl_3_ (δ_H_ = 7.26 and δ_C_ = 77.16) as the internal standard. Chemical shifts (δ) are reported in parts per million (ppm) relative to tetramethylsilance (TMS) as a reference and coupling constants (*J*) in Hertz (Hz). The multiplicity of each signal is indicated as s-singlet, d-doublet, t-triplet, q-quartet, quint-quintet, m-multiplet or a combination of these. All assignments were made with the help of COSY, HSQC, HMBC, NOESY or DEPT correlation spectra. Spectra were processed and analysed using MestReNova© 12 NMR software. High-resolution mass spectra (HRMS) were recorded on a Waters LCT Premier mass spectrometer, operating in electrospray ionization (ESI) (positive (+) and negative (-)) mode at the Department of Chemistry, Imperial College London. The *m/z* values are reported in Daltons.

#### Synthesis

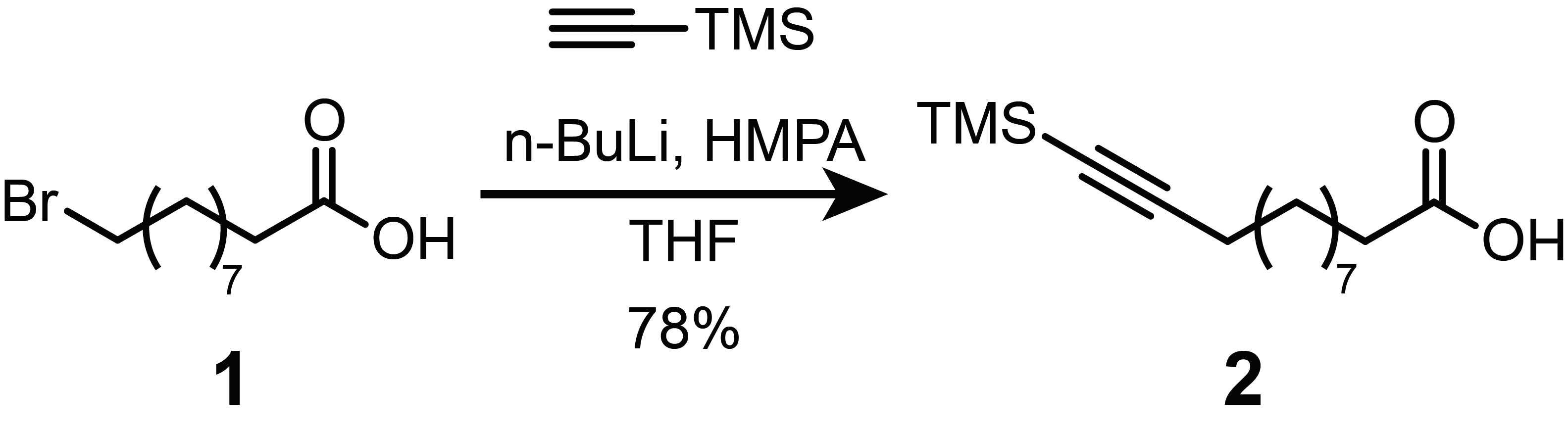

**Preparation of 12-(trimethylsilyl)dodec-11-ynoic acid (2)**. The procedure is modified from [2]. Trimethylsilylacetylene (608 µL, 4.4 mmol) in dry THF (12 mL) was cooled to -78°C in dry ice-acetone bath. A solution of n‑BuLi in hexane (2.5 M, 1.76 mL, 4.4 mmol) was added dropwise under argon and stirred for 20 min. 10‑Bromodecanoic acid **1** (502 mg, 2.0 mmol) was dissolved in dry THF (4 mL) and added to the reaction mixture followed by addition of hexamethylphosphoramide (HMPA) (4 mL, 22 mmol). The solution was stirred overnight and allowed to warm up to RT. The reaction mixture was quenched with saturated NH_4_Cl _(aq.)_, the pH was adjusted to 3 (with 1 M HCl). The product was extracted with EtOAc and washed with brine. The organic phase was dried over Na_2_SO_4_ (anh.) and concentrated *in vacuo*. The crude residue was purified by flash chromatography eluting with 12:87:1 EtOAc:Hexane:AcOH mixture, to give acid **2** (330 mg, 62%).^1^H NMR (400 MHz, CDCl_3_): δ 2.35 (t, *J* = 7.5 Hz, 2H), 2.21 (t, *J* = 7.1 Hz, 2H), 1.63 (quint, *J* = 7.5 Hz, 2H), 1.50 (quint, *J* = 7.2 Hz, 2H), 1.41-1.23 (m, 10H), 0.14 (s, 9H). ^13^C NMR (101 MHz, CDCl_3_): δ 179.68, 107.88, 84.42, 34.08, 29.40, 29.31, 29.17, 29.15, 28.90, 28.75, 24.81, 19.99, 0.33.

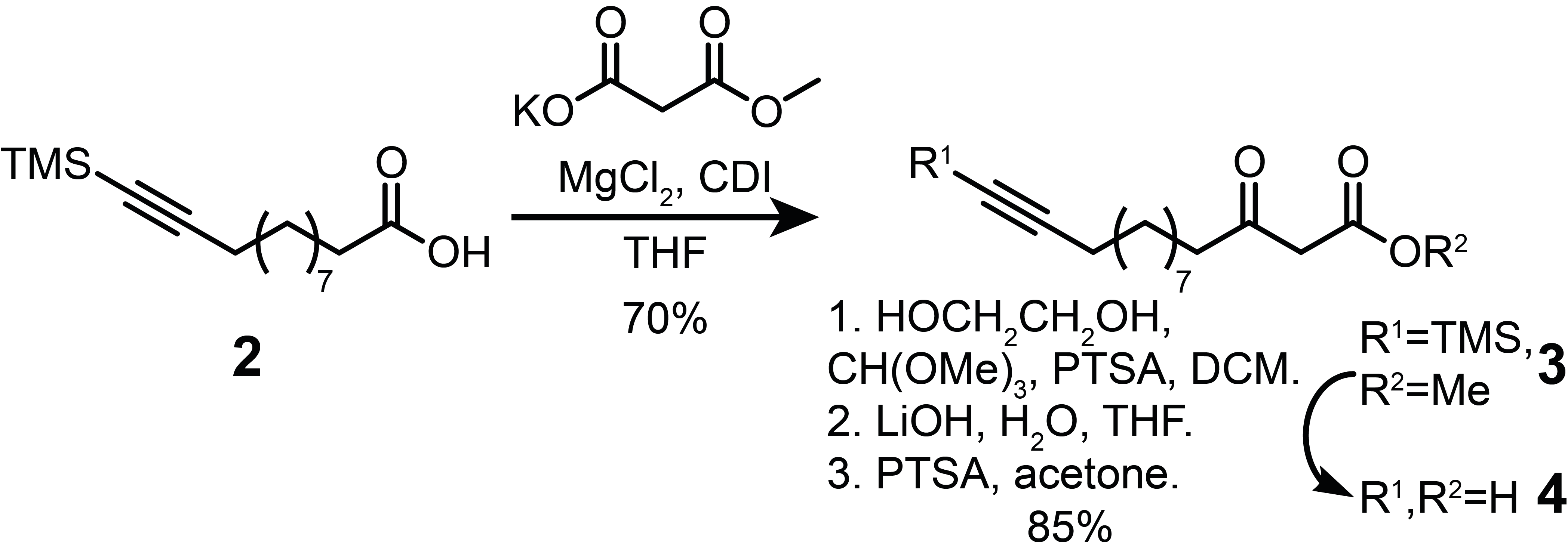

**Preparation of methyl 3-oxo-14-(trimethylsilyl)tetradec-13-ynoate (3).** The procedure is modified from [3]. A suspension of MgCl_2_ (1.15 g, 7.6 mmol) and methyl potassium malonate (2.48 g, 15.9 mmol) in dry THF (20 mL) was stirred at 50°C for 4 h. Separately, carbonyldiimidazole (CDI) (1.47 g, 9.1 mmol) was added to the solution of acid **2** (2.03 g, 7.6 mmol) in dry THF (20 mL) in 4°C ice bath. The mixture was allowed to warm up to RT, stirred for 1 h and then added dropwise to methyl magnesium malonate suspension. The resulting mixture was stirred overnight, then concentrated *in vacuo* and resuspended in EtOAc. The solution was washed with 1 M KHSO_4 (aq.)_, NaHCO_3 (aq.)_ and brine. The organic phase was dried over Na_2_SO_4_ (anh.) and concentrated *in vacuo*. The crude residue was purified by flash chromatography eluting with 1:4 EtOAc:Hexane mixture, to give β-keto ester **3** (1.968 g, 80%). .H

^1^H NMR (400 MHz, CDCl_3_): δ 3.73 (s, 3H), 3.44 (s, 2H), 2.52 (t, *J* = 7.5 Hz, 2H), 2.19 (t, *J* = 7.2 Hz, 2H), 1.58 (quint, *J* = 7.5 Hz, 2H), 1.49 (quint, *J* = 7.3 Hz, 2H), 1.40-1.22 (m, 10H), 0.13 (s, 9H). ^13^C NMR (101 MHz, CDCl_3_): δ 202.94, 167.83, 107.85, 84.39, 52.45, 49.15, 43.20, 29.40, 29.38, 29.13, 29.10, 28.88, 28.73, 23.57, 19.97, 0.31. HRMS (ESI^+^, m/z): calcd. for C_18_H_32_O_3_NaSi [M+Na]^+^ 347.2018; found 347.2033.

**Preparation of 3-oxotetradec-13-ynoic acid (4)**. β-Keto ester **3** (1.47 g, 4.5 mmol) was dissolved in DCM (15 mL), followed by addition of ethylene glycol (1.41 g, 22.7 mmol), *p*-toluenesulfonic acid hydrate (PTSA) (86 mg, 0.45 mmol) and trimethyl orthoformate (1.01 g, 9.5 mmol). The mixture was refluxed overnight and quenched with saturated NaHCO_3_ _(aq.)_ (15 mL). The crude product was extracted with DCM and washed with brine. The organic phase was dried over Na_2_SO_4_ (anh.) and concentrated *in vacuo* to obtain a glycol-protected intermediate **3’** in quantitative yield (1.65 g, 100%), that was used without further purification.

A glycol-protected intermediate **3’** (1.65 g, 4.5 mmol) was dissolved in THF (54 mL), 1 M LiOH _(aq.)_ (18 mL) was added and the mixture was stirred at 60°C for 4 h. The pH was then adjusted to 3 (with 1 M HCl). The crude product was extracted with DCM and washed with brine. The organic phase was dried over Na_2_SO_4_ (anh.) and concentrated *in vacuo* to obtain acid intermediate **3’’** (1.17 g, 92%).

The acid intermediate **3’’** (402 mg, 1.43 mmol) was dissolved in acetone (5 mL), PTSA (68 mg, 0.36 mmol) was added, and the mixture was stirred for 2 h at RT. EtOAc was added, and the solution was washed with H_2_O and brine. The organic phase was dried over Na_2_SO_4_ (anh.) and concentrated *in vacuo*. The crude residue was purified by flash chromatography eluting with 20:80:1 EtOAc:Hexane:AcOH mixture, to yield acid **4** (317 mg, 93 %). ^1^H NMR (400 MHz, CDCl_3_): δ 3.52 (s, 2H), 2.56 (t, *J* = 7.5 Hz, 2H), 2.18 (td, *J* = 7.1, 2.7 Hz, 2H), 1.94 (t, *J* = 2.6 Hz, 1H), 1.67-1.57 (m, 2H), 1.57-1.47 (m, 2H), 1.43-1.22 (m, 10H). HRMS (ESI^-^, m/z): calcd. for C_14_H_21_O_3_ [M-H]^-^ 237.1491; found 237.1500.

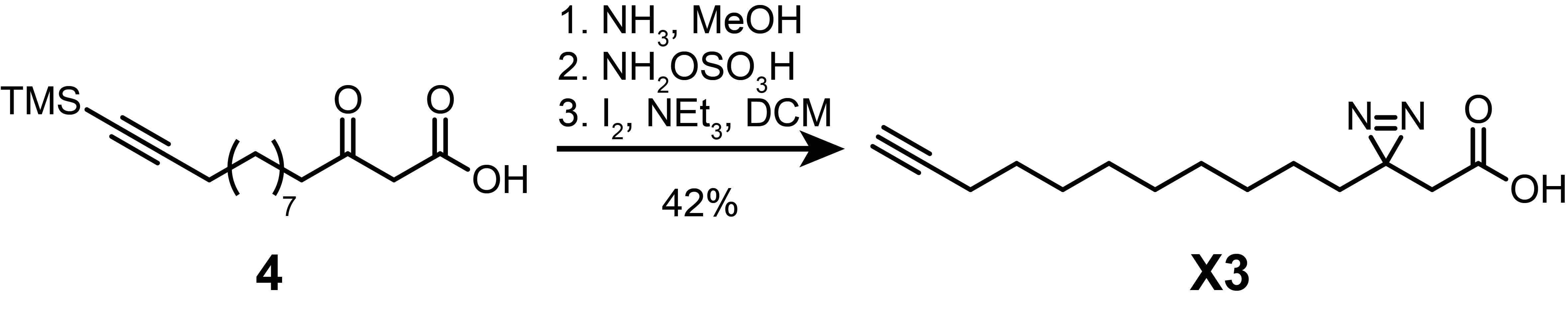

**Preparation of 2-(3-(undec-10-yn-1-yl)-3H-diazirin-3-yl)acetic acid (X3)**. The procedure is modified from [4]. Concentrated NH_3_ _(aq.)_ was boiled in a 3-neck flask and carried under the flow of argon through a gas trap with NaOH pellets. Liquid NH_3_ (~ 5 mL) was collected into a 3-neck flask equipped with a dry ice-acetone condenser at -78°C. Acid **4** (150 mg, 0.75 mmol) was dissolved in dry MeOH (5 mL) and added to the liquid NH_3_. The mixture was kept stirring at -30 to -40°C for 3 h by addition of dry ice. Then, hydroxylamine-*O*-sulfonic acid (94 mg, 0.83 mmol) was dissolved in dry MeOH (2 mL) and added to the reaction mixture over 30 min at -78°C in dry ice-acetone bath. The reaction was left to warm up to RT overnight. Next, argon was bubbled through the solution for 1 h to volatilize leftover NH_3_. The solution was filtered through sintered glass, and the residue was washed twice with dry MeOH. The filtrate was concentrated *in vacuo* at RT and the residue was cooled on ice and dissolved in DCM (5 mL), followed by NEt_3_ (220 µL, 2.25 mmol). Iodine (~1 g) solution in DCM (10 mL) was added dropwise until brown color persisted and the mixture was left stirring at RT for 1 h. The solution was then washed with 1 M HCl (5 mL), 1 M Na_2_S_2_O_3 (aq.)_ (5 mL) and brine (5 mL). The organic phase was dried over Na_2_SO_4_ (anh.) and concentrated *in vacuo*. The crude residue was purified by flash chromatography eluting with 20:80:1 EtOAc:Hexane:AcOH mixture, yielding diazirine **X3** as a white powder (66 mg, 42%).^1^H NMR (400 MHz, CDCl_3_): δ 2.30 (s, 2H), 2.17 (td, *J* = 7.1, 2.7 Hz, 2H), 1.94 (t, *J* = 2.6 Hz, 1H), 1.59-1.47 (m, 4H), 1.43-1.32 (m, 2H), 1.31-1.18 (m, 8H), 1.14-1.03 (m, 2H). ^13^C NMR (101 MHz, CDCl_3_): δ 175.20, 84.91, 68.22, 39.62, 32.53, 29.41, 29.38, 29.14 (two overlapping peaks), 28.83, 28.59, 26.02, 23.86, 18.53. HRMS (ESI^-^, m/z): calcd. for C_14_H_21_N_2_O_2_ [M-H]^-^ 249.1603; found 249.1597.

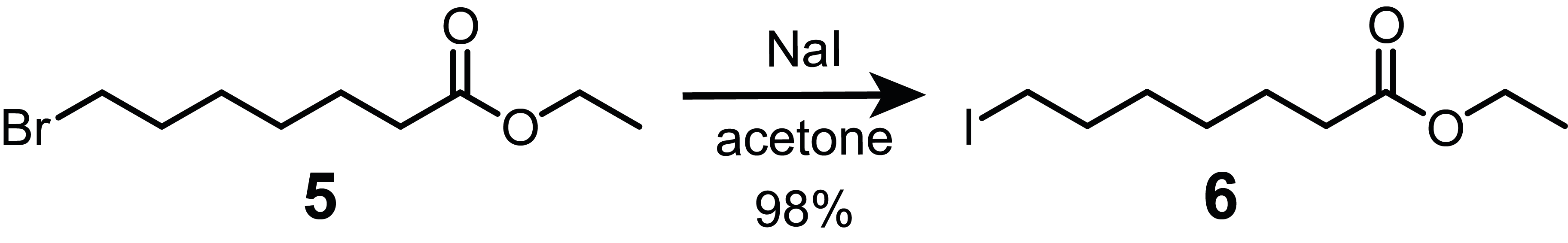

**Preparation of ethyl 7-iodoheptanoate (6)**. To a clear solution of ethyl 7-bromoheptanoate **5** (6.08 g, 25.7 mmol) in acetone (50 mL) was added NaI (11.54 g, 77 mmol). The mixture was stirred at RT overnight. Completion of the reaction was checked by ^1^H NMR. After full conversion, the mixture was filtered through sintered glass and concentrated *in vacuo,* followed by addition of H_2_O (100 mL) and DCM (100 mL). The layers were separated, and the aqueous layer was extracted again with DCM (100 mL). The combined organic layers were washed with 1 M Na_2_S_2_O_3 (aq.)_ (50 mL), brine (50 mL), dried over Na_2_SO_4_ (anh.) and concentrated *in vacuo,* yielding the crude iodide **6** (7.1 g, 98%), that was used without further purification. ^1^H NMR (400 MHz, CDCl_3_): δ 4.12 (q, *J* = 7.1 Hz, 2H), 3.18 (t, *J* = 7.0 Hz, 2H), 2.29 (t, *J* = 7.5 Hz, 2H), 1.82 (quint, *J* = 7.0 Hz, 2H), 1.63 (quint, *J* = 7.3 Hz, 2H), 1.48-1.30 (m, 4H), 1.25 (t, *J* = 7.1 Hz, 3H).

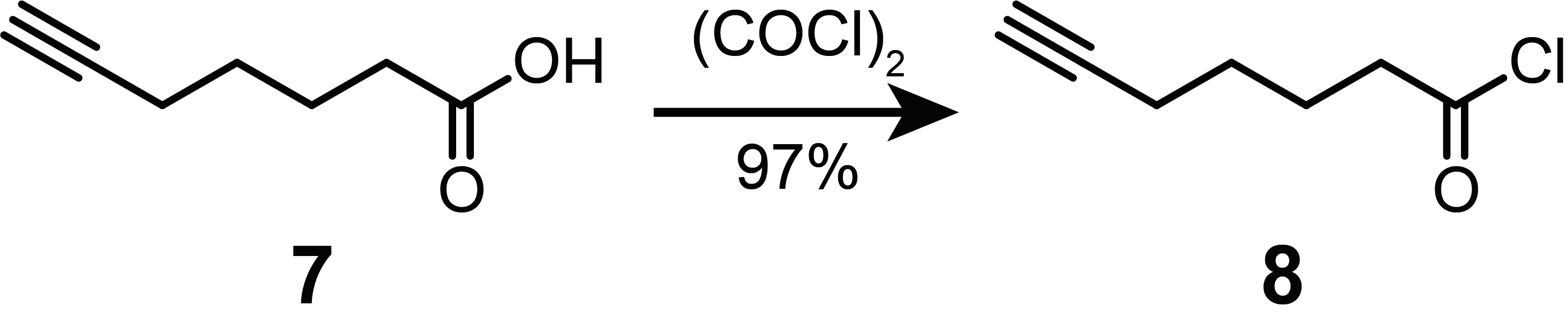

**Preparation of hept-6-ynoyl chloride (8)**. The procedure is modified from [5]. Oxalyl chloride solution in DCM (2 M, 17.5 mL, 35 mmol) was mixed with a solution of 6-pentynoic acid **7** (0.63 g, 5 mmol) in DCM (50 mL) under argon, the mixture was heated to 70˚C, refluxed for 1 h and stirred overnight at RT. The volatiles were evaporated *in vacuo*, yielding the crude chloride **8** in quantitative yield (0.72 g), that was used without further purification. ^1^H NMR (400 MHz, CDCl_3_): δ 2.93 (t, *J* = 7.2 Hz, 2H), 2.23 (td, *J* = 7.0, 2.7 Hz, 2H), 1.97 (t, *J* = 2.6 Hz, 1H), 1.89-1.80 (m, 2H), 1.64-1.55 (m, 2H).

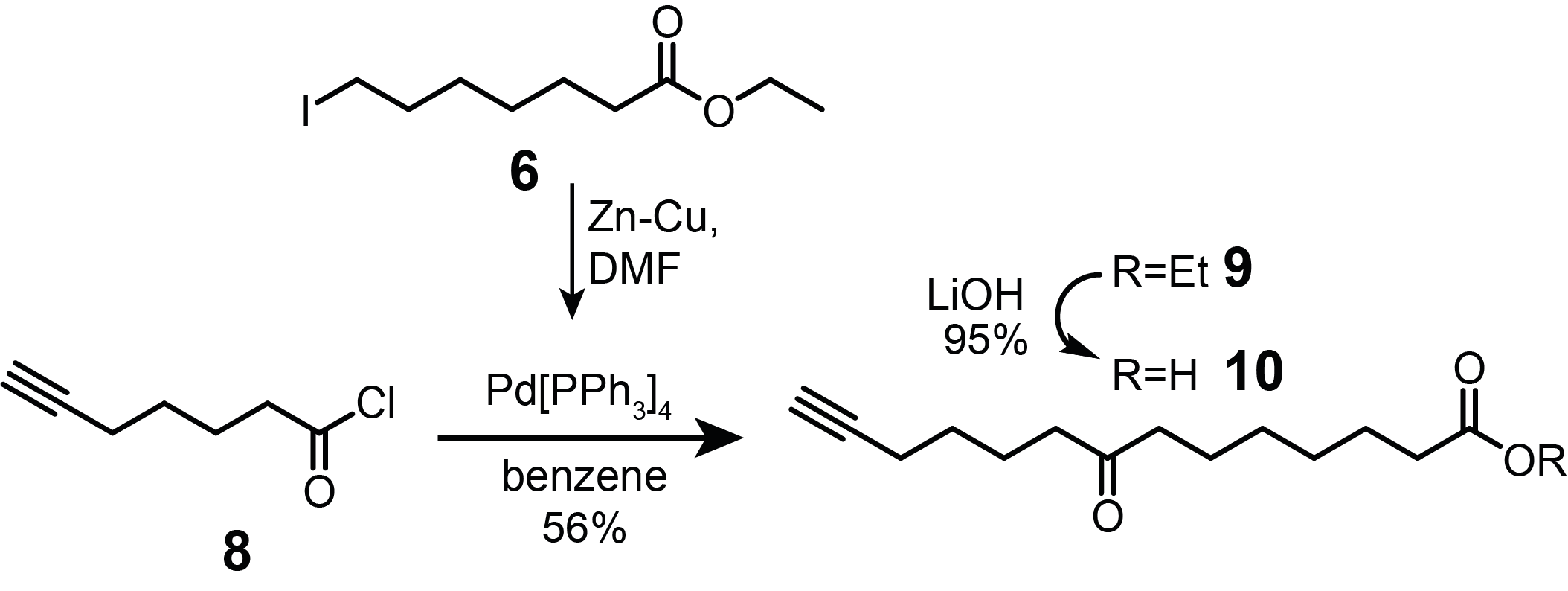

**Preparation of ethyl 8-oxotetradec-13-ynoate (9)**. The procedure is modified from [6]. Zn-Cu powder (1-3% Cu, 392 mg, 6 mmol) was dried at 150°C *in vacuo*. Next, iodide **6** (1.42 g, 5 mmol), dry benzene (20 mL) and DMF (0.5 mL, 6.5 mmol) were mixed under argon at RT, the solution was heated to 80°C and stirred for 5 h to form alkylzinc iodide. In the second flask, chloride **8** (0.65 g, 4.5 mmol) and freshly prepared Pd[PPh_3_]_4_ [7] (174 mg, 0.15 mmol) were mixed and dissolved in dry benzene (15 mL) under argon. The solution of alkylzinc iodide was cooled down to RT and added to the second flask via a syringe pump over the course of 40 min and left to stir overnight at RT. The reaction was quenched with saturated NH_4_Cl _(aq.)_ (50 mL), followed by addition of EtOAc (50 mL). The layers were separated, and the aqueous layer was extracted again with EtOAc (50 mL). The combined organic phases were washed with brine, dried over Na_2_SO_4_ (anh.) and concentrated *in vacuo*. The crude residue was purified by flash chromatography eluting with a three-step gradient of 7-10-12% of EtOAc in hexane, obtaining ethyl ester **9** as brown transparent crystals (673 mg, 57%). ^1^H NMR (400 MHz, CDCl_3_): δ 4.10 (q, *J* = 7.2 Hz, 2H), 2.42-2.35 (m, 4H), 2.26 (t, *J* = 7.5 Hz, 2H), 2.18 (td, *J* = 7.0, 2.6 Hz, 2H), 1.93 (t, *J* = 2.7 Hz, 1H), 1.71-1.45 (m, 8H), 1.33-1.26 (m, 4H), 1.23 (t, *J* = 7.1 Hz, 3H). HRMS (ESI^+^, m/z): calcd. for C_16_H_27_O_3_ [M+H]^+^ 267.1960; found 267.1973.

**Preparation of 8-oxotetradec-13-ynoic acid (10)**. Ethyl ester **9** (1.36 g, 5.1 mmol) was dissolved in THF (45 mL), followed by 1 M LiOH _(aq.)_ (15 mL) and the mixture was stirred overnight at RT. Next, the pH was adjusted to 2 (with 1 M HCl). The crude product was extracted with EtOAc (×3) and washed with brine. The organic phase was dried over Na_2_SO_4_ (anh.) and concentrated *in vacuo*. The crude residue was purified by flash chromatography eluting with 20:80:1 EtOAc:Hexane:AcOH mixture, yielding acid **10** (1.15 g, 95%). ^1^H NMR (400 MHz, CDCl_3_): δ 2.44-2.37 (m, 4H), 2.34 (t, *J* = 7.5 Hz, 2H), 2.19 (td, *J* = 7.0, 2.7 Hz, 2H), 1.94 (t, *J* = 2.6 Hz, 1H), 1.74-1.47 (m, 8H), 1.39-1.23 (m, 4H). HRMS (ESI^+^, m/z): calcd. for C_14_H_23_O_3_ [M+H]^+^ 239.1647; found 239.1644.

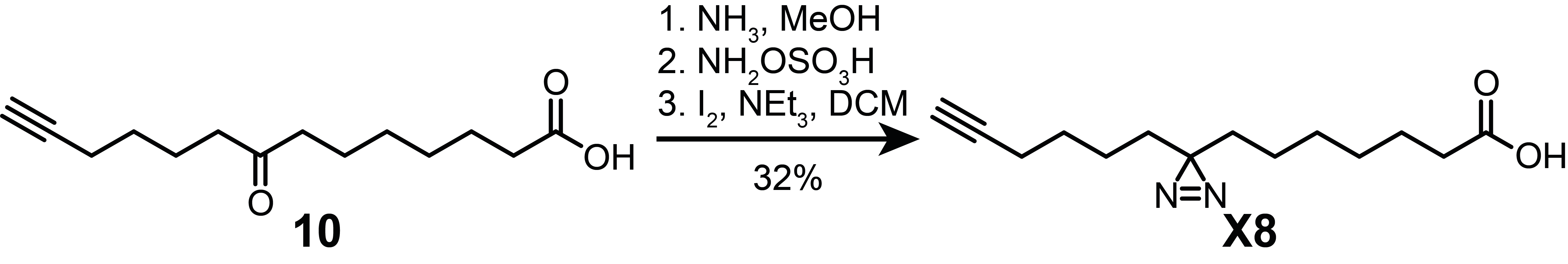

**Preparation of 7-(3-(hex-5-yn-1-yl)-3H-diazirin-3-yl)heptanoic acid (X8)**. Diazirine **X8** was synthesized following the same procedure as described for diazirine **X3** (see above) starting with acid **10** (196 mg, 0.82 mmol). The crude residue was purified using flash chromatography eluting with 15:85:1 EtOAc:Hexane:AcOH mixture, yielding diazirine **X8** as a white powder (63 mg, 32%). ^1^H NMR (400 MHz, CDCl_3_): δ 2.33 (t, *J* = 7.4 Hz, 2H), 2.14 (td, *J* = 7.0, 2.7 Hz, 2H), 1.94 (t, *J* = 2.7 Hz, 1H), 1.60 (quint, *J* = 7.4 Hz, 2H), 1.46 (quint, *J* = 7.4 Hz, 2H), 1.39-1.32 (m, 4H), 1.32-1.15 (m, 6H), 1.12-1.03 (m, 2H). ^13^C NMR (101 MHz, CDCl_3_): δ 180.24, 84.11, 68.68, 34.07, 32.85, 32.57, 28.91, 28.90, 28.73, 28.02, 24.58, 23.75, 23.10, 18.32. HRMS (ESI^+^, m/z): calcd. for C_14_H_23_N_2_O_2_ [M+H]^+^ 251.1760; found 251.1758.

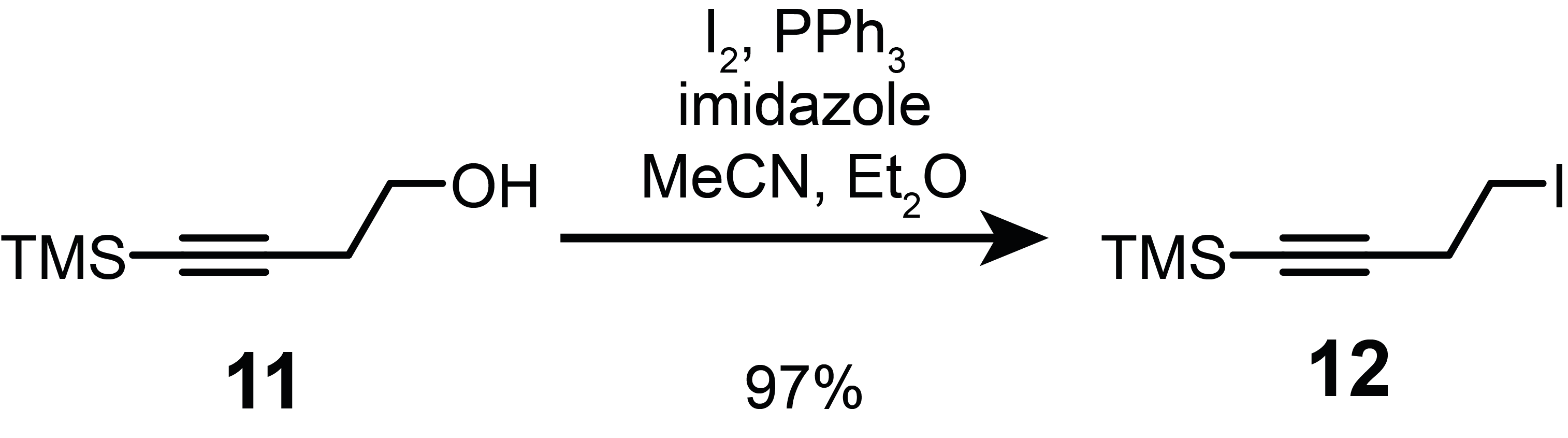

**Preparation of (4-iodobut-1-yn-1-yl)trimethylsilane (12)**. 4-(Trimethylsilyl)but-3-yn-1-ol **11** (2.44 mL, 15 mmol) was dissolved in the mixture of MeCN (20 mL) and Et_2_O (60 mL). Triphenylphosphine (5.25 g, 20 mmol), imidazole (1.36 g, 20 mmol) and iodine (5.08 g, 20 mmol) were added to the solution. The reaction mixture was stirred for 2 h at RT, followed by quenching with saturated NaHCO_3_ _(aq.)_. The organic phase was washed with 1 M Na_2_S_2_O_3 (aq.)_, brine and dried over Na_2_SO_4_ (anh.). The solution was passed through a layer of silica on a glass filter and washed off with hexane. The filtrate was concentrated *in vacuo* to obtain iodide **12** (3.67 g, 97%) that was used without further purification. ^1^H NMR (400 MHz, CDCl_3_): δ 3.22 (t, *J* = 7.5 Hz, 2H), 2.79 (t, *J* = 7.5, 2H), 0.16 (s, 9H).

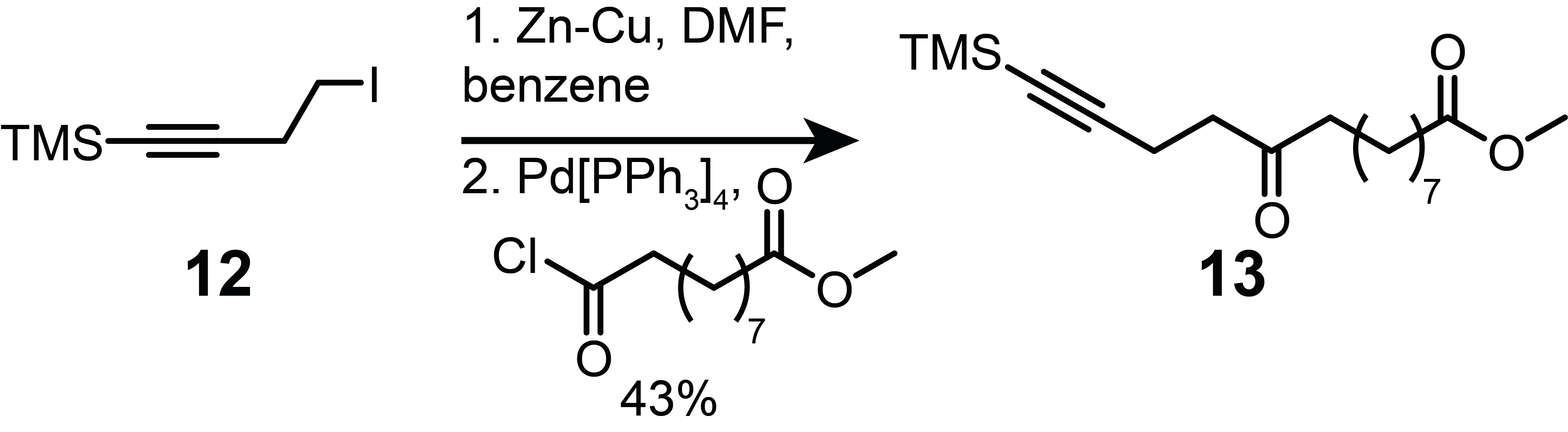

**Preparation of methyl 10-oxo-14-(trimethylsilyl)tetradec-13-ynoate (13)**. Ethyl ester **13** was synthesized following the same procedure as described for the ester **9** (see above) starting with iodide **12** (2.52 g, 10 mmol) and methyl 10-chloro-10-oxodecanoate (1.91 g, 8.1 mmol). The crude residue was purified using flash chromatography eluting with 5:95 EtOAc:Hexane mixture, yielding ethyl ester **13** (1.25 g, 43%). The product was ~78% pure due to the methyl 10-chloro-10-oxodecanoate contamination (calculated by ^1^H NMR). ^1^H NMR (400 MHz, CDCl_3_): δ 3.65 (s, 3H), 2.65-2.59 (m, 2H), 2.49-2.43 (m, 2H), 2.40 (t, *J* = 7.4 Hz, 2H), 2.28 (t, *J* = 7.5 Hz, 2H), 1.64-1.51 (m, 4H), 1.32-1.22 (m, 8H), 0.11 (s, 9H).

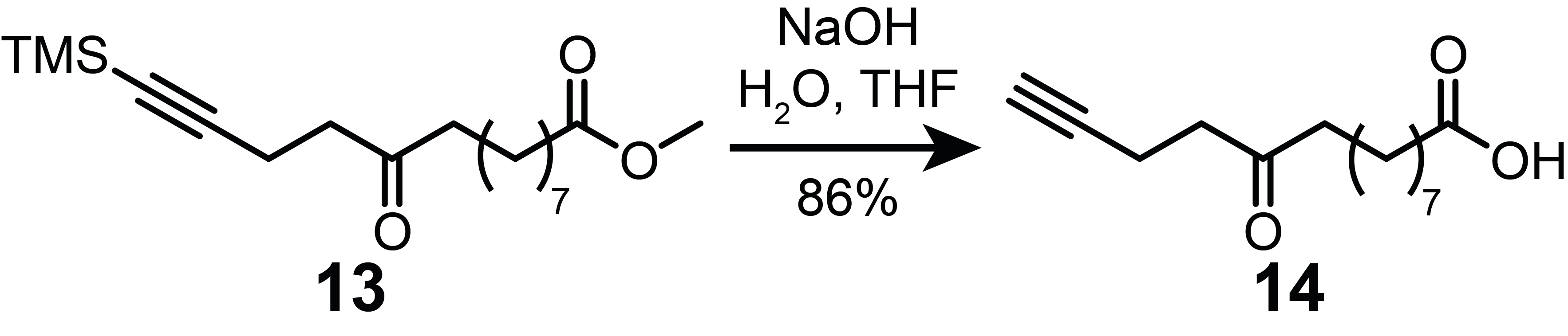

**Preparation of 10-oxotetradec-13-ynoic acid (14)**. Acid **14** was synthesized following the same procedure as described for the acid **10** (see above) starting with ethyl ester **13** (1.25 g, 4.4 mmol). The crude residue was purified using flash chromatography eluting with 20:80:1 EtOAc:Hexane:AcOH mixture, yielding acid **14** (0.79 g, 86%). ^1^H NMR (400 MHz, CDCl_3_): δ 2.68-2.63 (m, 2H), 2.47-2.39 (m, 4H), 2.33 (t, *J* = 7.5 Hz, 2H), 1.94 (t, *J* = 2.7 Hz, 1H), 1.67-1.53 (m, 4H), 1.36-1.23 (m, 8H).

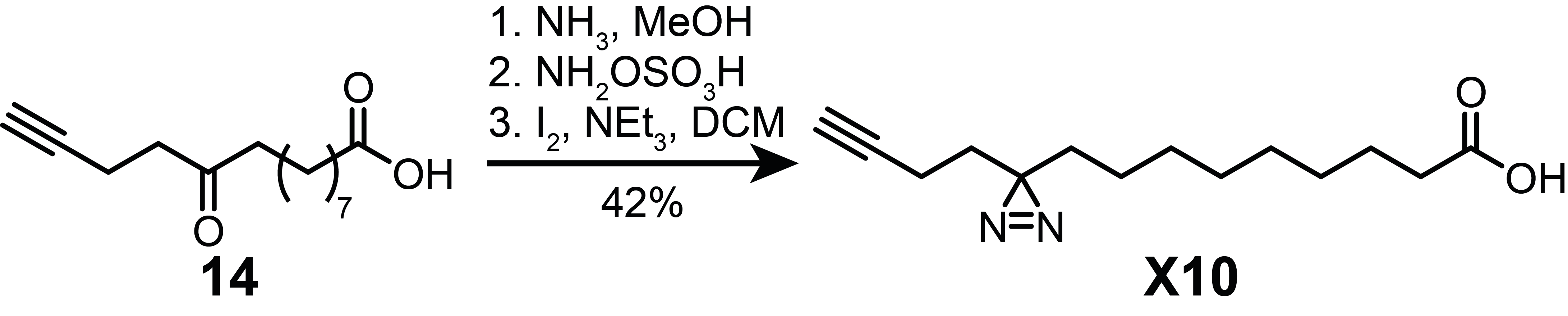

**Preparation of 9-(3-(but-3-yn-1-yl)-3H-diazirin-3-yl)nonanoic acid (X10)**. Diazirine **X10** was synthesized following the same procedure as described for diazirine **X3** (see above) starting with acid **14** (150 mg, 0.63 mmol). The crude residue was purified using flash chromatography eluting with 15:85:1 EtOAc:Hexane:AcOH mixture, yielding diazirine **X10** as a white powder (67 mg, 42%). ^1^H NMR (400 MHz, CDCl_3_): δ 2.34 (t, *J* = 7.5 Hz, 2H), 2.02-1.96 (m, 3H), 1.65-1.58 (m, 4H), 1.44-1.38 (m, 2H), 1.36-1.19 (m, 8H), 1.13-1.02 (m, 2H). ^13^C NMR (101 MHz, CDCl_3_): δ 179.73, 83.03, 69.13, 34.06, 32.75, 32.55, 29.27, 29.20, 29.18, 29.09, 28.39, 24.75, 23.89, 13.48. HRMS (ESI^-^, m/z): calcd. for C_14_H_21_N_2_O_2_ [M-H]^-^ 249.1603; found 249.1593.

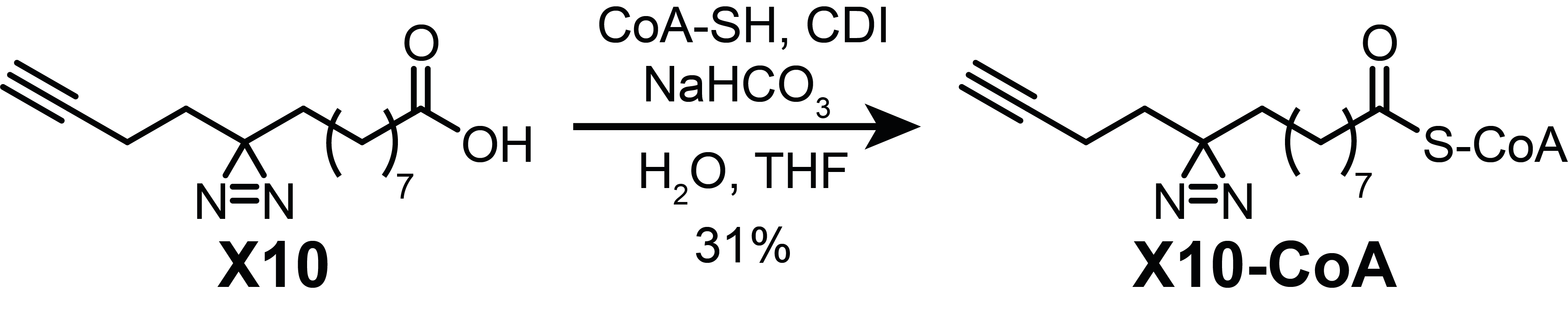

**Preparation of X10-CoA conjugate. X10-CoA** conjugate was synthesized using the procedure described in [8]. Diazirine **X10** (16 mg, 0.064 mmol) was dissolved in dry THF (1.0 mL) under nitrogen. Solution of CDI (10 mg, 0.064 mmol) in DCM (0.5 mL) was added and reaction mixture was stirred at RT for 30 min. The mixture was concentrated *in vacuo* and resuspended in dry THF (2.0 mL). CoA-SH (25 mg, 0.032 mmol) was dissolved in NaHCO_3_ _(aq.)_ (0.5 M, 2.0 mL) and added to the reaction mixture. The reaction was stirred for 4 h at RT under argon. THF was then removed *in vacuo* and the remaining aqueous phase was purified by preparative RP-HPLC. For preparative RP-HPLC a gradient of MeCN in 10 mM NH_4_OAc pH 5.2 (0–5 min 0% MeCN, 5–10 min linear gradient 0-50% MeCN, 10–15 min 50% MeCN) was used. The collected fractions were combined and concentrated *in vacuo* to give **X10-CoA** (21 mg, 31%) as a white solid. HRMS (ESI^-^, m/z): calcd. for C_35_H_55_N_9_O_17_SP_3_ [M-H]^-^ 998.2650; found 998.2640.

### Biological methods and proteomics

#### Reagents

All reagents were purchased from commercial sources and used as supplied. Human embryonic kidney 293T cells (HEK293T) and HeLa cells were obtained from ATCC and verified by STR by The Francis Crick Cell Services. MyrCoA, YnMyr, HsNMT1, HsNMT2, Hs pp60src(2–9) N-terminal peptide H‑GSNKSKPK‑NH_2_ were available from the laboratory and were described previously [9], [10]. Click reagents AzTB (azido-TAMRA-biotin used for enrichment on the streptavidin beads followed by in-gel fluorescence analysis or Western blotting), AzT (azido-TAMRA used for in-gel fluorescence analysis) and AzRB (azido-arginine-biotin used for enrichment on neutravidin beads and proteomics analysis) were available from the laboratory, their syntheses have been described previously [11]. Ultrapure water was obtained on a MilliQ Millipore water purification system.

#### Antibodies

DDX46 (Atlas antibodies, HPA036554; rabbit, 1:1000 dilution), FSP1 (Proteintech, 20886-1-AP, rabbit, 1:1000 dilution), GAPDH (Proteintech, 60004-1-Ig; mouse, 1:1000 dilution), NMT1 (Atlas antibodies, HPA022963; rabbit, 1:500 dilution), PSMC1 (Atlas antibodies, HPA000872; rabbit, 1:500 dilution), TOMM40 (Proteintech, 18409-1-AP; rabbit, 1:1000 dilution), Tra2β (Abcam, ab31353; rabbit, 1:1000 dilution), tubulin (Merck, T5168; mouse, 1:5000 dilution).

#### Recombinant HsNMT1 and crystallization

HsNMT1 was produced as a 6-His-tagged protein as described [9] and was available in the laboratory.

HsNMT1 was crystallized in complex with **X10-CoA** as previously described for HsNMT1:**Myr-CoA** complex using the sitting-drop vapor diffusion method at 20˚C [12]. Crystals of HsNMT1 in complex with **X10-CoA** we obtained in 2 µL droplets formed by mixing equal volumes of a protein solution at 7.05 mg/mL (supplemented with 0.5 mM of **X10-CoA** and 5% DMSO) and a cryosolution containing the precipitant components (16-20% (v/w) PEG 4K, 5 mM Ni(II) chloride, 100 mM sodium citrate pH 4.5, and 2.5% (v/v) glycerol). Crystals were then flashed cooled in liquid nitrogen and stored for data collection. A complete X-ray data set at 2.37 Å resolution was collected at 100*K* and using synchrotron radiation at Diamond Light Source (in Oxford, UK). The data set was integrated with XDS [13], and scaled and reduced using AIMLESS [14] from CCP4 package. Crystals belong to the *P*2_1_2_1_2 space group and have unit cell dimensions *a*=79.97 Å, *b*=177.45 Å, *c*=58.24 Å. Structure resolution was accomplished using the molecular replacement method with the software PHASER[15] and the crystallographic structure of HsNMT1:**Myr-CoA** complex as input model (PDB entry 4C2Y[12]). The initial model was subjected to alternate cycles of model building using COOT [16] and refinement using REFMAC-5 [17] and PHENIX 1.10.1 [18]. The structure consists of two molecules of HsNMT1:**X10-CoA** complex per asymmetric unit (chains A and B), and thus it was refined using NCS restrictions. Overall, the resulting electron density allowed to modeled almost entirely both NMT1 molecules, with the exception of 9 N-terminal residues of both A and B chains (NMT residues 109-114 and 3 more residues of the N-terminal tag), residues 182-188, 308-319, 407-414 of chain A, and residues 312-318 of chain B, which were flexible and thus not visible on the map. Electron density maps of good quality were obtained for the **X10-CoA** in both molecules of complex (Figure S1B). The cif file for the refinement of the **X10-CoA** probe was generated with ELBOW [19] as part of the PHENIX package. Difference electron density maps were calculated in PHENIX. The geometry of the final models was validated using MOLPROBITY [20]. Figures were generated using PYMOL (DeLano Scientific LLC, <http://pymol.sourceforge.net/>). X-ray data collection and statistics are summarized in Table 1.

#### CPM assay

The assay was performed as previously described in [10]. Briefly, various concentrations of either **Myr‑CoA** or **X10‑CoA** were added to hNMT1 or hNMT2 (200 ng/mL), peptide substrate (16 μM Hs pp60^src^(2–9) N-terminal peptide H-Gly-Ser-Asn-Lys-Ser-Lys-Pro-Lys-NH2) and 7-diethylamino-3-(4′-maleimidylphenyl)-4-methylcoumarin (8 μM) on a 96-well plate, 4 replicates per condition. Accumulation of fluorescence signal at 470 nm was measured at one-minute intervals during the first 15 min of the reaction. Rates of the enzymatic reaction were calculated based on the plotted linear range of fluorescence signal over time using GraphPad Prizm 5. The pseudo first order dependence of enzymatic rates on substrate concentration were fitted using Michaelis-Menten equation.

#### Generation of plasmids

mCitrine-TST (twin-strep-tag), Src-TST and Src6Q-TST constructs in pCMV vector were a kind gift from A. Konitsiotis.

Other TST-tagged constructs were assembled using NEBuilder® HIFI DNA Assembly Master Mix (New England Biolabs). The vector was PCR amplified (primers FW: TGGAGCCATCCGCAGTTTG; REV: CATGGTGGCGAAGCTTGAG). Genes of interest were amplified from HEK293T cells cDNA or plasmids available in the laboratory using the following primers:

RP2 FW: gctcaagcttcgccaccatgGGCTGCTTCTTC;

RP2 REV: tcaaactgcggatggctccaTATTCCCATCTGTATATCAGCAAAG;

FAM49B FW: gctcaagcttcgccaccatgGGGAATCTTCTTAAAGTTTTG;

FAM49B REV: tcaaactgcggatggctccaTTGCAGCATGGATTTAATTTG;

DDX46 FW: GGTACTCAAGGGCACCAGTATTC;

DDX46 REV: CCACATACATCATAGACCAGCAC;

NOL3 FW: gctcaagcttcgccaccatgGGCAACGCGCAGGAG;

NOL3 REV tcaaactgcggatggctccaGGAATCTTCGGACTCGTCCCTTTC;

GNAI2 FW gctcaagcttcgccaccatgGGCTGCACCGTGAG;

GNAI2 REV tcaaactgcggatggctccaGAAGAGGCCGCAGTCC;

GNAI3 FW gctcaagcttcgccaccatgGGCTGCAC;

GNAI3 REV tcaaactgcggatggctccaATAAAGTCCACATTCCTTTAAGTTG;

FSP1 FW gctcaagcttcgccaccatgGGGTCCCAGGTCTC;

FSP1 REV tcaaactgcggatggctccaAGGTGGAGACTGCCTCATG;

NCS1 FW gctcaagcttcgccaccatgGGGAAATCCAACAGCAAG;

NCS1 REV tcaaactgcggatggctccaTACCAGCCCGTCGTAGAG;

BASP1 FW gctcaagcttcgccaccatgGGAGGCAAGCTCAG;

BASP REV tcaaactgcggatggctccaCTCTTTCACGGTTACGGTTTG;

ARF1 FW: gctcaagcttcgccaccatgGGGAACATCTTCGCCAAC;

ARF1 REV: tcaaactgcggatggctccaCTTCTGGTTCCGGAGCTG;

CHP1 FW: gctcaagcttcgccaccatgGGTTCTCGGGCCTC;

CHP1 REV: tcaaactgcggatggctccaGTGAAGAAATCGGATGCTCATTTTCTG;

GNAO1 FW: gctcaagcttcgccaccatgGGATGTAC;

GNAO1 REV: tcaaactgcggatggctccaGTACAAGCCGCAGC;

PSMC1 FW: gctcaagcttcgccaccatgGGTCAAAGTCAGAGTG;

PSMC1 REV: tcaaactgcggatggctccaGAGATACAGCCCCTCAG;

ABL1 FW: gtttagtgaaccgccaccatgGGGCAGCAGCCTG;

ABL1 short REV: tcaaactgcggatggctccaAAAGGCTTGGTGGATTTCAGCAAAG;

NEF FW: gctcaagcttcgccaccatgGGGGGCAAGTGGTC;

NEF REV: tcaaactgcggatggctccaGCAGTCTTTGTAGTACTCCGGATG;

GAGpol p17 FW: gctcaagcttcgccaccatgGGTGCGAGAGCGTC;

GAGpol p17 REV: tcaaactgcggatggctccaGTAATTTTGGCTGACCTGGCTGTTG;

ARL17B FW: gctcaagcttcgccaccatgGGAAACATTTTTGAAAAGC;

ARL17B REV: tcaaactgcggatggctccaCTTTACCACAGCTGATTTC;

G2A mutants of FSP1 and DDX46 were cloned using Q5® Site-Directed Mutagenesis Kit (New England Biolabs) using the following primers:

FSP1 FW: GCCACCATGGcGTCCCAGGTC;

FSP1 REV: GAAGCTTGAGCTCGAGATCTG;

DDX46 FW: GCCACCATGGccCGGGAGTCAC;

DDX46 REV: GAAGCTTGAGCTCGAGATC.

#### Cell culture

HeLa and HEK293T cells were grown in low-glucose Dulbecco’s Modified Eagle’s Medium (DMEM) (Merck), supplemented with 10% (v/v) fetal bovine serum (FBS, Gibco). All cells were maintained in a humidified incubator at 37°C and 5% CO_2._

#### Metabolic labelling

HeLa or HEK293T cells were seeded into 10 cm plates or 6-well plates and grown for 24 h. Next, cells were incubated with cell growth media containing probes **YnMyr** (5 μM), **X3** (100 μM), **X8** (100 μM) or **X10** (100 μM) for additional 24 h.

#### UV crosslinking

After incubation, cells were carefully washed (×3) with PBS, placed on ice, and irradiated (in PBS) at 365 nm for 5 min using a Spectrolinker XL-1000 (Fisher Scientific) equipped with six 365 nm UV lamps. Irradiated cells were immediately lysed according to the protocol below.

#### Cell lysis

The cells were lysed with 1 mL (10 cm plates) or 200 µL (6-well plates) of lysis buffer containing 1% (v/v) Triton X-100, 0.1% (w/v) SDS, supplemented with cOmplete™ EDTA-free protease inhibitor cocktail (1×, Roche) in PBS on ice for 15 min. The lysates were clarified by centrifugation at 4°C for 10 min (at 17000 × g) and the supernatant recovered. Protein concentration was determined using the DC™ Protein Assay (Bio-Rad) in a 96-well plate as per manufacturer’s instructions.

#### Protein denaturation

10% SDS was added to lysates to 1% final concentration. Lysates were then heated to 95˚C for 10 min.

#### Click chemistry (CuAAC ligation)

Click chemistry was performed as described in [12]. Briefly, 0.06 volumes of click master mix was prepared by combining:

- 0.01 volume of click reagent (10 mM of AzT, AzTB or AzRB in DMSO)
- 0.02 volumes of 50 mM CuSO_4_ in MilliQ water
- 0.02 volumes of 50 mM TCEP in MilliQ water
- 0.01 volume of 10 mM TBTA in DMSO.

0.94 volumes of lysates were diluted to 2.13 mg/mL with PBS lysis buffer (corresponding to a concentration of 2 mg/mL in the final mixture). 0.06 volumes of the click master mix were added to 0.94 volumes of protein solution and the mixture was stirred for 1 h at RT. The reaction was then quenched by addition of 0.02 volumes of 0.5 M EDTA. 2 volumes of MeOH followed by 0.5 volumes of CHCl_3_ and followed by 1 volume of MilliQ water were then added to precipitate proteins. The precipitate was pelleted by centrifugation at 4°C for 10 min (17000 × g). The pellet was then washed twice with cold MeOH by resuspension and centrifugation. Finally, the pellet was dissolved in 0.1 volume of 2% (w/v) SDS in PBS and diluted by 0.9 volumes of PBS.

#### SDS-PAGE and in-gel fluorescence

Lysates (10-20 ug) were mixed with 4× Laemmli sample loading buffer (250 mM Tris-HCl pH 6.8, 30% (v/v) glycerol, 10% (w/v) SDS, 0.05% (w/v) bromophenol blue) supplemented with 20% v/v β-mercaptoethanol (BME) and incubated at 95°C for 10 min. The samples were separated on 10% (w/v) SDS-polyacrylamide gels that were cast according to manufacturer's protocol (National Diagnostics). The gels were run for 15 min at 80 V, followed by 60 min at 160 V in Tris/glycine/SDS running buffer (0.192 M glycine, 25 mM Tris base, 1% (w/v) SDS). Fluorescent images of the gels were recorded on a Typhoon™ FLA 9500 (GE Healthcare; equipped with 532 nm excitation laser and an LPG emission filter). Molecular weights were estimated using Precision Plus Protein™ All Blue Prestained Protein Standard (Bio-Rad). For protein loading quantification, after the imaging, the gels were placed in a Coomassie staining buffer (10% (w/v) ammonium sulfate, 10% (v/v) phosphoric acid, 20% (v/v) methanol, 1.2% (w/v) Coomassie brilliant blue G) overnight. The gels were then washed in MilliQ water for 1 h and images were scanned using a Canon LiDE 120 office scanner.

#### Western blotting

SDS-PAGE gels were prepared as described above. Proteins were transferred to nitrocellulose membranes (GE Healthcare) in transfer buffer (192 mM glycine, 25 mM Tris base, 20% (v/v) MeOH) at 100 V for 1h and transfer was checked by staining with Ponceau S. Membranes were blocked with 5% (w/v) non-fat dried skimmed milk powder in TBS-T (50 mM Tris base (pH 7.4), 150 mM NaCl, 0.1% (v/v) Tween-20) or 3% (w/v) BSA in TBS-T for 1 h at RT. Membranes were then incubated with primary antibodies in blocking buffer for 1h at RT or overnight at 4°C. Membranes were washed 3 times with TBS-T, incubated with corresponding HRP-conjugated secondary antibodies in 5 % milk or 3 % BSA in TBS-T for 1 h at RT, washed again 3 times with TBS-T and visualized using HRP substrate (Luminata Crescendo, Millipore) on an ImageQuant™ LAS-4000 Imaging System.

#### Immunoprecipitation-Western blotting

C-terminally TST-tagged FSP1 and DDX46 as well as the corresponding G2A mutants were transfected with Lipofectamine 2000 in HEK293T cells and overexpressed for 48 h. Cells overexpressing FSP1 were optionally treated with 1 µM **IMP-1088** (NMTi) 24 h post-transfection for additional 24 h. The cells were subsequently lysed in a buffer containing 50 mM Tris pH 8.0, 150 mM NaCl and 1% NP-40 (IGEPAL CA-630, Merck). 15 µL of Streptactin®XT magnetic beads (IBA Lifesciences) were added per 600 µg of cell lysate and incubated at 4°C for 1 h. The beads with bound proteins were washed with 100 mM Tris pH 8.0, 150 mM NaCl and proteins were eluted by boiling in 2× Laemmli buffer. Fluorescence signal from secondary antibodies (LI-COR) was quantified using the Odyssey LI-COR system. Quantification was performed using Image Studio software (LI‑COR) and statistical analysis was performed using GraphPad Prism.

#### Streptavidin enrichment

15 μL of Dynabeads™ MyOne™ Streptavidin C1 (streptavidin beads) (Invitrogen) were washed three times with 100 μL 0.2% (w/v) SDS in PBS. Then 100 μL of 1 mg/mL “AzTB-clicked” lysates were added, and the suspension was stirred for 2 h at RT. The beads were washed three times with 1% (w/v) SDS in PBS and then resuspended in 15 μL of 1× Laemmli buffer. The tubes were placed in a heating block at 95°C for 10 min. The supernatants were then loaded onto 10% SDS-polyacrylamide gels.

#### Metabolic labeling and SILAC proteomics

Proteomics samples were prepared in LoBind microcentrifuge tubes (Eppendorf®). All buffer solutions were prepared fresh and filtered with a 0.22 μm syringe filter prior to use.

SILAC heavy spike-in mixture was prepared as follows. HeLa cells passaged in R10K8 SILAC heavy media (Dundee Cell Products) supplemented with 10% dialysed FBS (Merck) for at least 5 passages were treated in separate 10 cm tissue culture dishes with **YnMyr** (5 μM), **X3** (100 μM), **X8** (100 μM), **X10** (100 μM) and incubated for 24 h in a humidified incubator at 37°C and 5% CO_2_. Then, cells were lysed and protein concentration measured. Equal amounts of proteins from all four conditions were mixed to create a heavy-labeled spike-in.

HeLa cells in DMEM/10% FBS (light) media were treated in triplicates with **YnMyr** (5 μM), **X3** (100 μM), **X8** (100 μM), **X10** (100 μM) for 24 h with and without the addition of 0.5 μM of **IMP-366** (DDD85646) NMT inhibitor (NMTi). Cells were lysed and protein concentration measured as described above. 250 μg of heavy spike-in was added to 250 μg of each lysate. The proteins were ligated to AzRB regent using click chemistry as described above.

30 μL of NeutrAvidin® Agarose Resin slurry (Thermo Fisher Scientific) were washed (×3) with 500 μL of 0.2% (w/v) SDS in PBS by vortexing for 1 min and centrifugation in a microcentrifuge for 1 min. 1 mL of 1 mg/mL protein solution from AzRB click chemistry reaction was added to the beads and they were stirred at for 2 h at RT. The beads were washed (×3) with 500 μL 1% (w/v) SDS in PBS, followed by (×2) with 4 M urea in 50 mM NH_4_HCO_3_ (AMBIC), followed by (×3) with 50 mM (AMBIC). The beads were resuspended in 50 μL of 50 mM AMBIC and 5 μL of 100 mM TCEP and were stirred for 1 h at 37°C in a thermomixer (Eppendorf®). The beads were pelleted in a microcentrifuge and the supernatant was swapped for another 50 μL of 50 mM AMBIC and 5 μL of 100 mM iodoacetamide and left in the dark without stirring for 30 min. The beads were washed (×2) with 500 μL of 50 mM AMBIC. 50 μL of 50 mM AMBIC was again added to the beads followed by 1 μL of trypsin solution (20 μg Sequencing Grade Modified Trypsin from Promega vials were resuspended in 100 μL of 50 mM AMBIC) and left shaking in a thermomixer at 37°C overnight. The supernatant was collected, and the beads resuspended in 70 μL of 50 mM AMBIC and stirred for 15 min. The supernatant was collected and combined with a previous fraction. The beads were resuspended in 70 μL of 1.5% (v/v) TFA in MilliQ water and stirred for 15 min. The supernatant was collected again and combined with previous fractions. The peptides solution was desalted on a STAGE (stop-and-go extraction) tip. A stack of three Empore™ SDB XC polystyrene-divinylbenzene copolymer extraction disks (Merck) was cut and fit into a p200 pipette tip. The tips were activated with 150 μL of MeOH by centrifugation for 2 min at 2000 × g and washed with 150 μL of MilliQ water for 2 min at 2000 × g. The peptide samples were loaded onto the tips and centrifuged for 2 min at 2000 × g. The tips were washed with 150 μL of MilliQ water for 2 min at 2000 × g. The peptides were eluted from the tips using 60 μL of 79% MeCN in water by centrifugation for 2 min at 2000 × g. The samples were then dried in a Savant SPD1010 SpeedVac Concentrator (Thermo Scientific™) and stored at -80°C for subsequent proteomics analysis.

#### UV-crosslinking and SILAC proteomics

SILAC heavy spike-in mixture was prepared as follows. HeLa cells passaged in R10K8 SILAC heavy media (Dundee Cell Products) supplemented with 10% dialysed FBS (Merck) for at least 5 passages were treated in separate 10 cm tissue culture dishes with either **X3** (100 μM) or **X10** (100 μM), incubated for 24 h, UV crosslinked and lysed. HeLa cells in DMEM/10% FBS (light) media were treated in triplicates with either **X3** (100 μM) or **X10** (100 μM) for 24 h with or without the addition of 1 μM of **IMP‑366** (DDD85646). Cells were UV crosslinked, lysed and protein concentration measured. 200 μg of heavy spike-in was added to 400 μg of each lysate. The proteins were ligated to AzRB regent using click chemistry as described above. Enrichment and sample preparation was performed as described above for the metabolic labeling SILAC proteomics experiment**.**

#### Label-free interactome proteomics

HEK293 cells were plated into 6 cm tissue culture dishes and incubated overnight. Cells were transfected in triplicates with plasmids (1-4 mg) expressing a protein of interest using Lipofectamine LTX (3-12 μL) with PLUS reagent (Invitrogen) in Opti-MEM® (Gibco) according to the manufacturer’s protocol. After 6 h post-transfection, media was removed, replaced with fresh DMEM/10% FBS and cells were treated with 100 μM **X10.** After 24 h (30 h post-transfection) cells were washed (×3) with PBS, placed on ice and UV crosslinked. The cells were gently scraped, collected in a pellet and centrifuged for 3 min at 1000 × g.

60 μL of 1% (w/v) SDS in PBS was added to the pellets to initiate lysis. Lysates were then denaturated at 95˚C for 10 min to remove non-covalent interactions. After cooling to RT, 240 μL of 1.25% (v/v) Triton X-100 in PBS was added. DNA was broken down by repeatedly passing the lysates through 25G needles. The lysates were then centrifuged at 4˚C for 15 min (at 17000 × g). Supernatant was collected and protein concentration measured.

32 μL of StrepTactin beads (MagStrep "type3" XT from IBA Lifesciences) per sample were washed (×3) with 0.2% (w/v) SDS, 1% (v/v) Triton X-100 in PBS. 400 μg of lysate proteins were added and incubated with the beads at 4˚C overnight. The beads were then washed (×3) with 500 μL 0.2% (w/v) SDS, 1% (v/v) Triton X-100 in PBS, followed by (×3) with 50 mM AMBIC. The beads were resuspended in 50 μL of 50 mM AMBIC containing 10 mM TCEP and were stirred for 1 h at 37°C in a thermomixer (Eppendorf®). The beads were pelleted in a microcentrifuge and the supernatant was swapped for 50 μL of 50 mM AMBIC and 5 μL of 10 mM iodoacetamide and left in the dark without stirring for 30 min. The beads were washed (×2) with 500 μL of 50 mM AMBIC. 50 μL of 50 mM AMBIC was again added to the beads followed by trypsin solution containing 0.11 μg of trypsin (20 μg Sequencing Grade Modified Trypsin from Promega vials were resuspended in 100 μL of 50 mM AMBIC) and left shaking in a thermomixer at 37°C for 4 h. The supernatant was collected, and the beads were resuspended in 70 μL of 50 mM AMBIC and stirred for 15 min. The supernatant was collected and combined with a previous fraction. The beads were resuspended in 70 μL of 1.5% (v/v) TFA in MilliQ water and stirred for 15 minutes. The supernatant was collected again and combined with previous fractions.

The peptides solution was desalted on a STAGE (stop-and-go extraction) tip. A stack of three Empore™ SDB XC polystyrene-divinylbenzene copolymer extraction disks (Merck) was cut and fit into a p200 pipette tip. The tips were activated with 150 μL of MeOH by centrifugation for 2 min at 2000 × g and washed with 150 μL of MilliQ water for 2 min at 2000 × g. The peptide samples were loaded onto the tips and centrifuged for 2 min at 2000 × g. The tips were washed with 150 μL of MilliQ water for 2 min at 2000 × g. The peptides were eluted from the tips using 60 μL of 79% MeCN in water by centrifugation for 2 min at 2000 × g. The samples were then dried in a Savant SPD1010 SpeedVac Concentrator (Thermo Scientific™) and stored at -80°C for subsequent proteomics analysis.

#### Shotgun proteomics

Dried peptide samples were dissolved in 15 µL of Optima™ LC/MS H_2_O (Fisher Scientific) containing 2% (v/v) UPLC grade MeCN and 0.5% (v/v) TFA, vortexed and sonicated. 3 µL of the solution were injected into the LC-MS/MS system. 3 µL of the peptides sample was injected and separated on an EASY-Spray™ Acclaim PepMap C18 column (50 cm × 75 μm inner diameter) (Thermo Fisher Scientific) using a 2-hour gradient method with solvents A (2% MeCN supplemented with 0.1% formic acid) and B (80% MeCN supplemented with 0.1% formic acid) at a flow rate of 250 nL/min. Liquid chromatography was coupled to a QExactive mass spectrometer via an easy-spray source (Thermo Fisher Scientific). The QExactive was operated in data-dependent mode with survey scans acquired at a resolution of 70,000 at m/z 200. Scans were acquired from 350 to 1800 m/z. Up to 10 of the most abundant isotope patterns with charge +2 or higher from the survey scan were selected with an isolation window of 3.0 m/z and fragmented by HCD with normalized collision energies of 25 W. The maximum ion injection times for the survey scan and the MS/MS scans (acquired with a resolution of 17,500 at m/z 200) were 250 and 80 ms, respectively; the ion target value for MS was set to 10^6^ and for MS/MS to 10^5^.

#### Proteomics data processing (MaxQuant, Perseus and PEAKS)

Proteomics results (*.RAW files) were processed using MaxQuant (version 1.5.5.1) or PEAKS 8.0 using latest releases of *Homo sapiens* proteome from uniprot.org.

In MaxQuant, digestion was set as Trypsin/P, oxidation (M) and acetylation (N-term) were set as variable modifications, carbamidomethylation (C) was set as fixed modification and other settings were as by default. For SILAC quantification in MaxQuant multiplicity was set to 2 and corresponding arginine and lysine labels were selected.

In PEAKS, digestion was set as Trypsin, precursor mass tolerance was set at 5.0 ppm, fragment ion mass tolerance was set 0.1 Da. Oxidation (M) and acetylation (N-term) were set as variable modifications, carbamidomethylation (C) was set as fixed modification. PEAKS PTM was used to find additional peptide modifications: Myr’ (ΔM=463.2907, at N-terminal glycines), Myr’-N2 (ΔM=489.2812, at N-terminal glycines) and Myr-Xlink (ΔM=479.2856, at any residue).

Perseus 1.5.5.3 was used to analyze files proteinGroups.txt obtained as result of MaxQuant searches. Potential contaminants, reverse and only identified by site proteins were discarded before the analysis. All data was transformed using log2(x) function before any further manipulation. Data was normalized by subtracting a median of each data column.

t-test was performed using default Perseus settings, volcano plot representations have a visual cut-off line with parameters: s0=0.1, FDR=0.05, unless stated otherwise.

GraphPad Prism 8 was used to visualize the data.

#### StringDB analysis of protein interactions

Proteins significantly enriched in the DDX46-TST transfected sample (DDX46, ZC3H13, PPIG, SRRM2, TRA2A, TRA2B, ACIN1, LMAN1) were used as input to StringDB. The graph was built using default parameters and with 5 additional interactors shown.
