## Supplementary material for "Discovery of lipid-mediated protein-protein interactions in living cells using metabolic labeling with photoactivatable clickable probes": NMR spectra

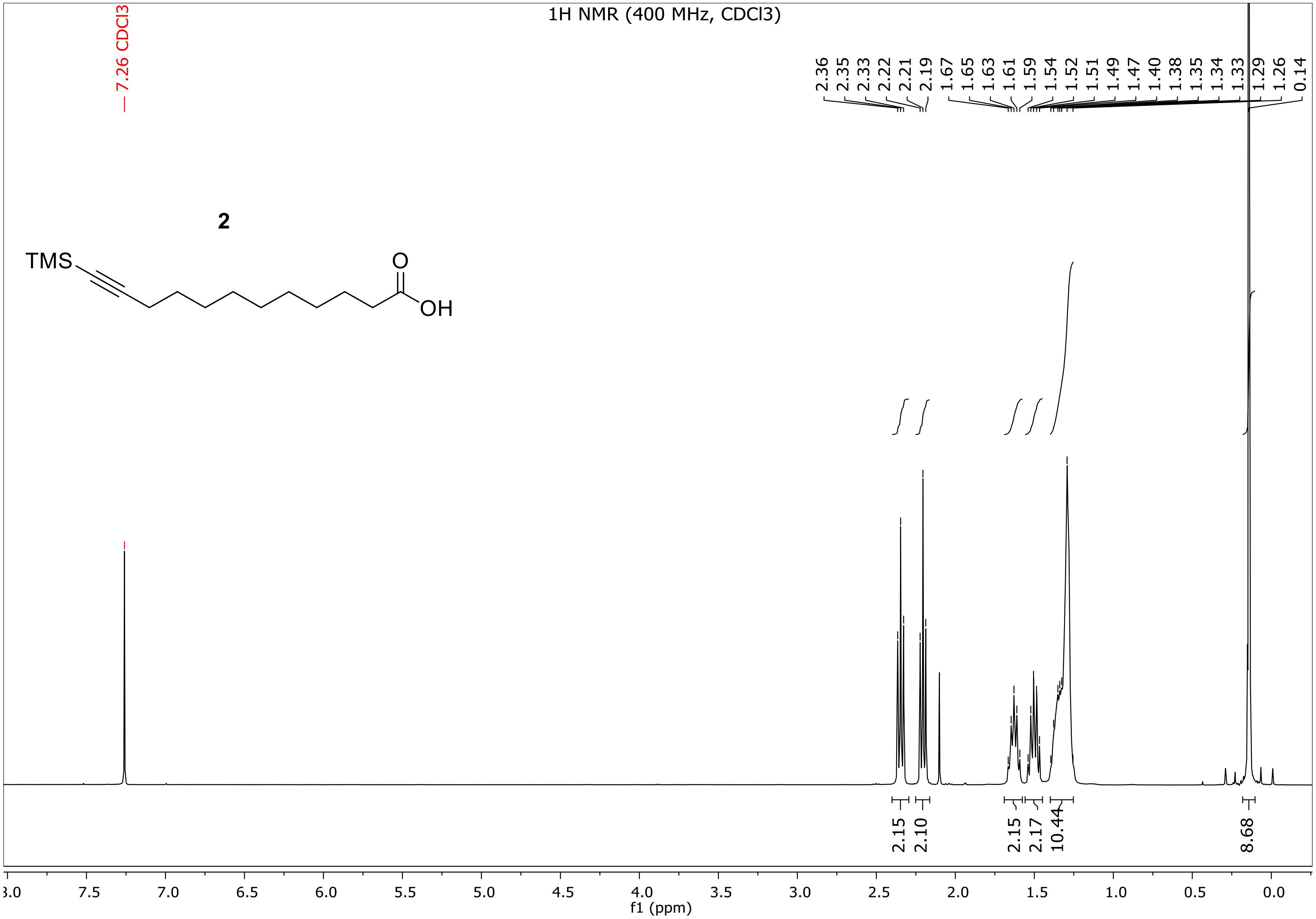

1H NMR (400 MHz, CDCl3)
7.26 CDCl3
2.36
2.35
2.33
2.22
2.21
2.19
1.67
1.65
1.63
1.61
1.59
1.54
1.52
1.51
1.49
1.47
1.40
1.38
1.35
1.34
1.33
1.29
1.26
0.14
2
TMS
O
OH
2.15
2.10
2.15
2.17
10.44
8.68
f1 (ppm)

13C NMR (101 MHz, CDCl3)

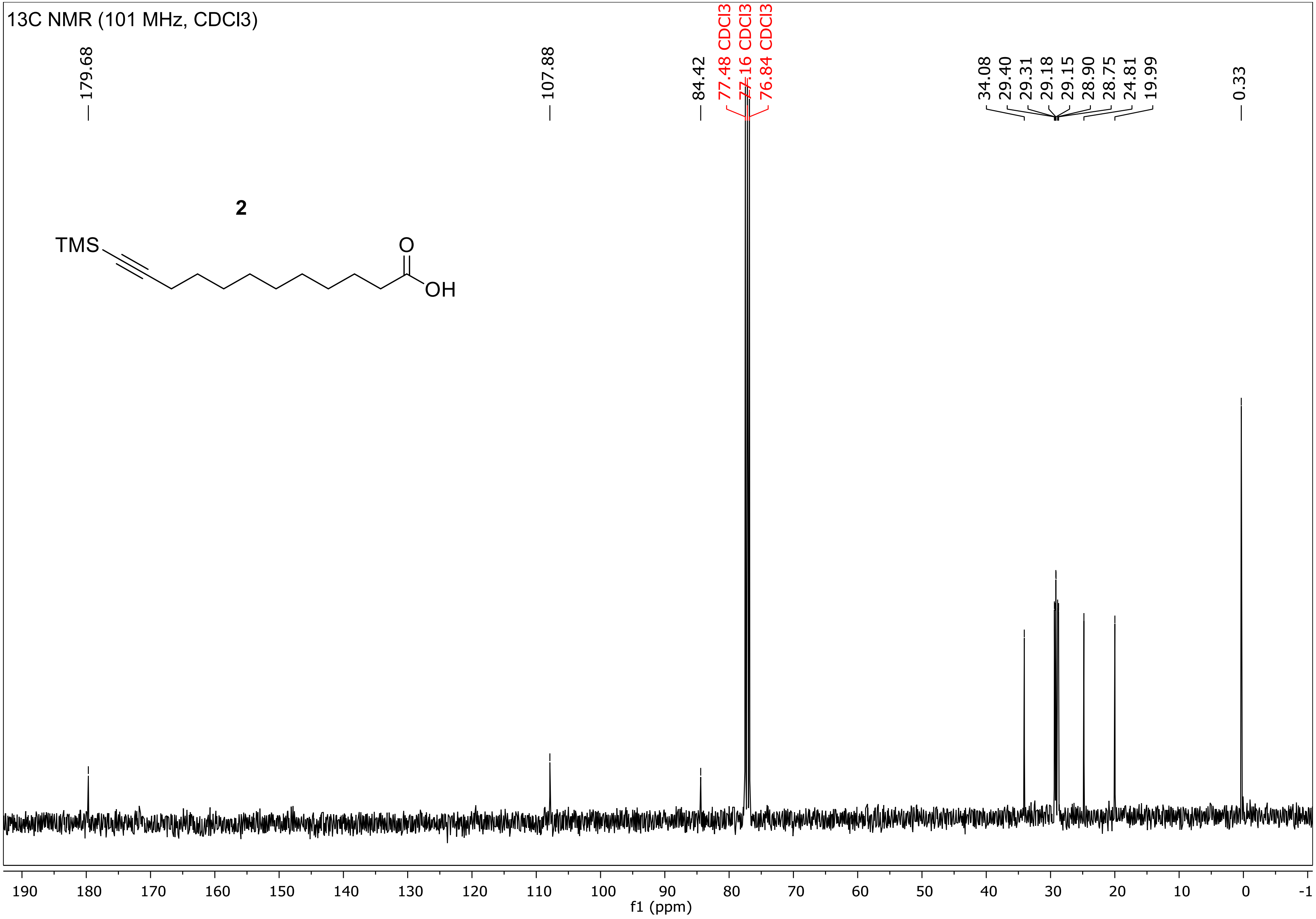

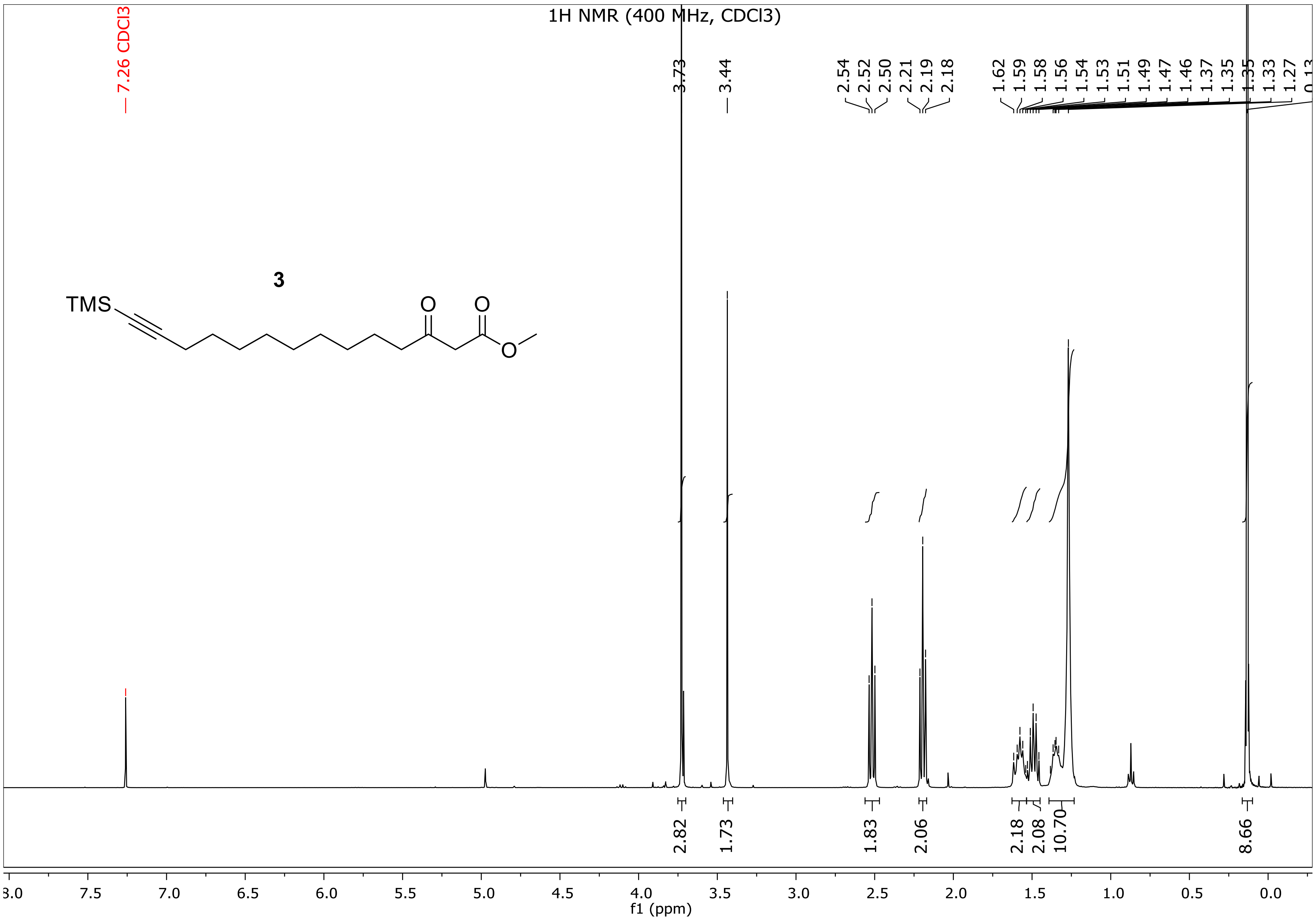

1H NMR (400 MHz, CDCl3)
7.26 CDCl3
3.73
3.44
2.54
2.52
2.50
2.21
2.19
2.18
1.62
1.59
1.58
1.56
1.54
1.53
1.51
1.49
1.47
1.46
1.37
1.35
1.35
1.33
1.27
3
TMS
O
O
O
2.82
1.73
1.83
2.06
2.18
2.08
10.70
8.66
7.5
7.0
6.5
6.0
5.5
5.0
4.5
4.0
3.5
3.0
2.5
2.0
1.5
1.0
0.5
0.0
f1 (ppm)

13C NMR (101 MHz, CDCl3)

3

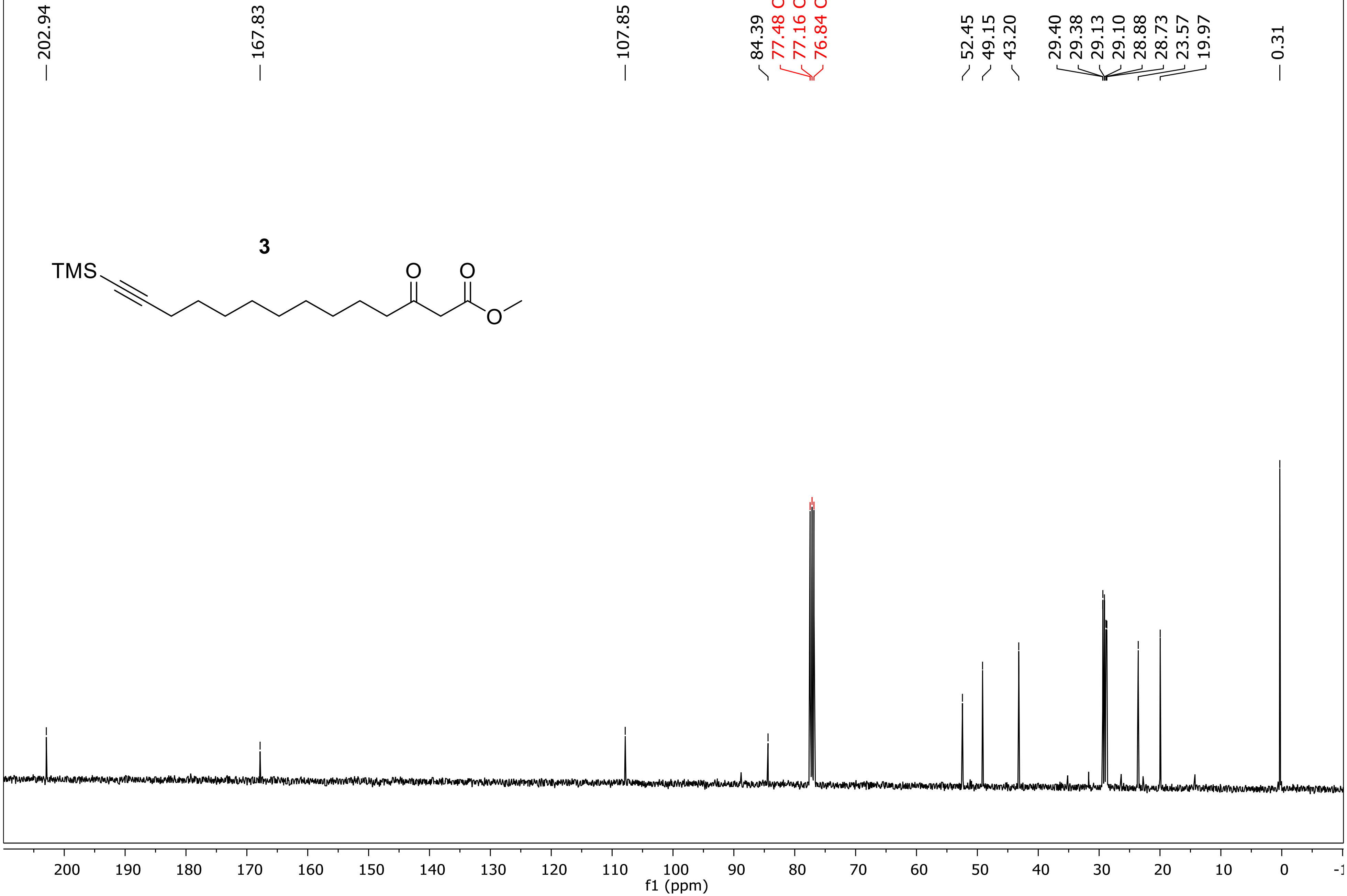

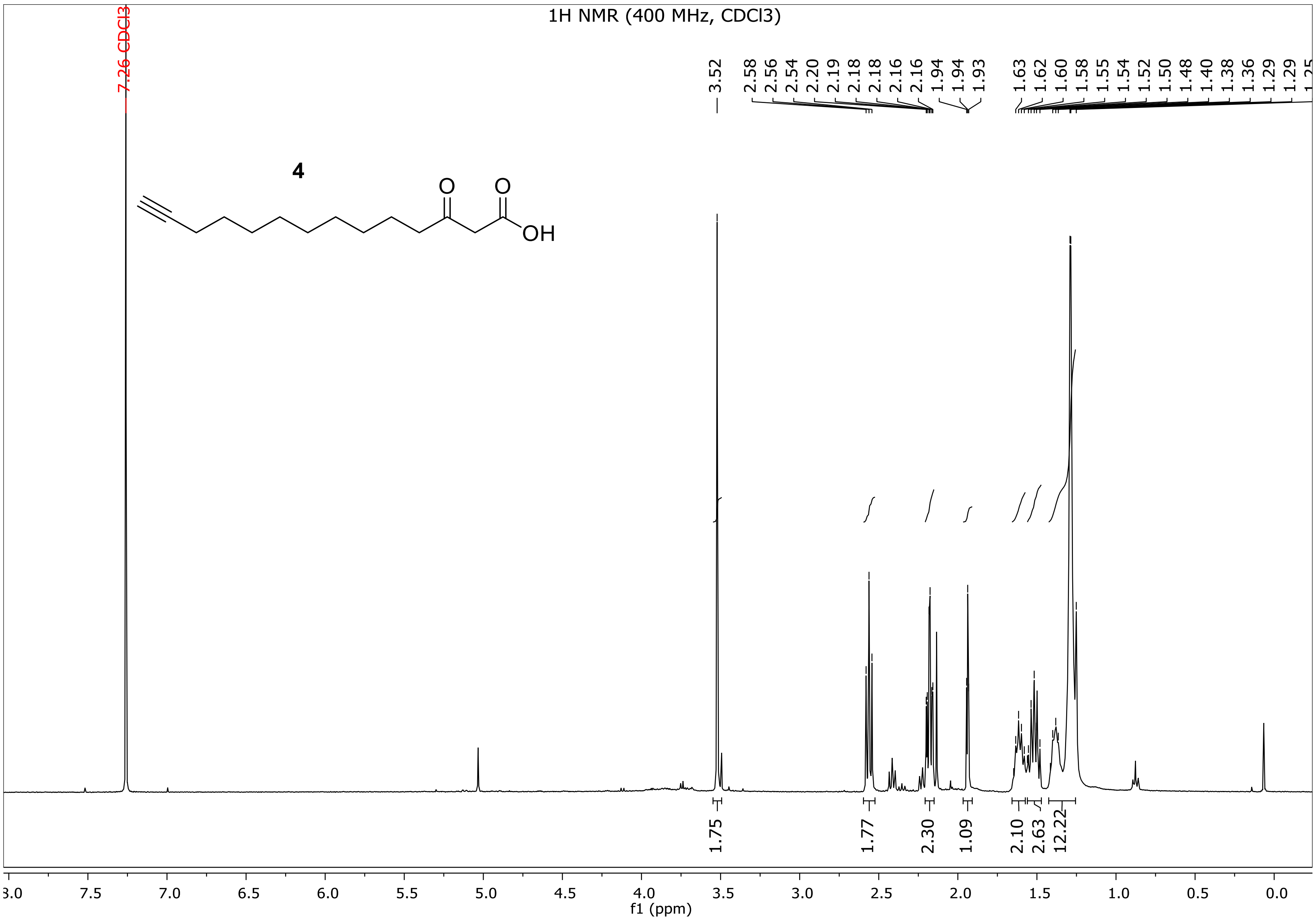
1H NMR (400 MHz, CDCl3)
7.26 CDCl3
3.52
2.58 2.56 2.54 2.20 2.19 2.18 2.18 2.16 2.16 1.94 1.94 1.93
1.63 1.62 1.60 1.58 1.55 1.54 1.52 1.50 1.48 1.40 1.38 1.36 1.29 1.29
4
O
O
OH
1.75
1.77
2.30
1.09
2.10
2.63
12.22
8.0 7.5 7.0 6.5 6.0 5.5 5.0 4.5 4.0 3.5 3.0 2.5 2.0 1.5 1.0 0.5 0.0
f1 (ppm)

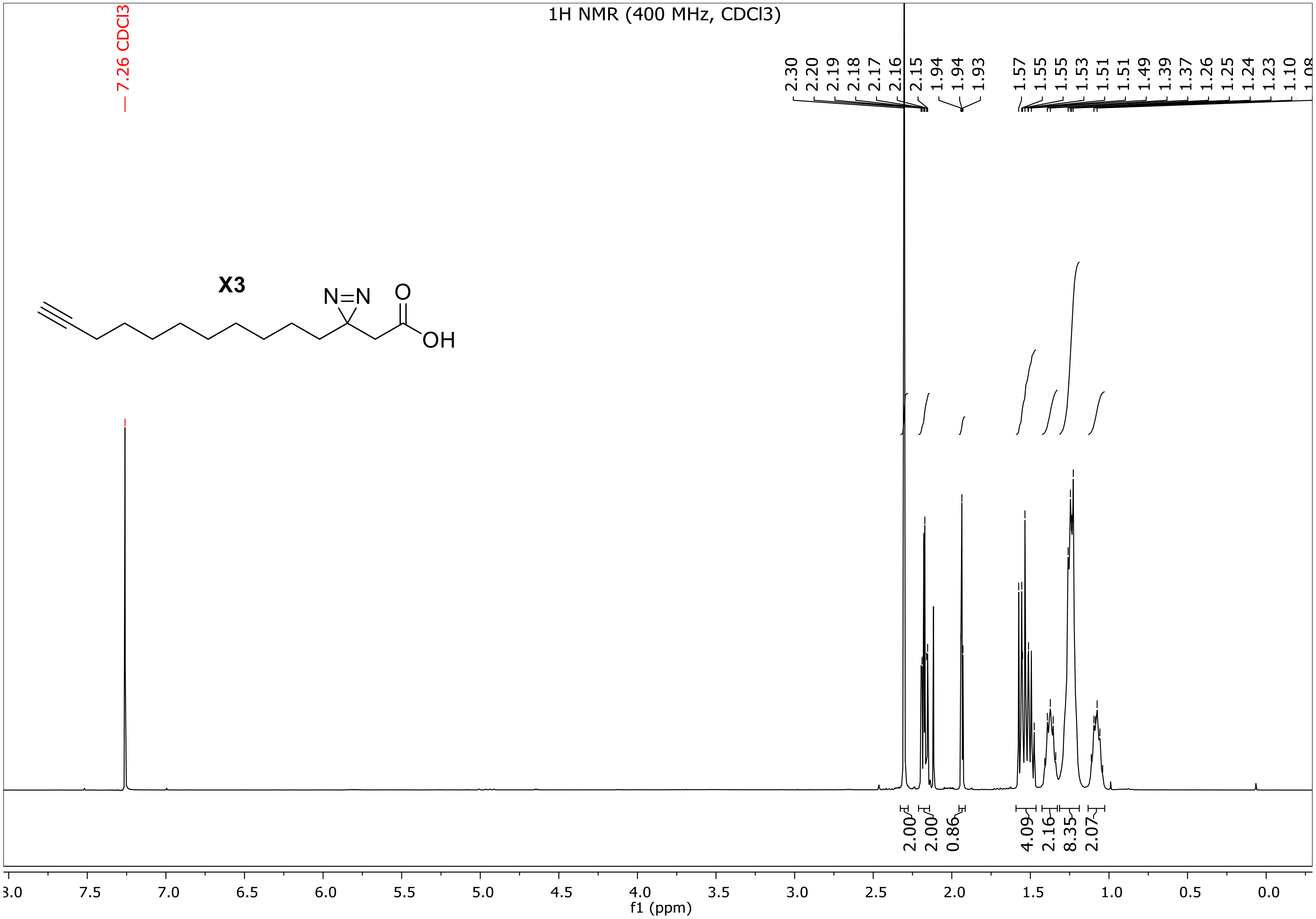

1H NMR (400 MHz, CDCl3)
7.26 CDCl3
2.30 2.20 2.19 2.18 2.17 2.16 2.15 1.94 1.94 1.93
1.57 1.55 1.55 1.53 1.51 1.51 1.49 1.39 1.37 1.26 1.25 1.24 1.23 1.10
X3
N=N
O
OH
2.00 2.00 0.86 4.09 2.16 8.35 2.07
7.5 7.0 6.5 6.0 5.5 5.0 4.5 4.0 3.5 3.0 2.5 2.0 1.5 1.0 0.5 0.0
f1 (ppm)

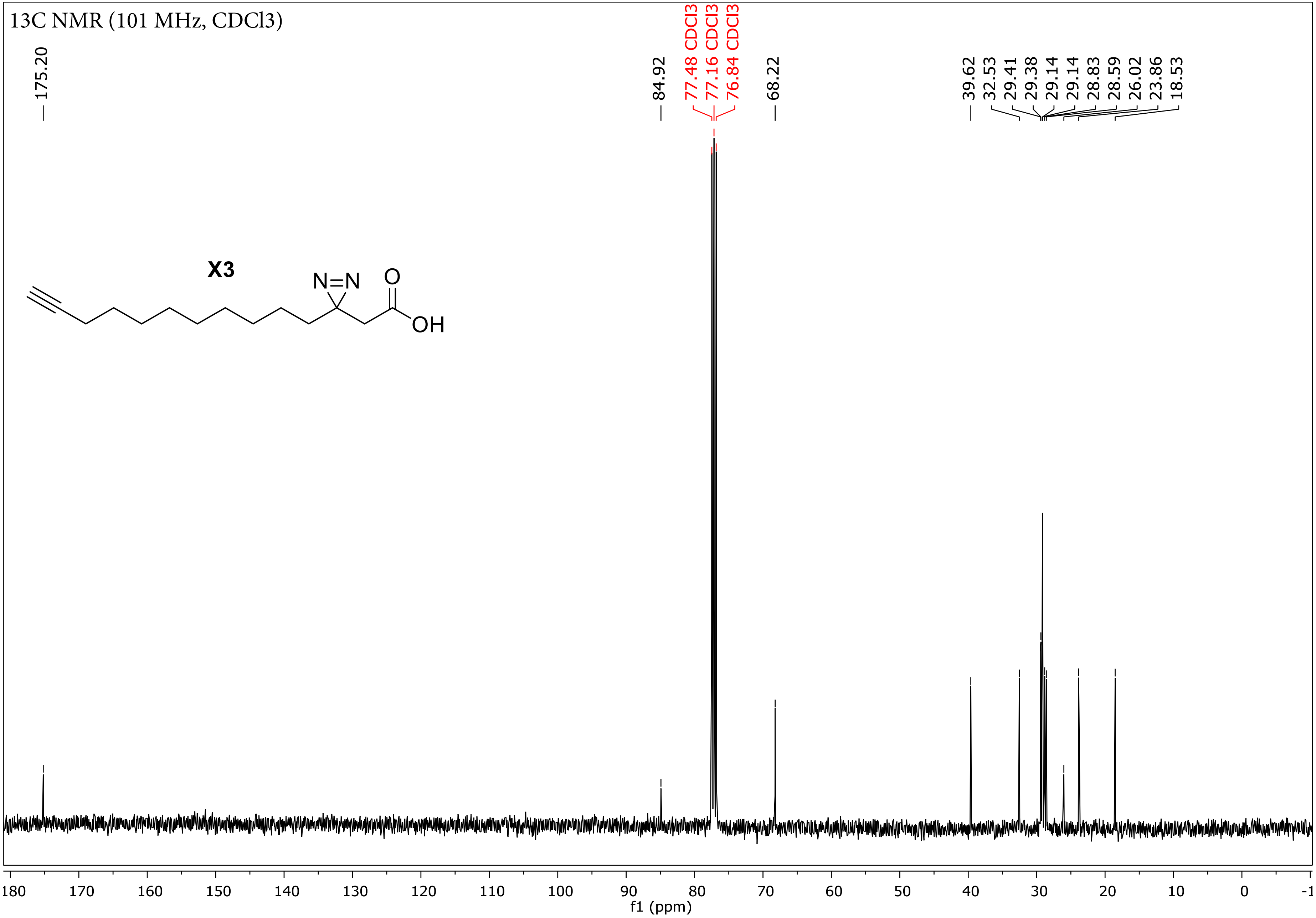

13C NMR (101 MHz, CDCl3)
175.20
84.92
77.48 CDCl3
77.16 CDCl3
76.84 CDCl3
68.22
39.62
32.53
29.41
29.38
29.14
29.14
28.83
28.59
26.02
23.86
18.53
X3
N=N
O
OH
180 170 160 150 140 130 120 110 100 90 80 70 60 50 40 30 20 10 0 -1
f1 (ppm)

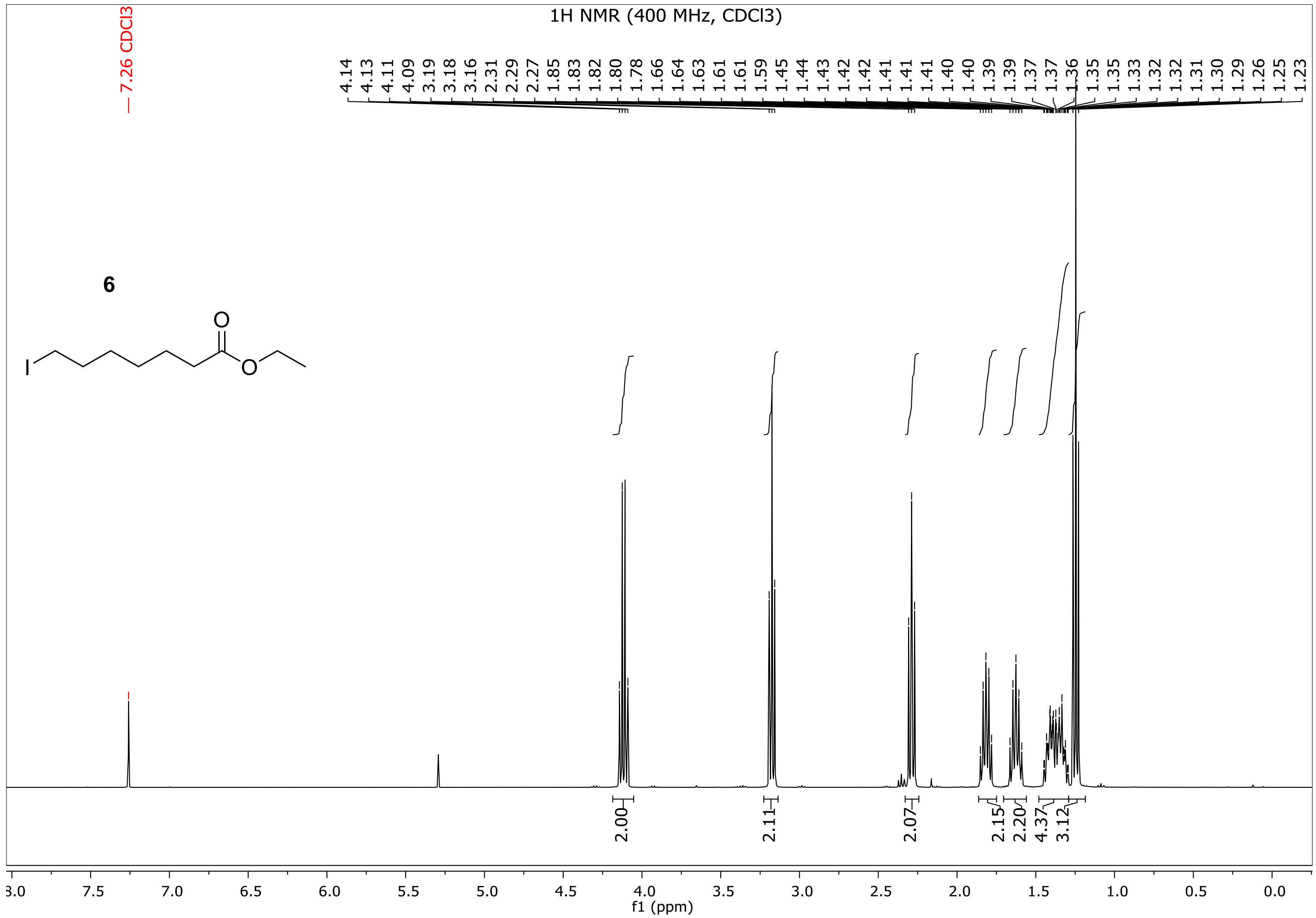
1H NMR (400 MHz, CDCl3)
7.26 CDCl3
4.14 4.13 4.11 4.09 3.19 3.18 3.16 2.31 2.29 2.27 1.85 1.83 1.82 1.80 1.78 1.66 1.64 1.63 1.61 1.61 1.59 1.45 1.44 1.43 1.42 1.42 1.41 1.41 1.41 1.40 1.40 1.39 1.39 1.37 1.37 1.36 1.35 1.35 1.33 1.32 1.32 1.31 1.30 1.29 1.26 1.25 1.23
6
I
O
O
2.00
2.11
2.07
2.15
2.20
4.37
3.12
8.0 7.5 7.0 6.5 6.0 5.5 5.0 4.5 4.0 3.5 3.0 2.5 2.0 1.5 1.0 0.5 0.0
f1 (ppm)

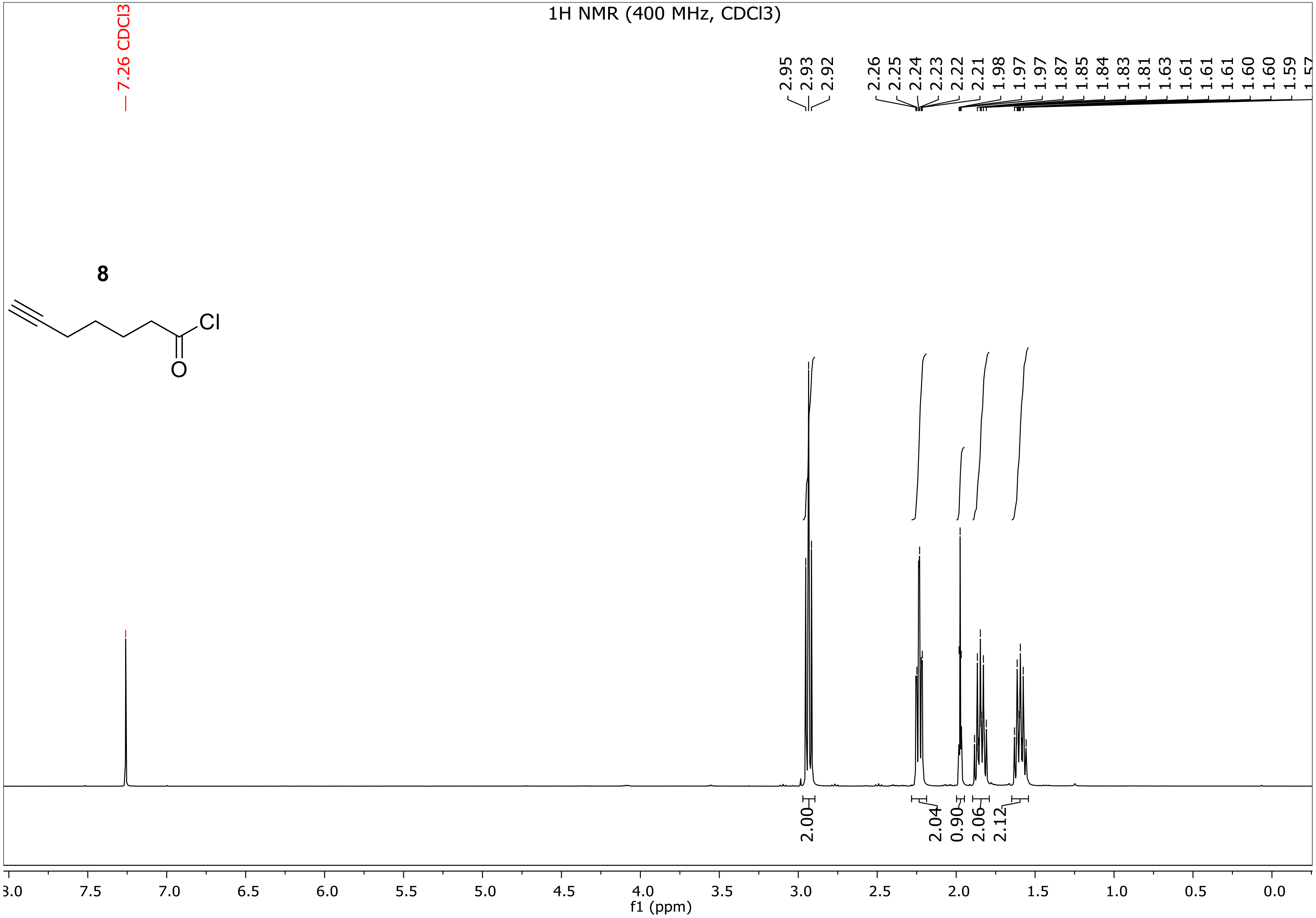

1H NMR (400 MHz, CDCl3)
7.26 CDCl3
2.95
2.93
2.92
2.26
2.25
2.24
2.23
2.22
2.21
1.98
1.97
1.97
1.87
1.85
1.84
1.83
1.81
1.63
1.61
1.61
1.61
1.60
1.60
1.59
8
Cl
O
2.00
2.04
0.90
2.06
2.12
7.5
7.0
6.5
6.0
5.5
5.0
4.5
4.0
3.5
3.0
2.5
2.0
1.5
1.0
0.5
0.0
f1 (ppm)

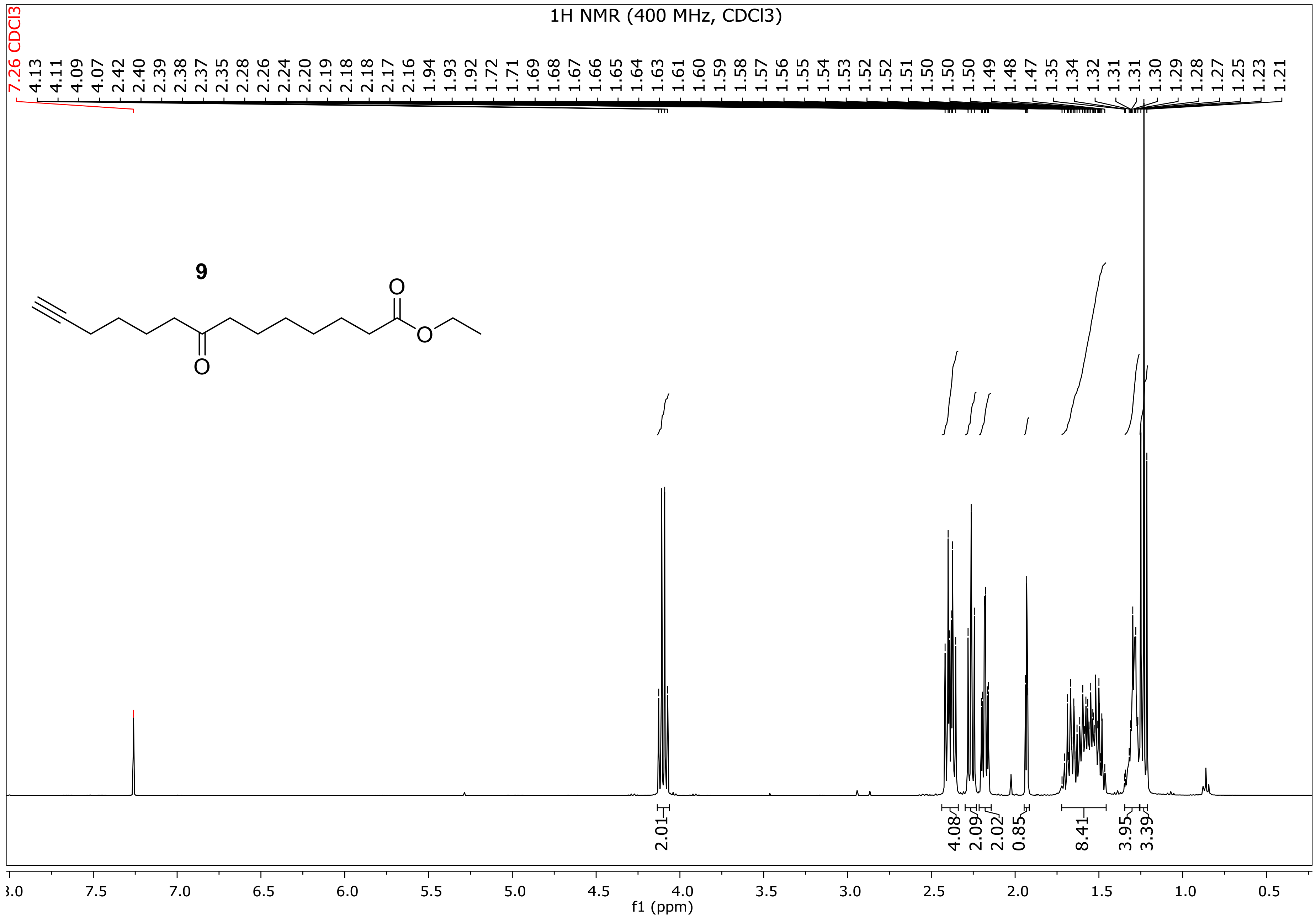
1H NMR (400 MHz, CDCl3)
7.26 CDCl3
4.13 4.11 4.09 4.07 2.42 2.40 2.39 2.38 2.37 2.35 2.28 2.26 2.24 2.20 2.19 2.18 2.18 2.17 2.16 1.94 1.93 1.92 1.72 1.71 1.69 1.68 1.67 1.66 1.65 1.64 1.63 1.61 1.60 1.59 1.58 1.57 1.56 1.55 1.54 1.53 1.52 1.52 1.51 1.50 1.50 1.50 1.49 1.48 1.47 1.35 1.34 1.32 1.31 1.31 1.30 1.29 1.28 1.27 1.25 1.23 1.21
9
O
O
O
2.01
4.08
2.09
2.02
0.85
8.41
3.95
3.39
8.0 7.5 7.0 6.5 6.0 5.5 5.0 4.5 4.0 3.5 3.0 2.5 2.0 1.5 1.0 0.5
f1 (ppm)

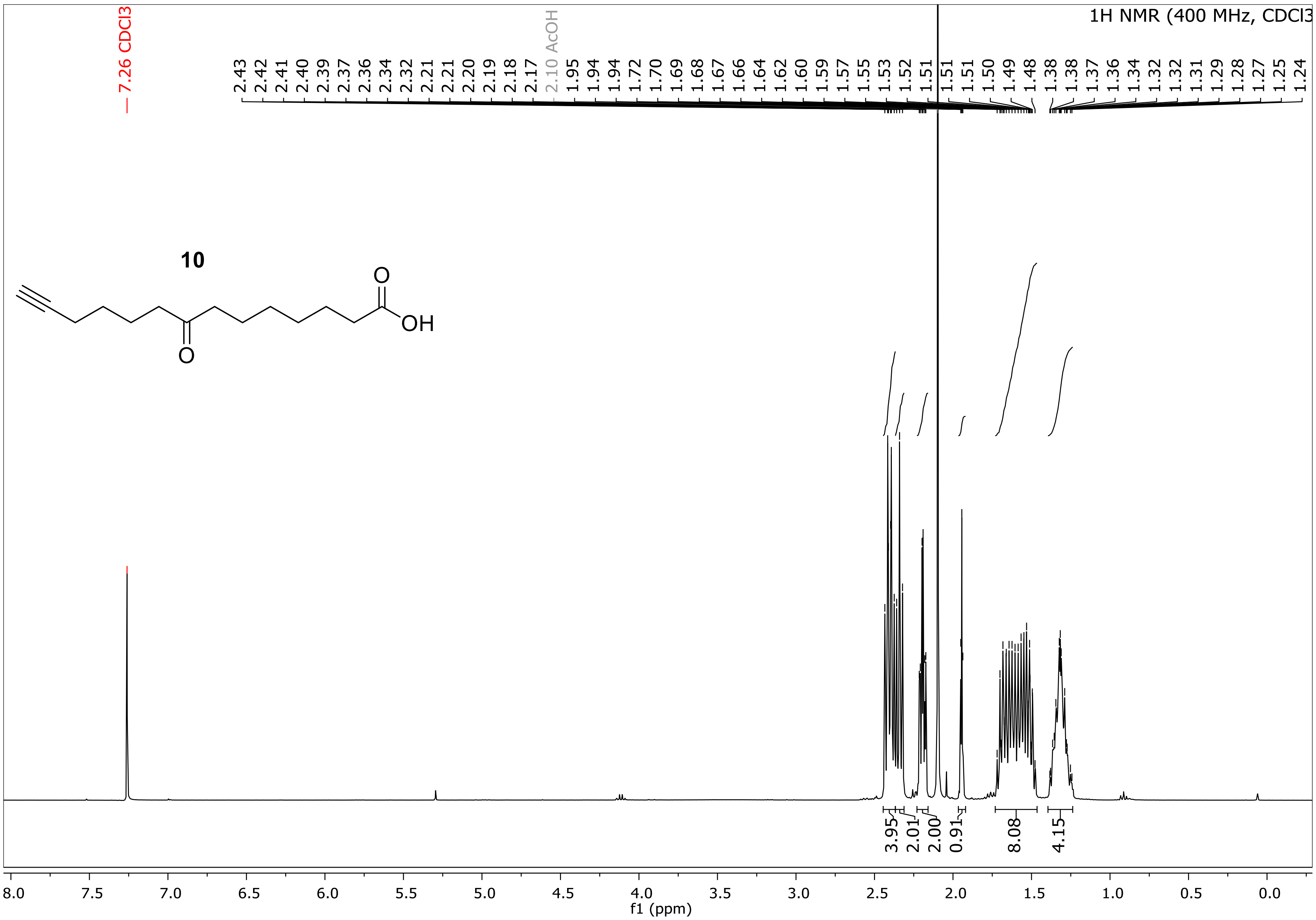

1H NMR (400 MHz, CDCl3
7.26 CDCl3
2.10 AcOH
2.43 2.42 2.41 2.40 2.39 2.37 2.36 2.34 2.32 2.21 2.21 2.20 2.19 2.18 2.17 1.95 1.94 1.94 1.72 1.70 1.69 1.68 1.67 1.66 1.64 1.62 1.60 1.59 1.57 1.55 1.53 1.52 1.51 1.51 1.51 1.50 1.49 1.48 1.38 1.38 1.37 1.36 1.34 1.32 1.32 1.31 1.29 1.28 1.27 1.25 1.24
10
O
OH
O
3.95 2.01 2.00 0.91 8.08 4.15
8.0 7.5 7.0 6.5 6.0 5.5 5.0 4.5 4.0 3.5 3.0 2.5 2.0 1.5 1.0 0.5 0.0
f1 (ppm)

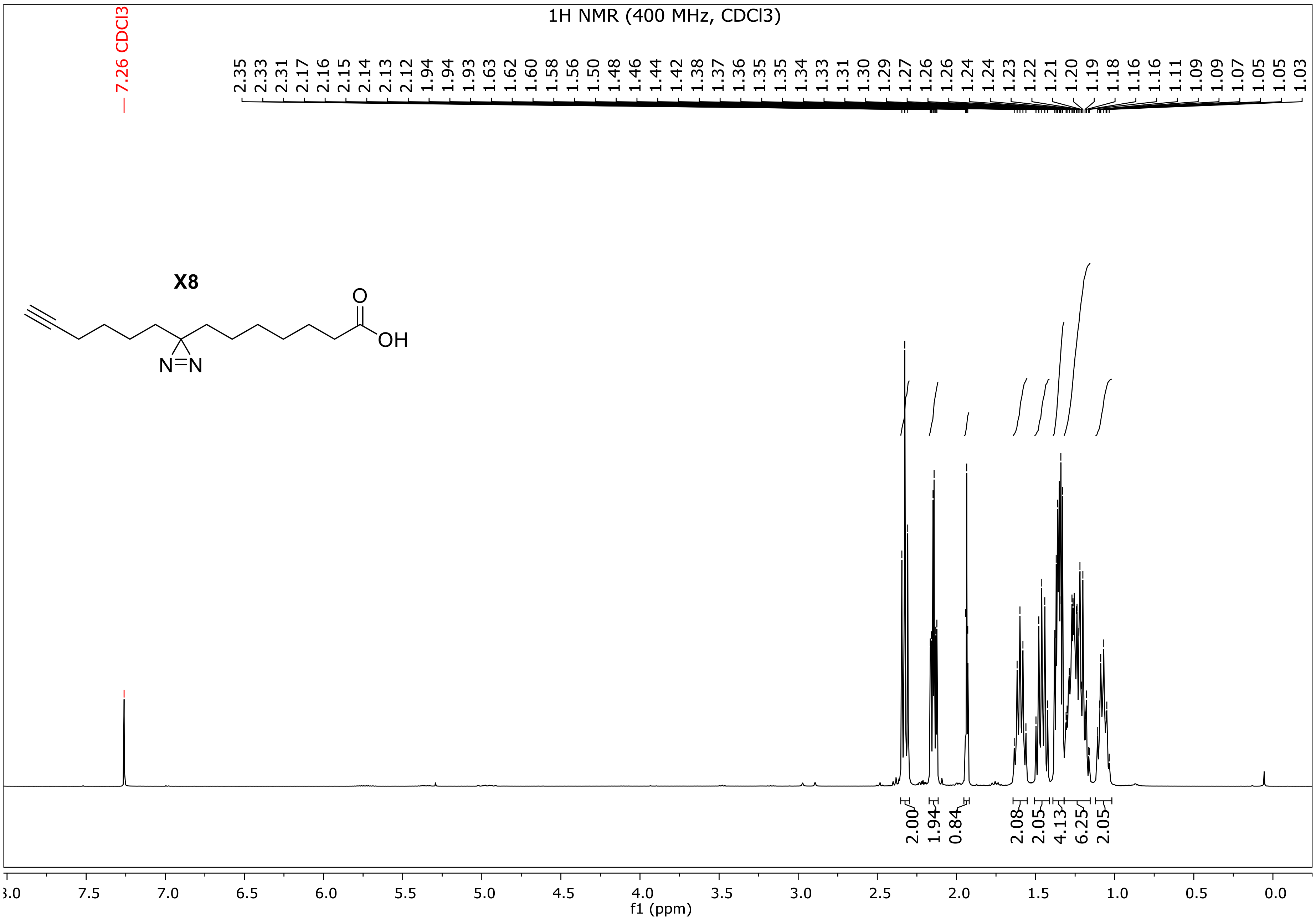
1H NMR (400 MHz, CDCl3)
7.26 CDCl3
2.35 2.33 2.31 2.17 2.16 2.15 2.14 2.13 2.12 1.94 1.94 1.93 1.63 1.62 1.60 1.58 1.56 1.50 1.48 1.46 1.44 1.42 1.38 1.37 1.36 1.35 1.35 1.34 1.33 1.31 1.30 1.29 1.27 1.26 1.26 1.24 1.24 1.23 1.22 1.21 1.20 1.19 1.18 1.16 1.16 1.11 1.09 1.09 1.07 1.05 1.05 1.03
X8
O
OH
N=N
2.00 1.94 0.84 2.08 2.05 4.13 6.25 2.05
8.0 7.5 7.0 6.5 6.0 5.5 5.0 4.5 4.0 3.5 3.0 2.5 2.0 1.5 1.0 0.5 0.0
f1 (ppm)

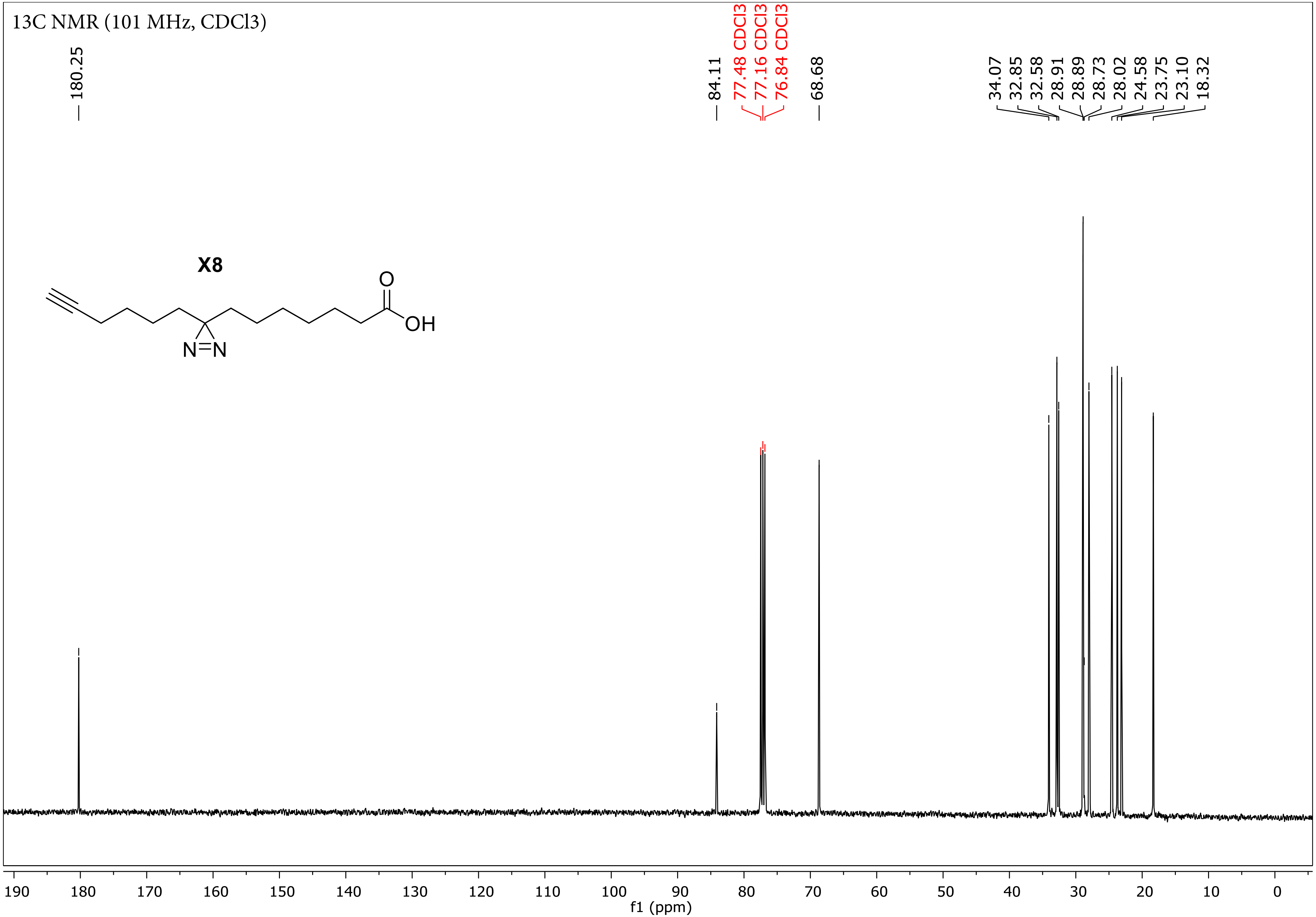
13C NMR (101 MHz, CDCl3)
180.25
84.11
77.48 CDCl3
77.16 CDCl3
76.84 CDCl3
68.68
34.07
32.85
32.58
28.91
28.89
28.73
28.02
24.58
23.75
23.10
18.32
X8
O
OH
N=N
190
180
170
160
150
140
130
120
110
100
90
80
70
60
50
40
30
20
10
0
f1 (ppm)

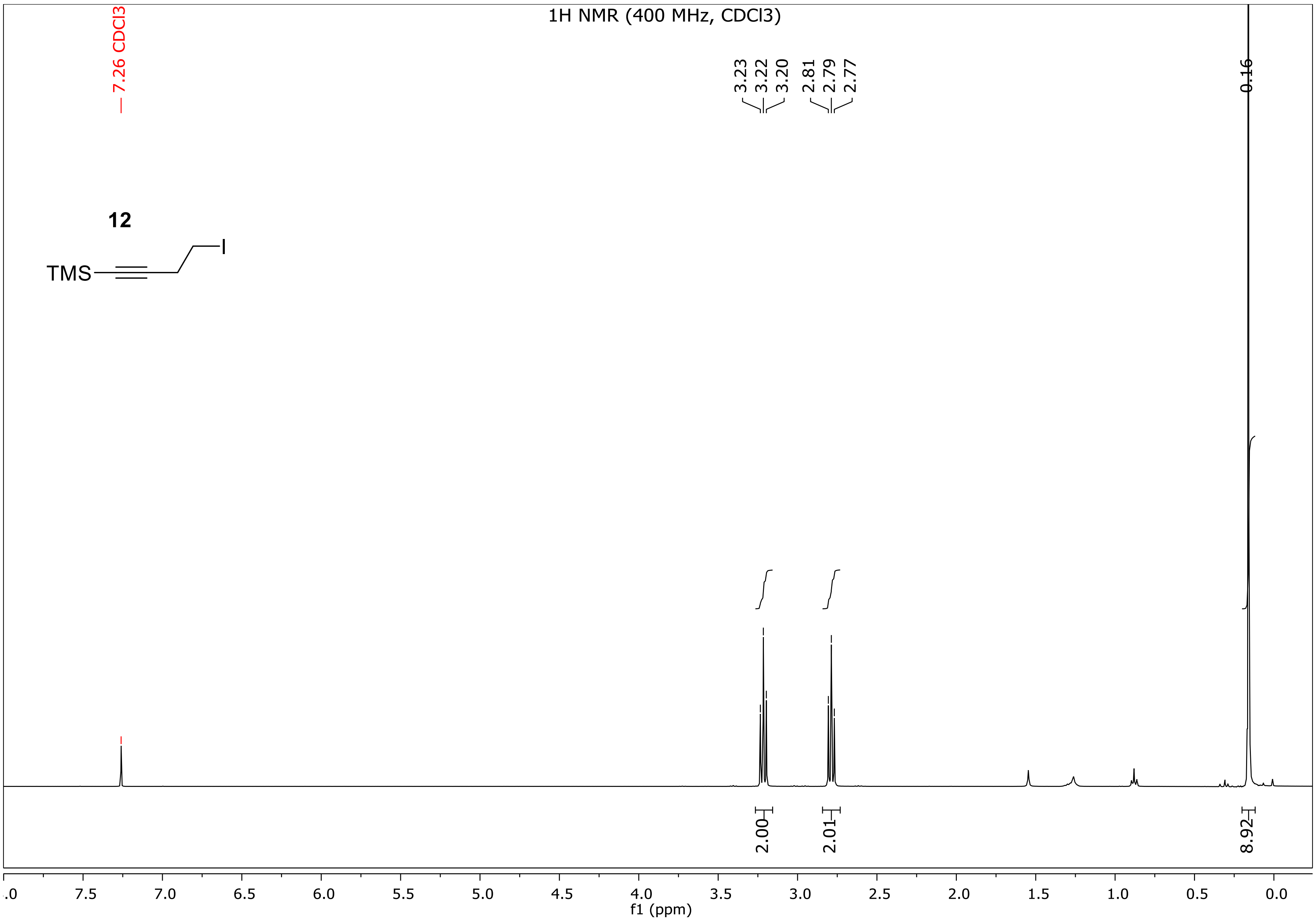
1H NMR (400 MHz, CDCl3)
7.26 CDCl3
12
TMS
I
3.23
3.22
3.20
2.81
2.79
2.77
0.16
2.00
2.01
8.92
f1 (ppm)

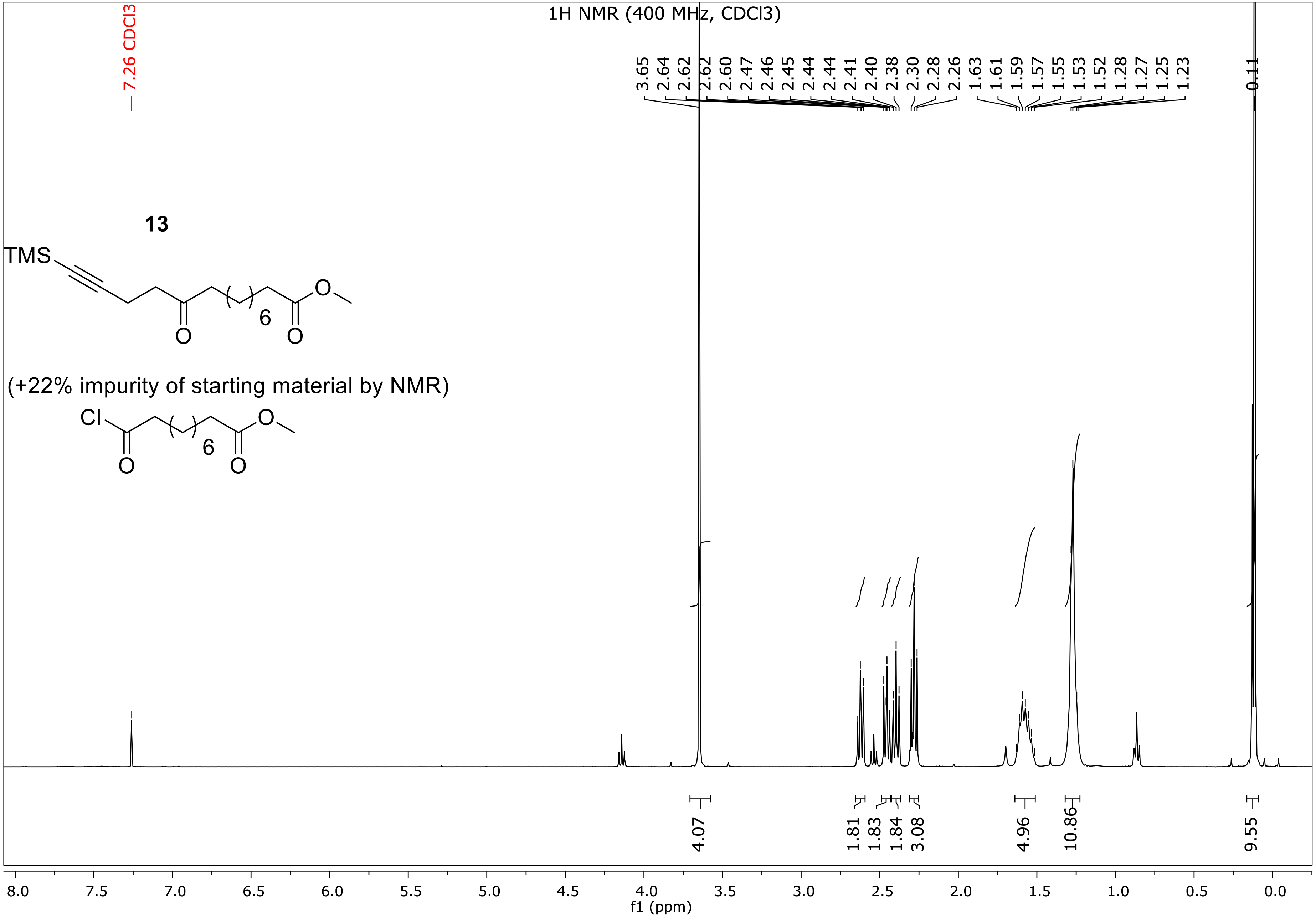
1H NMR (400 MHz, CDCl3)
7.26 CDCl3
3.65
2.64
2.62
2.62
2.60
2.47
2.46
2.45
2.44
2.44
2.41
2.40
2.38
2.30
2.28
2.26
1.63
1.61
1.59
1.57
1.55
1.53
1.52
1.28
1.27
1.25
1.23
0.11
13
TMS
6
O
O
O
(+22% impurity of starting material by NMR)
Cl
6
O
O
O
4.07
1.81
1.83
1.84
3.08
4.96
10.86
9.55
8.0
7.5
7.0
6.5
6.0
5.5
5.0
4.5
4.0
3.5
3.0
2.5
2.0
1.5
1.0
0.5
0.0
f1 (ppm)

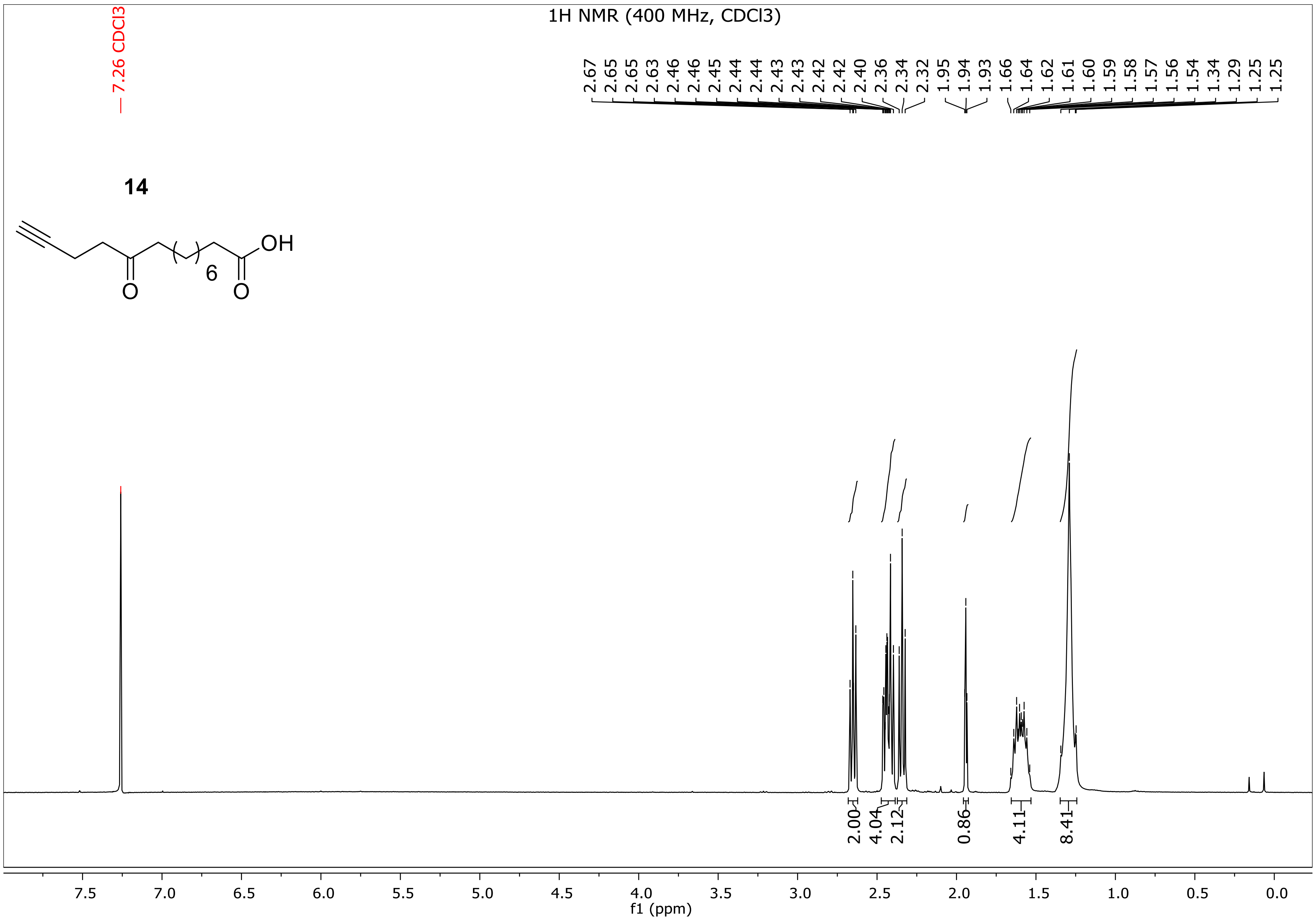

1H NMR (400 MHz, CDCl3)
7.26 CDCl3
2.67
2.65
2.65
2.63
2.46
2.46
2.45
2.44
2.44
2.44
2.43
2.43
2.42
2.42
2.40
2.36
2.34
2.32
1.95
1.94
1.93
1.66
1.64
1.62
1.61
1.60
1.59
1.58
1.57
1.56
1.54
1.34
1.29
1.25
1.25
14
OH
O
O
6
2.00
4.04
2.12
0.86
4.11
8.41
7.5
7.0
6.5
6.0
5.5
5.0
4.5
4.0
3.5
3.0
2.5
2.0
1.5
1.0
0.5
0.0
f1 (ppm)

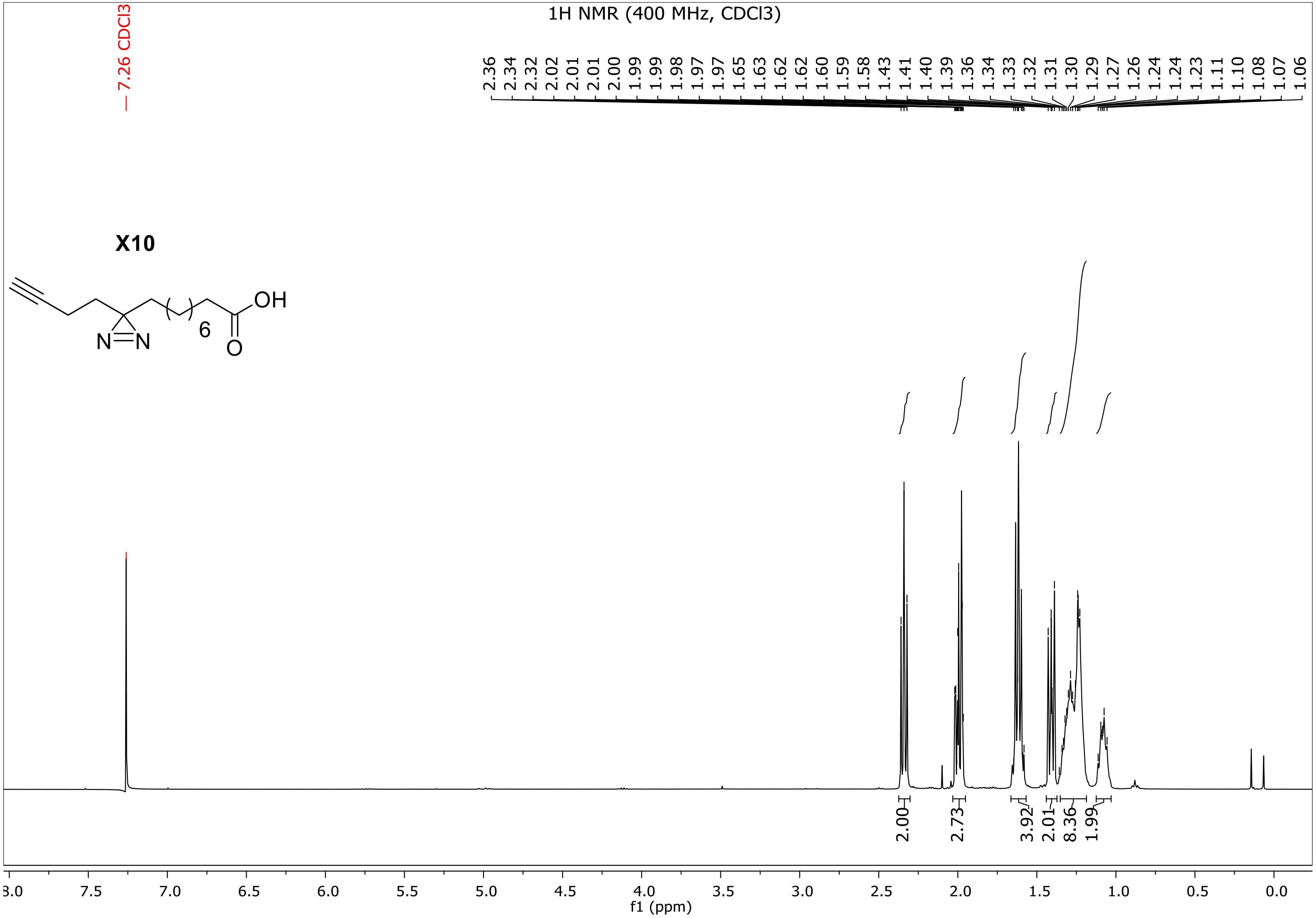
1H NMR (400 MHz, CDCl3)
7.26 CDCl3
2.36 2.34 2.32 2.02 2.01 2.01 2.00 1.99 1.99 1.98 1.97 1.97 1.65 1.63 1.62 1.62 1.60 1.59 1.58 1.43 1.41 1.40 1.39 1.36 1.34 1.33 1.32 1.31 1.30 1.29 1.27 1.26 1.24 1.24 1.23 1.11 1.10 1.08 1.07 1.06
X10
OH
6
N
N
O
2.00
2.73
3.92
2.01
8.36
1.99
8.0 7.5 7.0 6.5 6.0 5.5 5.0 4.5 4.0 3.5 3.0 2.5 2.0 1.5 1.0 0.5 0.0
f1 (ppm)

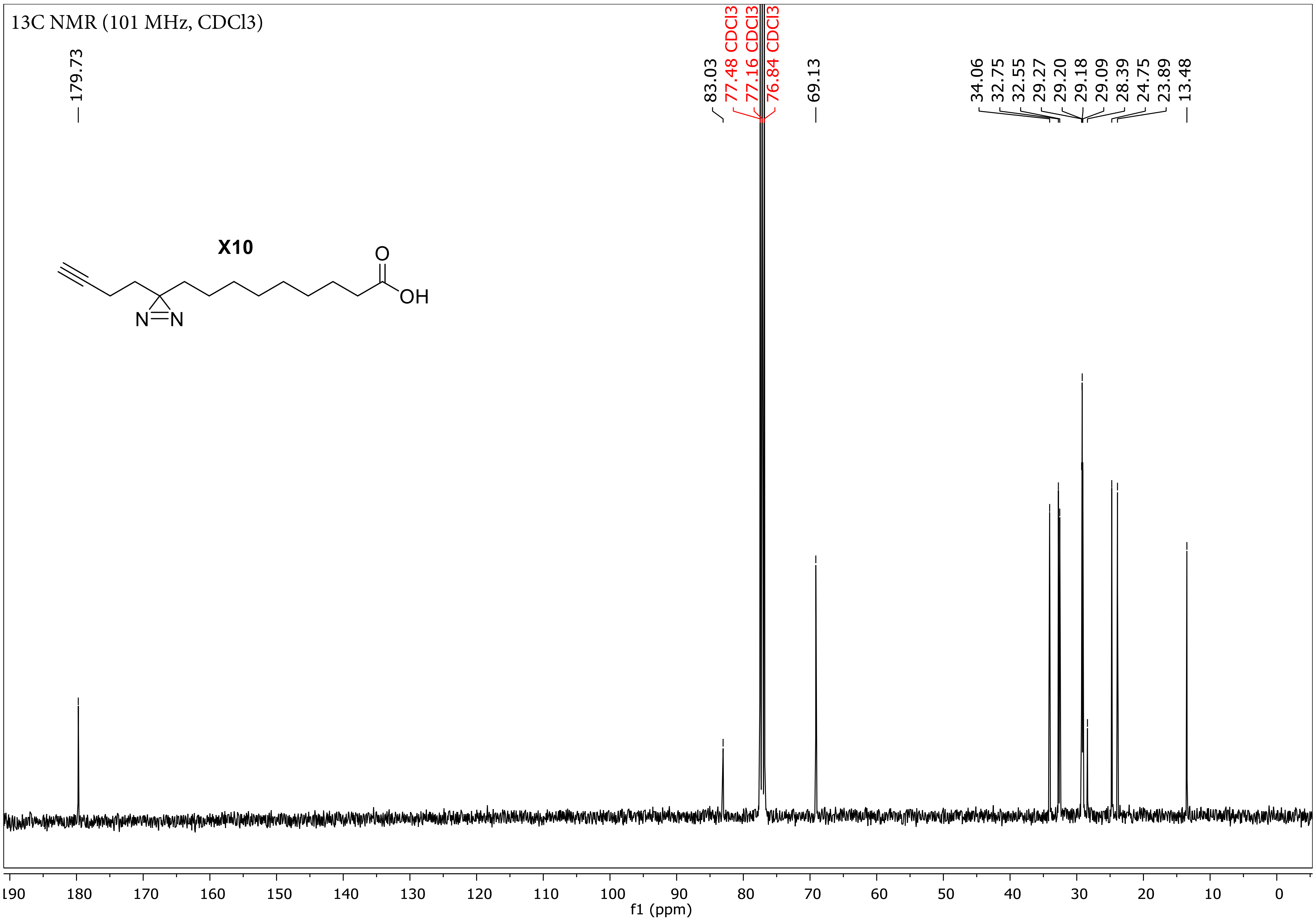
13C NMR (101 MHz, CDCl3)
179.73
83.03
77.48 CDCl3
77.16 CDCl3
76.84 CDCl3
69.13
34.06
32.75
32.55
29.27
29.20
29.18
29.09
28.39
24.75
23.89
13.48
X10
O
OH
N
N
190
180
170
160
150
140
130
120
110
100
90
80
70
60
50
40
30
20
10
0
f1 (ppm)
